# Evolutionary conservation of the structure and function of meiotic Rec114−Mei4 and Mer2 complexes

**DOI:** 10.1101/2022.12.16.520760

**Authors:** Dima Daccache, Emma De Jonge, Pascaline Liloku, Karen Mechleb, Marita Haddad, Sam Corthaut, Yann G.-J. Sterckx, Alexander N. Volkov, Corentin Claeys Bouuaert

## Abstract

Meiosis-specific Rec114−Mei4 and Mer2 complexes are thought to enable Spo11-mediated DNA double-strand-break (DSB) formation through a mechanism that involves DNA-dependent condensation. However, the structure, molecular properties, and evolutionary conservation of Rec114−Mei4 and Mer2 are unclear. Here, we present AlphaFold structures of Rec114−Mei4 and Mer2 complexes, supported by nuclear magnetic resonance (NMR) spectroscopy, small-angle X-ray scattering (SAXS), and mutagenesis. We show that dimers composed of the Rec114 C-terminus form α-helical chains that cup an N-terminal Mei4 α-helix, and that Mer2 forms a parallel homotetrameric coiled coil. Both Rec114−Mei4 and Mer2 bind preferentially to branched DNA substrates, indicative of multivalent protein-DNA interactions. Indeed, the Rec114−Mei4 interaction domain contains two DNA-binding sites that point in opposite directions and drive condensation. The Mer2 coiled-coil domain bridges co-aligned DNA duplexes, likely through extensive electrostatic interactions along the length of the coiled coil. Finally, we show that the structure of Rec114−Mei4 and Mer2 are conserved across eukaryotes, while DNA-binding properties vary significantly. This work provides insights into the mechanism whereby Rec114−Mei4 and Mer2 complexes promote the assembly of the meiotic DSB machinery, and suggests a model where Mer2 condensation is the essential driver of assembly, with the DNA-binding activity of Rec114−Mei4 playing a supportive role.

## Introduction

In most eukaryotes, the formation of haploid gametes requires Spo11-dependent catalysis of DNA double-strand breaks (DSBs) to initiate meiotic recombination, which is essential for the accurate segregation of homologous chromosomes (Hunter 2015). Spo11 activity relies on a higher-order assembly that is tied to chromosome structure and subject to overlapping regulatory pathways that control the timing, number and distribution of DSBs (Lam and Keeney 2015; Yadav and Claeys Bouuaert 2021). However, the molecular assemblies required for meiotic DSB formation are not well characterized, and the degree to which they are conserved is unclear.

From fungi to plants and animals, Spo11 activity depends on a cohort of accessory factors. While Spo11 itself is ubiquitous and well-conserved at the sequence level, the auxiliary proteins that constitute the DSB machinery vary more broadly between organisms, and functional homologs tend to be highly divergent (Keeney 2008; de Massy 2013; Lam and Keeney 2015). Nevertheless, some of the key partners are found throughout eukaryotes, including a sub-group referred to as RMM (Rec114, Mei4 and Mer2) in S. cerevisiae (Arora et al. 2004; Li et al. 2006; Maleki et al. 2007; Kumar et al. 2010; Stanzione et al. 2016; Tesse et al. 2017; Vrielynck et al. 2021).

We recently showed that the RMM proteins constitute two distinct sub-complexes, a Rec114−Mei4 heterotrimer and a Mer2 homotetramer (Claeys Bouuaert et al. 2021). In vitro, both complexes undergo DNA-driven condensation independently and can mingle together to form mixed condensates. Condensation is a fundamental property of RMM proteins, and mutations that reduce DNA binding compromise condensation in vitro and meiotic DSB formation in vivo, suggesting that this activity is important for their biological function (Claeys Bouuaert et al. 2021). Hence, we proposed that RMM condensation organizes discrete chromatin sub-compartments within which DSB formation takes place. Nevertheless, little was known regarding the structures of Rec114−Mei4 and Mer2 complexes, how they relate to their biological functions, and whether their structural and molecular properties are conserved.

Here, we address this using AlphaFold structural modeling, supported by biochemical and biophysical characterization. We show that Rec114−Mei4 and Mer2 complexes show similar architectures throughout eukaryotes and reveal mechanistic insights into their multivalent interactions with DNA that underlie the assembly of the DSB machinery by DNA-driven condensation.

## Results

### Structure of a minimal Rec114−Mei4 complex

Rec114 and Mei4 form a heterotrimeric complex with a 2:1 stoichiometry where the C-terminus of Rec114 homodimerizes and interacts with the N-terminus of Mei4 (**Fig. 1A**) (Claeys Bouuaert et al. 2021). We purified the Rec114 dimerization domain (residues 375-428) and a minimal trimeric Rec114−Mei4 complex (Mei4 residues 1-43). Thermal shift analyses revealed apparent melting temperatures of 74.5 ± 1.0 °C and 81.5 ± 0.9 °C for Rec114 and the Rec114−Mei4 complex, respectively, indicating that the presence of Mei4 stabilizes Rec114 (**Fig. 1B**).

**Figure 1:**
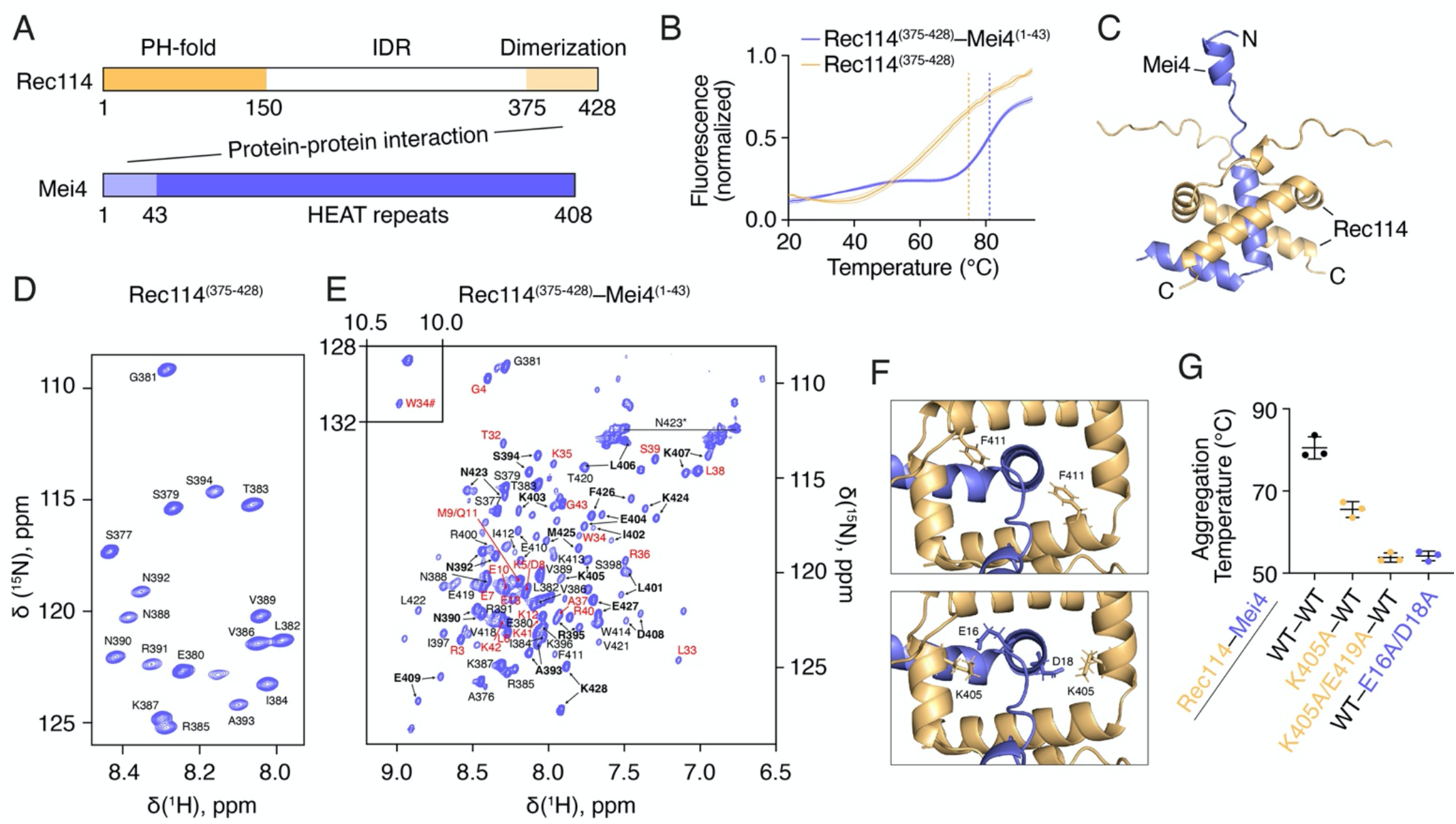
Structure of a minimal Rec114−Mei4 complex. (A) Domain organization of Rec114 and Mei4 proteins. PH domain, Pleckstrin Homology; IDR, intrinsically-disordered region. (B) Thermal shift analyses of Rec114 C-terminal domain and the minimal trimeric Rec114−Mei4 complex. Error bars show mean ± SD of three replicates. The dotted lines indicate the apparent melting temperature for each complex. (C) AlphaFold structure of a minimal Rec114−Mei4 complex. (D) [^1^H,^15^N] HSQC spectrum of the Rec114 C-terminal domain annotated with its backbone amide assignments. (E) [^1^H,^15^N] HSQC spectrum of the minimal trimeric Rec114−Mei4 complex. Black and red labels show backbone amide resonance assignments of Rec114 and Mei4, respectively. The Rec114 residues exhibiting two sets of peaks are in bold. The side-chain NH_2_ peaks of N423 are indicated by an asterisk and joined by a horizontal line. The indole amide resonance of Mei4 W34 is labelled by a hash symbol and shown in the inset. (F) Binding interface between Rec114 and Mei4. Rec114 F411 was previously shown to be important for the interaction with Mei4 and for DSB formation (Claeys Bouuaert et al. 2021). Mei4 residues E16 and D18 make strong hydrogen bonds with Rec114 K405 in both dimer chains. (G) Thermal shift analysis of wild type and mutant Rec114−Mei4 minimal complexes. Error bars show mean ± SD of three replicates.

We used AlphaFold to predict the structure of the minimal Rec114−Mei4 complex (Jumper et al. 2021). This yielded a high-quality model showing two Rec114 α-helical chains (residues 399-426) cupping a Mei4 α-helix (residues 16-29) (**Fig. 1C**). Quality assessment indicated high-confidence predictions of the folded regions and the relative orientations of the interacting domains (**Supplemental Fig. S1**).

To experimentally verify this structural model, we purified isotopically labeled U-[^13^C, ^15^N] Rec114 and Rec114−Mei4 complexes, and studied the proteins by nuclear magnetic resonance (NMR) spectroscopy. Although NMR analyses did not allow us to determine the structure of the complexes *de novo* (**Supplemental Fig. S2A**), the data strongly support the AlphaFold models.

First, AlphaFold predicts that Rec114 and Mei4 peptides feature two α-helices preceded by N-terminal unstructured tails. This topology is confirmed by NMR chemical shift index analysis, which revealed a good agreement between the NMR and AlphaFold α-helical regions (**Supplemental Fig. S2B)**.

Second, the presence of Mei4 breaks the symmetry of the Rec114 dimer. This is clearly seen in the HSQC spectrum of the Rec114−Mei4 complex, where multiple Rec114 residues give rise to two backbone amide resonances (**Fig. 1D, E**). This indicates that the same Rec114 amino acid experiences different chemical environments in the two protein chains. Mapping the differences in chemical shifts of the double Rec114 HSQC peaks shows that the largest effects are observed for the residues in the first α-helix (residues 399-407) and the C-terminal part of the protein (residues 423-428) (**Supplemental Fig. S2C**). Indeed, these are the Rec114 regions with the highest dissimilarity in the predicted structure of the complex, where residues in one protein chain interact with the C-terminal α-helix of Mei4 (residues 32-43), while the same groups in the other chain do not.

Third, we recorded and analyzed nuclear Overhauser effect spectra (NOESY), which allow to detect pairs of ^1^H atoms lying in close spatial proximity (typically < 5Å (Wüthrich 1986)). We successfully assigned a number of sidechain methyl and aromatic groups of both Rec114 and Mei4, which are buried in the core of the protein complex and, thus, can be used to detect specific residue-residue contacts. A set of well-resolved methyl resonances of Leu, Ile, Val, and Met, as well as aromatic protons of Trp and Phe, were inspected in NOESY spectra, which allowed identification of several key ^1^H-^1^H interactions (**Supplemental Fig. S2D**). In total, 13 unambiguous NOEs were detected (**Supplemental Table S1**). In particular, we observed contacts between residues in the same protein chain, interchain Rec114−Rec114 interactions, and several intermolecular Rec114−Mei4 contacts. Overall, the observed NOEs are fully consistent with the predicted structure and validate the AlphaFold model of the minimal heterotrimeric Rec114−Mei4 complex.

Next, we sought to test the model by mutagenesis. We previously showed that a Rec114 F411A mutation abolishes the interaction with Mei4 in yeast-two-hybrid and pulldown assays, and abolishes meiotic DSB formation (Claeys Bouuaert et al. 2021). The AlphaFold model indicates that the F411 residue points directly towards Mei4 (**Fig. 1F**, top), explaining these results. Based on the model, we selected for mutagenesis other residues located within the predicted interaction surface. Rec114 residues K405 and E419, and Mei4 residues E16 and D18 make strong intermolecular hydrogen bonds (**Fig. 1F**, bottom). Alanine substitutions at these positions strongly lowered the aggregation temperature of the complex in a thermal shift assay, indicating that these residues indeed stabilize the complex (**Fig. 1G**).

### Model of full-length Mei4 bound to Rec114

Next, we examined the architecture of the full-length Rec114−Mei4 complex. Rec114 is predicted to have a long central intrinsically-disordered region (IDR) preceded by a structured N-terminal domain (Claeys Bouuaert et al. 2021). The N-terminal domain of *M. musculus* REC114 has been crystallized and shows a Pleckstrin Homology (PH)-like fold composed of an α-helix sandwiched between two antiparallel β-sheets (Kumar et al. 2018; Boekhout et al. 2019). AlphaFold predicted with high confidence a similar structural fold for the yeast Rec114 N-terminal domain (residues 1-140) (**Supplemental Fig. S8A**).

In addition, AlphaFold prediction of full-length Mei4 revealed a structure composed of 19 α-helices, with the two N-terminal Rec114-binding helices pointing out of this ordered structure (**Fig. 2A, Supplemental Fig. S3A-C**). Analysis of the Mei4 structural fold using the DALI server (Holm 2022) revealed similarities with HEAT-repeat proteins, including NOT1 (PDB 5FU7 (Raisch et al. 2016), Cα RMSD of 3.7 Å over 202 residues) and CAND1 (PDB 4A0C (Fischer et al. 2011), Cα RMSD of 3.5 Å over 172 residues). Typical HEAT repeats form an α-solenoid consisting of helix A-turn-helix B motifs arranged in tandem with a ∼15° angle between the repeats (Andrade et al. 2001). Indeed, the predicted Mei4 structure comprises a core of four pairs of antiparallel helices stacked against each other, yielding a convex surface made of A helices and a concave surface made of B helices, similar to other HEAT-repeat structures (**Fig. 2B**). However, these are not assembled from canonical helix-turn-helix motifs, since two out of four pairs of antiparallel helices are interrupted by additional helices (**Fig. 2C**). Hence, Mei4 has an atypical HEAT-repeat structure.

**Figure 2:**
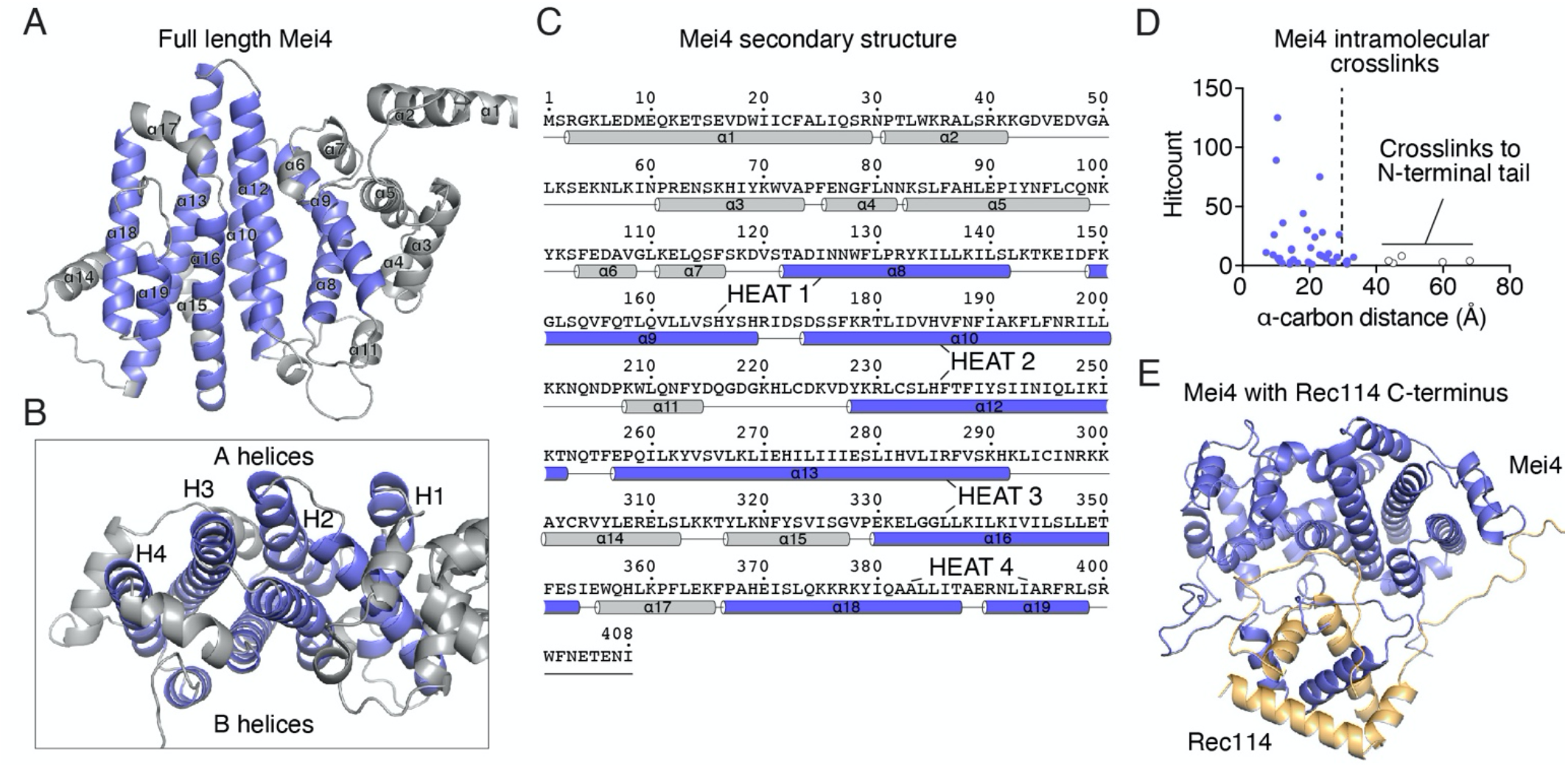
Structural model of full-length Mei4 with Rec114 C-terminus. (A) AlphaFold predicted structure of full-length Mei4 (AF-P29467). Helices that make up the HEAT-repeat core are in blue, other helices are grey. (B) Mei4 has a concave side (top) made up of HEAT-repeat A helices and a convex side (bottom) made up of B helices. (C) Secondary structures of Mei4 based on AlphaFold model. Helices that constitute the HEAT repeats are annotated. (D) Analysis of Mei4 intramolecular crosslinks (690 crosslinks, 44 distinct pairs) (Claeys Bouuaert et al. 2021). The histogram shows the distances separating α-carbons of crosslinked lysines. The crosslinkable limit (dashed line) is 27.4 Å. Model-clashing crosslinks (open circles) involve the flexible N-terminal tail of Mei4. (E) The lowest energy structure of the dimeric Rec114 C-terminus (orange) in complex with full-length Mei4 (blue).

We used our published crosslinking coupled to mass spectrometry (XL-MS) data of Rec114−Mei4 complexes (Claeys Bouuaert et al. 2021) to test the predicted Mei4 structure. The dataset contains 690 intramolecular Mei4 crosslinks (44 distinct pairs). Most of the α-carbons of the crosslinked lysines were closer to each other than the crosslinkable limit of 27.4 Å (**Fig. 2D**). The five pairs of crosslinked residues that were significantly farther apart involved the N-terminal Rec114-interacting tail, which is presumably flexible. Hence, intramolecular Mei4 crosslinks strongly support the predicted structure.

We sought to model the interaction of the Rec114 C-terminal domain with full-length Mei4. To do this, we used the Rec114−Mei4 XL-MS data as distance restraints to calculate the lowest-energy structure for the complex (**Fig. 2E**). This revealed a binding geometry in which the Rec114 C-terminal dimer is docked on the concave side of Mei4, formed by HEAT-repeat B helices (α9, α12, α16, α19). The refined structure satisfies most of the abundant crosslinks (>10 hits) of the XL-MS data (**Supplemental Fig. S3D, E, Supplemental Table S2**). AlphaFold predicted a similar arrangement, although with low confidence in the relative position of the Mei4 HEAT-repeat domain and Rec114−Mei4 interaction domain (**Supplemental Fig. S3F-H**).

### Rec114−Mei4 heterotrimers have two duplex-DNA binding sites

DNA-dependent condensation by Rec114−Mei4 is likely driven by multivalent interactions with DNA (Claeys Bouuaert et al. 2021). *In vitro*, this could result in a binding preference for branched DNA substrates, where the branched substrate provides multiple protein-binding sites that can be occupied simultaneously and stabilize the nucleoprotein complex. Hence, a binding preference for branched DNA can be indicative of multivalent protein-DNA interactions.

To investigate this, we used a gel-shift assay to quantify the affinity of Rec114−Mei4 to different DNA structures assembled with synthetic oligonucleotides. We found that Rec114−Mei4 indeed binds preferentially to branched DNA substrates, compared to duplex DNA (**Fig. 3A, B**). This was confirmed in a competition assay, where a DNA substrate that mimics a Holliday Junction (HJ) was a ∼10-fold better competitor than duplex DNA (**Fig. 3C**). In addition, binding to duplex DNA leads to well-shifts, while binding to the branched substrates leads to complexes that can enter the gel.

**Figure 3:**
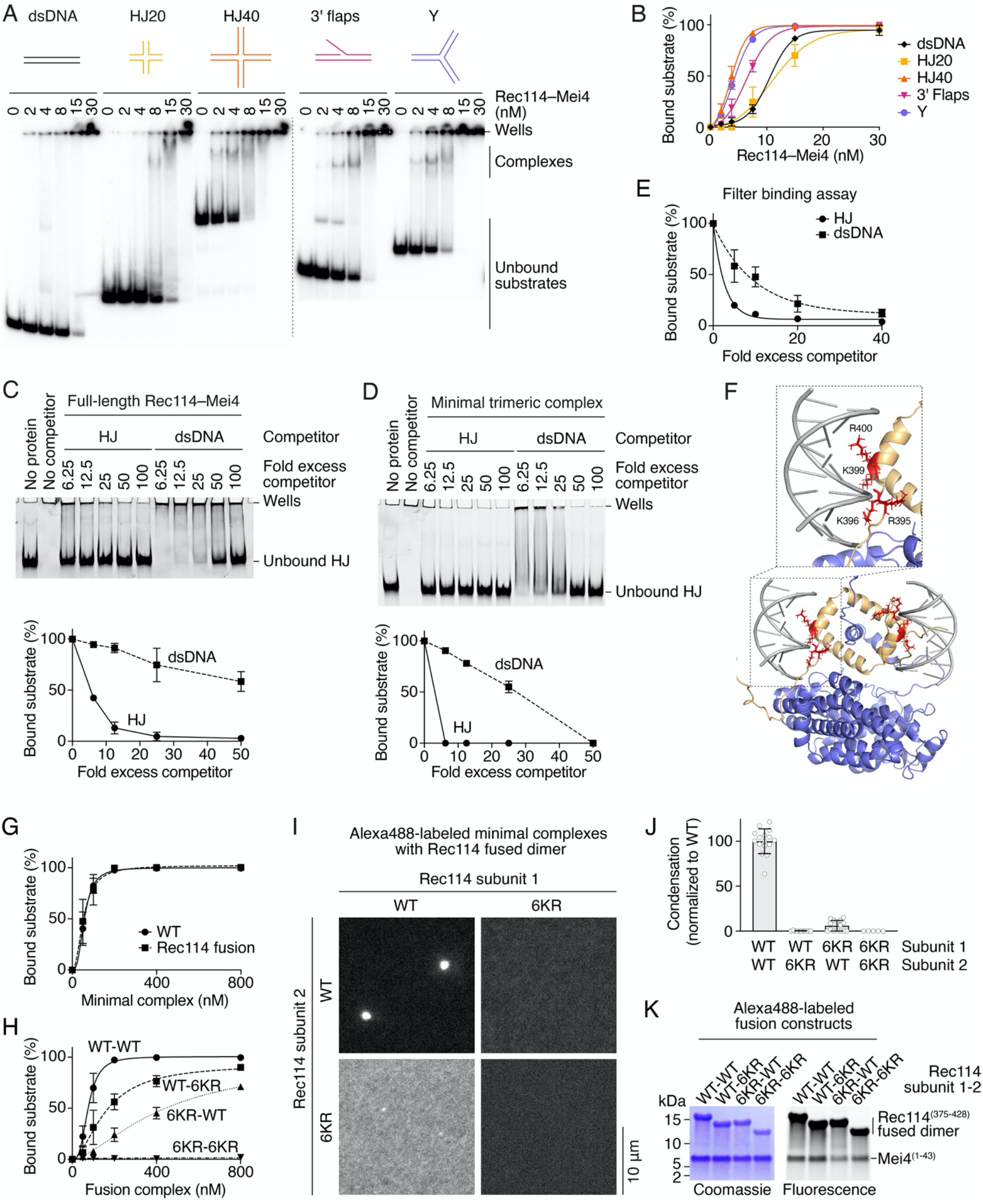
Rec114−Mei4 has two distinct DNA-binding sites. (A) Gel-shift assay of tagged Rec114−Mei4 binding to ^32^P-labeled 80-bp duplex DNA (dsDNA), Holliday Junctions with 20-bp (HJ20) or 40-bp (HJ40) arms, substrates with a 40-nt 3’ single-stranded flap (3’ flaps), or branched structures with three 40-bp arms (Y). Here and elsewhere, concentrations refer to a 2:1 Rec114−Mei4 heterotrimer. (B) Quantification of the gel-shift assay in panel A. Error bars are ranges from two independent experiments. Lines are sigmoidal curves fit to the data. The apparent affinities of Rec114–Mei4 for the DNA substrates are: 10.5 ± 1.4 nM (dsDNA, mean and range), 10.8 ± 3.0 nM (HJ20), 3.5 ± 0.7 nM (HJ40), 6.3 ± 1.0 nM (3’ flaps), 4.1 ± 0.4 nM (Y). (C, D) Competition assays of full-length Rec114−Mei4 (180 nM) (C) and minimal trimeric Rec114−Mei4 (200 nM) (D) complexes binding to a fluorescent HJ substrate (10 nM) in the presence of unlabeled HJ or dsDNA substrates. Error bars are ranges from two independent experiments. (E) Filter binding assay of minimal trimeric Rec114−Mei4 complexes binding to a fluorescent HJ substrate in the presence of unlabeled HJ or dsDNA substrates. Quantifications show the mean ± SD from three replicates. (F) Model of Rec114−Mei4 complex bound to two DNA duplexes. The zoom shows the position of Rec114 4KR residues (R395, K396, K399 and R400), previously implicated in DNA binding (Claeys Bouuaert et al. 2021). (G) Quantification of gel-shift assays of wild type minimal complexes and Rec114 fusion constructs to a fluorescent HJ substrate. (H, J, I) Effect of mutating one or both DNA-binding surfaces within the fused dimer construct on the duplex DNA binding activity (H) and condensation activity (I, J) of the minimal trimeric Rec114−Mei4 complex. Condensation reactions contained 4 μM protein. Each point is the average of the intensities of foci in a field of view, normalized to the overall mean for the wild type. Error bars show mean ± SD of 5-16 images. (K) SDS-PAGE analysis of Alexa488-labeled fusion constructs analyzed in panel I.

To gain further insights into the DNA-binding properties of Rec114−Mei4, we examined whether the minimal trimeric complex also interacts preferentially with branched DNA substrates. Indeed, the minimal complex showed higher affinity for a HJ substrate compared to a duplex DNA in a competition experiment and a filter-binding assay, similar to full-length Rec114−Mei4 (**Fig. 3D, E**). Hence the minimal Rec114−Mei4 complex must have multiple DNA-binding sites.

We previously found that alanine mutations of Rec114 residues R395, K396, K399 and R400 (4KR) compromise the DNA-binding and condensation activities of Rec114−Mei4 (Claeys Bouuaert et al. 2021). These residues form two patches of positively charged amino acids that face in opposite directions on the structural model, revealing two DNA-binding sites within the complex (**Fig. 3F**). This geometrical arrangement indicates that the two sites could not be occupied simultaneously with the same B-form DNA substrate, which explains the preference for branched structures.

In addition, residues K403/K407 and K417/K424 also participate in DNA binding (Liu et al. 2023). Residues K417 and K424 of one monomer contribute in *trans* to the DNA-binding surface composed of residues R395, K396, K399, R400, K403 and K407 (6KR) of the second monomer (**Supplemental Fig. S4A**). Mutating the 6KR residues to alanine abolishes the DNA-binding activity of the minimal complex (**Supplemental Fig. S4B**). However, these mutations do not completely abolish DNA binding by full-length Rec114−Mei4 (**Supplemental Fig. S4C, D**). Surprisingly, *in vivo*, alanine substitutions of K403/K407 and K417/K424 improve the spore viability of the *rec114-4KR* mutant (**Supplemental Fig. S4E**). These data indicate that DNA binding by the heterotrimeric Rec114−Mei4 interaction domain is important, but not essential, and may participate in an unknown regulatory function of the complex.

To test the hypothesis that the two DNA-binding surfaces of Rec114−Mei4 drive condensation, we sought to specifically mutate one of the surfaces and establish the impact on DNA binding and condensation. To achieve this, we produced minimal complexes with two Rec114 subunits separated by a short flexible linker (**Supplemental Fig. S4F, G**). The fusion construct retained wild-type DNA-binding activity (**Fig. 3G, Supplemental Fig. S4H**). Next, we mutated one of the two DNA-binding sites within the complex. As expected, the single-sided mutants retained some DNA-binding activity (**Fig. 3H, Supplemental Fig. S4I**), but condensation was abolished (**Fig. 3I-K**). Thus, the two DNA-binding surfaces of Rec114−Mei4 drive condensation.

### Structural model of Mer2 homotetramers

Mer2 has a central coiled-coil domain that tetramerizes, flanked by N- and C-terminal disordered regions (**Fig. 4A**). We previously proposed that the coiled coil is assembled as pairs of parallel α-helices arranged in an antiparallel configuration (Claeys Bouuaert et al. 2021). To gain further insights, we used AlphaFold to predict the structure of a tetrameric Mer2 coiled-coil domain (residues 41-224). Surprisingly, AlphaFold generated with high confidence a parallel tetrameric model (**Fig. 4B, Supplemental Fig. S5**).

**Figure 4:**
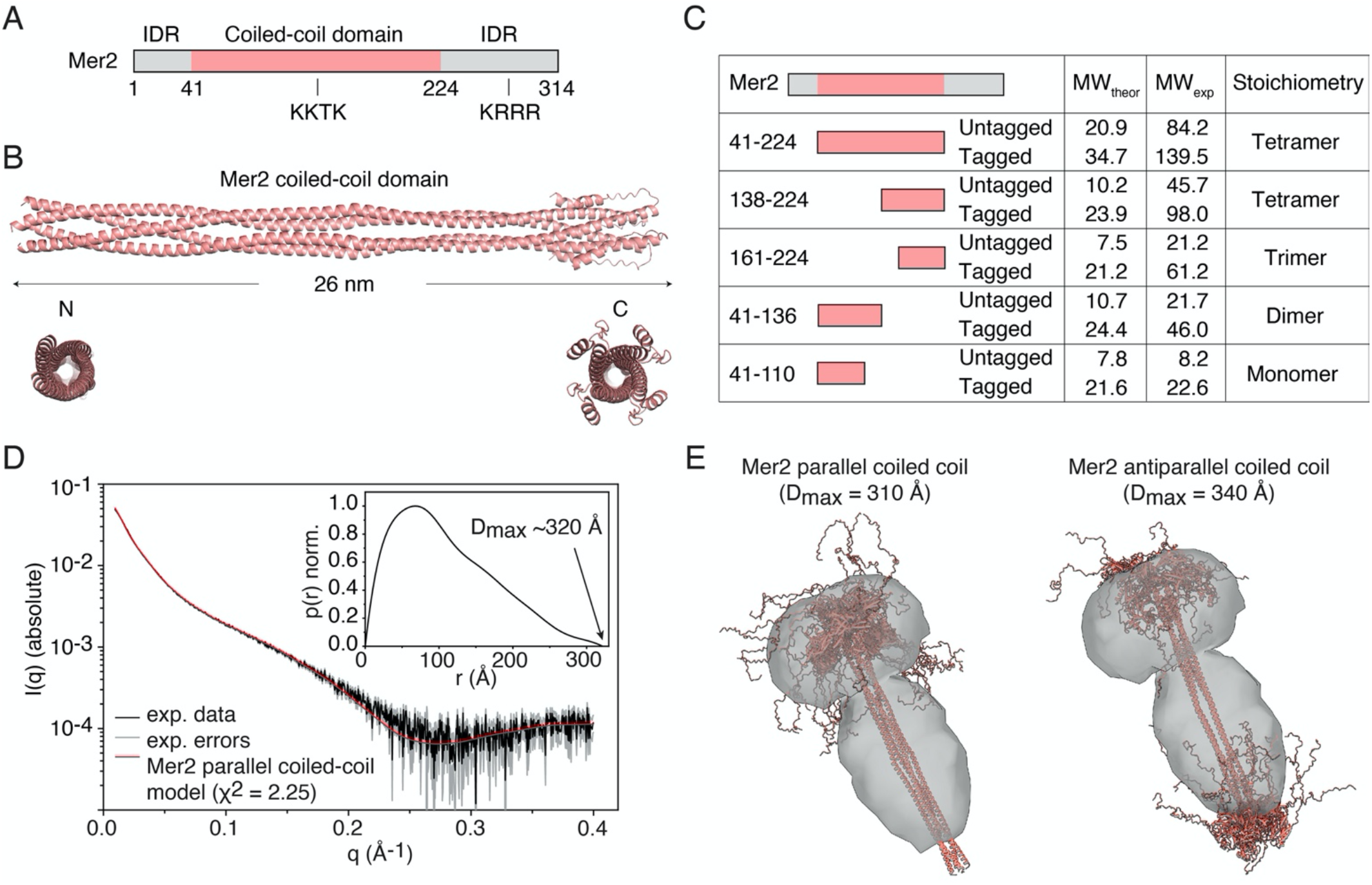
Structural model of Mer2 homotetramers. (A) Domain organization of Mer2. The positions of sequence motifs analyzed in this work are shown. IDR, intrinsically-disordered region. (B) AlphaFold predicted structure of the tetrameric Mer2 coiled-coil domain. (C) SEC-MALS analysis of untagged and HisSUMO-tagged Mer2 truncations (see **Supplemental Fig. S6**). (D) SEC-SAXS analysis of HisSUMO-tagged coiled-coil domain of Mer2. The main graph shows the experimental data (black), error margins (gray), and the fit of the parallel coiled-coil model ensemble to the data (red). The inset shows a normalised probability distance distribution function obtained from the experimental SAXS data, with the approximate D_max_ value indicated. (E) Overlay of parallel and antiparallel model ensembles with the *ab initio* reconstructed shape. The calculated D_max_ values for the model ensembles are indicated for convenience.

To assess this structural arrangement, we purified HisSUMO-tagged and untagged truncations of Mer2 and determined their molecular masses (MW) by size exclusion chromatography followed by multi-angle light scattering (SEC-MALS) (**Fig. 4C, Supplemental Fig. S6**). The Mer2 coiled coil and the C-terminal half alone (residues 138-224) form tetramers, which is not compatible with a parallel-antiparallel model (hereafter ‘antiparallel’). However, further truncating the C-terminal part of the coiled coil (residues 161-224) gives an apparent molecular weight that corresponds to a trimer, suggesting that the tetramer is unstable and partially dissociates during the chromatography. In addition, the N-terminal half of the coiled coil (residues 41-136) yields a molecular weight that corresponds to a dimer, while further truncating the N-terminal yields a monomer, indicating that the packing of the coiled-coil N-terminus is less stable than that of the C-terminus.

Analysis of the predicted structure using Twister (Strelkov and Burkhard 2002) identified about 20 heptad repeats interrupted by six stutters (insertion of 4 amino acids) and one skip (+1) located between residues 80 and 168 (**Supplemental Fig. S13A**). The AlphaFold model predicts geometrical distortions of the coiled coil in the vicinity of the stutters, which are mostly compensated by local unwinding of the coiled coil, with a local shift to a right-handed geometry. In addition, the model shows a wide coiled-coil radius around residues 120-150, which is indicative of sub-optimal hydrophobic packing of the coiled coil, resulting in a less stable structure (**Supplemental Fig. S13B**). Indeed, the confidence score of the AlphaFold model is lowest around the center of the coiled coil (**Supplemental Fig. S5C**).

We sought to definitively establish whether the coiled-coil domain of Mer2 indeed adopts a parallel configuration. To do this, we analyzed a HisSUMO-tagged Mer2 coiled-coil domain by small-angle X-ray scattering (SAXS). Reporting on the overall molecular shape in solution, SAXS provides a means to discriminate between the parallel (all fusion domains on the same side of the coiled coil) and the antiparallel (fusion domains at both ends) topologies. SAXS scattering curves were in excellent agreement with the profile expected for a parallel coiled coil (**Fig. 4D**). In addition, *ab initio* particle reconstitution from the experimental SAXS data shows a clear excess density at one end, consistent with a parallel configuration (**Fig. 4E**). Thus, the SAXS analysis validates the AlphaFold model and confirms the parallel arrangement of α-helices within Mer2.

### Mer2 engages in multivalent protein-DNA interactions

Mer2 was previously shown to bind double-stranded DNA *in vitro*, but binding preferences for distinct DNA structures had not been investigated (Tsai et al. 2020; Claeys Bouuaert et al. 2021; Rousová et al. 2021). Similar to Rec114−Mei4, we found that Mer2 binds preferentially to branched substrates (**Fig. 5A, B**). This was confirmed in a gel-shift competition and filter binding assays, which showed ∼10-20-fold increased affinity for an HJ substrate compared to duplex DNA (**Fig. 5C, D**).

**Figure 5:**
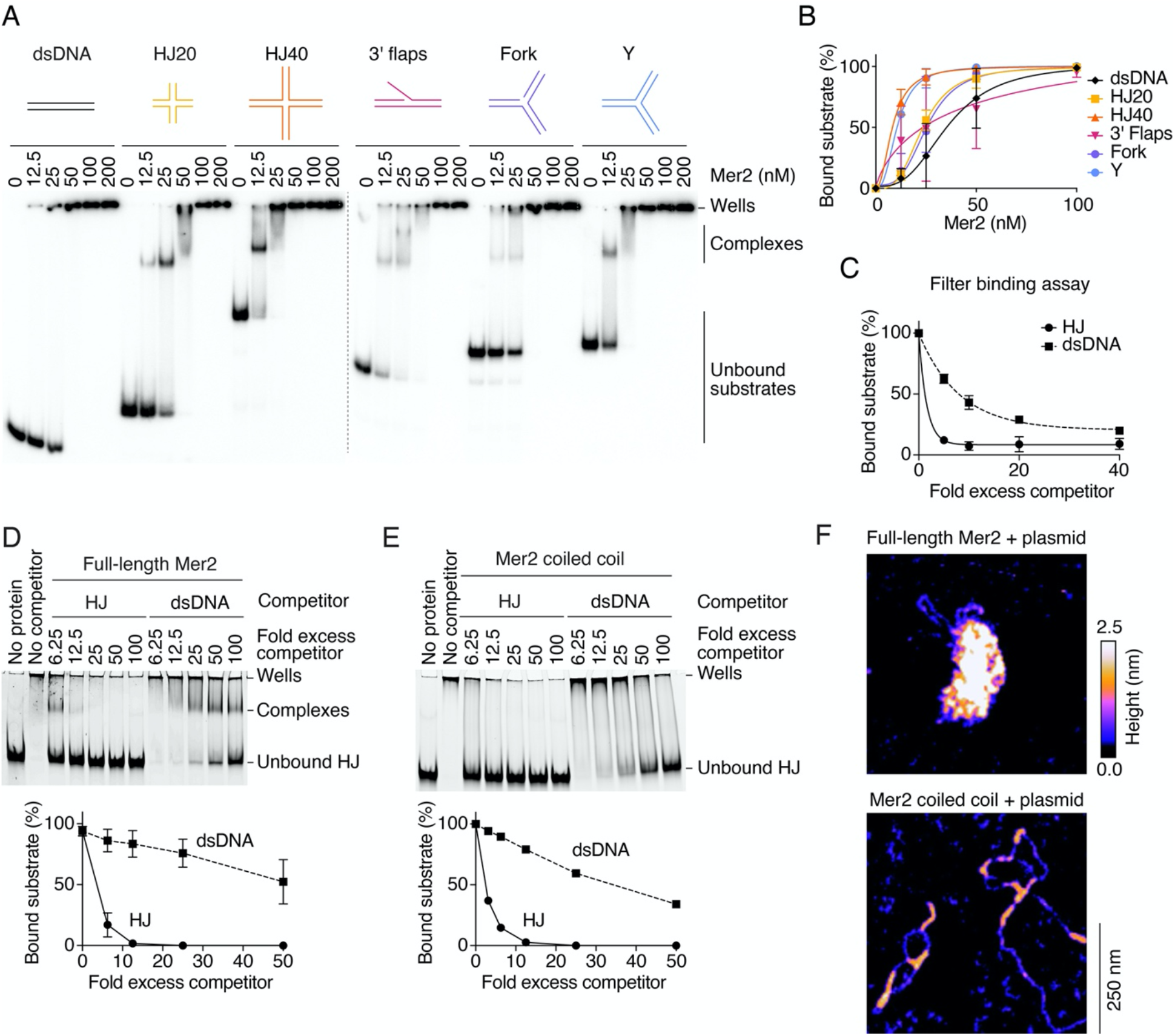
DNA-binding activities of Mer2. (A) Gel-shift assay of Mer2 binding to ^32^P-labeled 80-bp duplex DNA (dsDNA), Holliday Junctions with 20-bp (HJ20) or 40-bp (HJ40) arms, substrates with a 40-nt 3’ single-stranded flap (3’ flaps), or branched structures with three 40-bp arms without (Y) and with a central nick (Fork). Concentrations refer to Mer2 monomers. (B) Quantification of the gel-shift assay in panel A. Error bars are ranges from two independent experiments. Lines are sigmoidal curves fit to the data. The apparent affinities of Mer2 for the DNA substrates are: 36.8 ± 13.3 nM (dsDNA, mean and range), 23.9 ± 2.9 nM (HJ20), 9.2 ± 1.0 nM (HJ40), 34.4 ± 25.2 nM (3’ flaps), 24.4 ± 5.5 nM (Fork), 10.6 ± 1.2 nM (Y). (C) Filter binding assay of Mer2 binding to a fluorescent HJ substrate in the presence of unlabeled HJ or dsDNA substrates. Quantifications show mean ± SD from three replicates. (D, E) Competition assays of full-length Mer2 (200 nM) (D) or the coiled-coil domain (750 nM) (E) binding to a fluorescent HJ substrate (10 nM) in the presence of unlabeled dsDNA or HJ40 substrates. Error bars are ranges from two independent experiments. Error bars in panel E are too small to be visible. (F) AFM imaging of full-length Mer2 (100 nM) or the coiled-coil domain (100 nM) in the presence of 1 nM plasmid DNA (pUC19).

The Mer2 coiled-coil domain, which is necessary and sufficient for DNA binding (Claeys Bouuaert et al. 2021), also binds preferentially to branched DNA substrates (**Fig. 5E**). This indicates that the coiled-coil domain engages in multivalent protein-DNA interactions. Consistently, atomic force microscopy (AFM) imaging of the coiled-coil domain in the presence of plasmid DNA revealed that the coiled coil has DNA-bridging activity, while full-length Mer2 assembles higher-order nucleoprotein condensates (**Fig. 5F**).

A KKTK motif located at the center of the coiled coil (**Fig. 4A**) was previously implicated in DNA binding (Tsai et al. 2020). However, in our hands, mutating the three lysines of this motif to alanine did not affect the DNA-binding activity of Mer2 (**Supplemental Fig. S7A**). Hence, the DNA-binding residues within the Mer2 coiled-coil domain remain unknown. On the other hand, we previously showed that the KRRR motif located within the C-terminal IDR is important for DNA binding, condensation, and DSB formation (Claeys Bouuaert et al. 2021). As expected, the Mer2-KRRR mutant showed a decreased affinity for a branched substrate (**Supplemental Fig. S7A**). Consistently, yeast strains harboring a *mer2-KKTK* mutation have wild-type spore viability, while the *mer2-KRRR* mutant does not yield any viable spores (**Supplemental Fig. S7B**).

### Conservation of Rec114−Mei4 structure and DNA-binding properties

The RMM proteins are highly diverged across the eukaryotic kingdom, and pair-wise comparisons between homologs of distantly related species typically show sequence identities well below 10% (Kumar et al. 2010; Stanzione et al. 2016; Tesse et al. 2017; Wang et al. 2020). To gain insights into their structural conservation, we used AlphaFold to model the architecture of Rec114 and Mei4 orthologs from *M. musculus* (REC114, MEI4), *S. pombe* (Rec7, Rec24), *A. thaliana* (PHS1, PRD2) and *Z. mays* (PHS1, MPS1) (Molnar et al. 2001; Pawlowski et al. 2004; De Muyt et al. 2009; Kumar et al. 2010; Steiner et al. 2010; Bonfils et al. 2011; Rosu et al. 2013; Stamper et al. 2013; Hinman et al. 2021; Vrielynck et al. 2021).

As expected, Rec114 orthologs showed a long central IDR flanked by an N-terminal PH domain, and 2-4 C-terminal α-helices (**Supplemental Fig. S8A**). While the PH-fold was well defined for the *M. musculus, S. pombe*, and *Z. mays* orthologs, that of *A. thaliana* PHS1 was incomplete. Interestingly, PHS1 was recently shown to be dispensable for meiotic DSB formation (Vrielynck et al. 2021), in contrast to other Rec114 orthologs, including maize PHS1 (Molnar et al. 2001; Pawlowski et al. 2004; Rosu et al. 2013; Stamper et al. 2013; Kumar et al. 2018).

AlphaFold models of Mei4 orthologs revealed HEAT-repeat domains flanked by 3-5 flexibly-connected N-terminal helices (**Supplemental Figs. S8B & 9**). For each ortholog, four well-defined HEAT repeats could be identified (**Supplemental Fig. S9**). The helix A-turn-helix B motifs were more canonical than for yeast Mei4, and consecutive HEAT repeats were arranged with a 5-15° angle, creating slightly curved structures (**Supplemental Fig. S8B**).

Next, we modeled the interaction domains of Rec114 and Mei4 orthologs. In all cases, AlphaFold predicted similar heterotrimeric complexes with an N-terminal α-helix of the Mei4 ortholog inserted within a dimeric ring composed of C-terminal α-helices of Rec114 orthologs (**Fig. 6A, Supplemental Fig. S10**). The conserved phenylalanine equivalent to *S. cerevisiae* Rec114 F411 (Kumar et al. 2010; Steiner et al. 2010; Claeys Bouuaert et al. 2021) occupies a similar position inside the dimeric ring in all the models (**Fig. 6A**, bottom).

**Figure 6:**
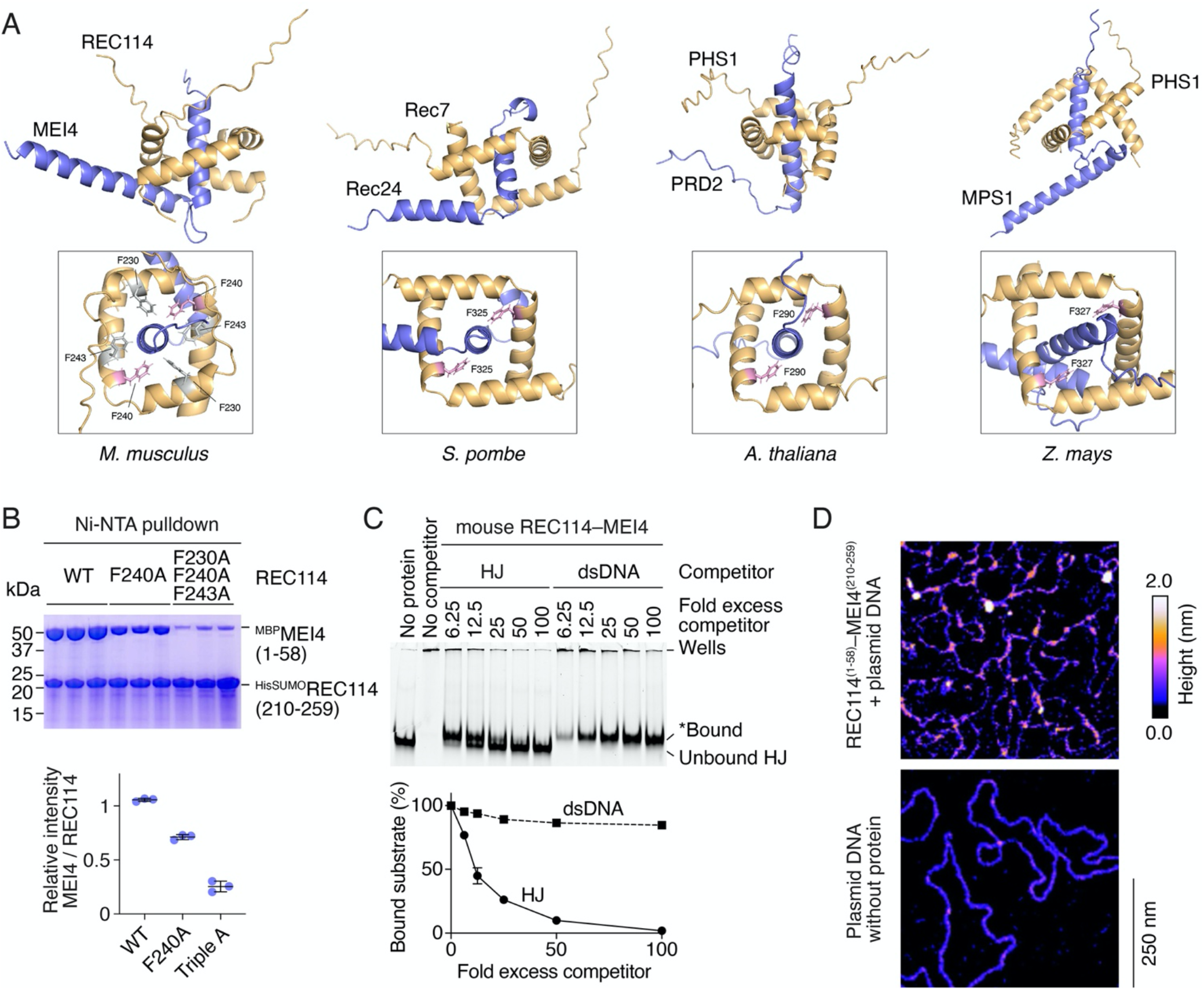
Conservation of Rec114−Mei4 structure and DNA-binding properties. (A) AlphaFold models of minimal trimeric complexes for Rec114−Mei4 orthologs in *M. musculus* (REC114−MEI4), *S. pombe* (Rec7−Rec24), *A. thaliana* (PHS1−PRD2), and *Z. mays* (PHS1−MPS1). The position of the conserved phenylalanine at the interface between Rec114 and Mei4 orthologs is shown (pink). Other residues mutated in panel B are shown in grey. (B) Pulldown analysis of the effect of *M. musculus* REC114 F230A, F240A, and F243A mutations on the interaction with MEI4. Error bars are ranges from three replicates. (C) Competition assay of tagged mouse REC114−MEI4 complex (1000 nM) binding to a fluorescent HJ substrate (10 nM) in the presence of unlabeled dsDNA or HJ substrates. The band labeled ‘*Bound’ migrates close to the position of the unbound substrate. It is likely due to the rapid dissociation of REC114−MEI4 from the substrate at the start of the electrophoresis. Error bars are ranges from two independent experiments (most are too small to be visible). (D) AFM imaging of minimal trimeric mouse REC114−MEI4 complexes in the presence of 1 nM plasmid DNA (pUC19).

To test these models, we co-expressed tagged complexes consisting of the C-termini of Rec114 orthologs and N-termini of Mei4 orthologs, and assayed protein-protein interactions by pulldown. Complexes could be purified for all orthologs, confirming that the respective domains interact (**Fig. 6B, Supplemental Fig. S11**). Next, we tested the AlphaFold models by mutagenesis. As expected, mutating the conserved phenylalanine (F240) of *M. musculus* REC114 to alanine decreased the interaction with MEI4, and this effect was exacerbated by mutating two additional phenylalanine residues (F230, F243) within the dimeric ring (**Fig. 6B**). Similarly, mutating three hydrophobic residues within the dimeric ring compromised the interaction between *S. pombe* Rec7 and Rec24 (**Supplemental Fig. S11A**), and *A. thaliana* PHS1 and PRD2 (**Supplemental Fig. S11B**), providing support for the AlphaFold models. However, a triple alanine mutation within the dimeric ring of *Z. mays* PHS1 did not significantly impact the interaction with MPS1 (**Supplemental Fig. S11C**).

To address whether the DNA-binding properties of Rec114−Mei4 are conserved, we performed gel-shift analyses with full-length *M. musculus* REC114−MEI4 in the presence of an HJ substrate. Similar to the yeast complex, REC114−MEI4 binds to an HJ substrate with >20-fold higher affinity than to duplex DNA (**Fig. 6C**), suggesting that the complex contains multiple DNA-binding sites. Consistently, AFM imaging revealed that a minimal mouse REC114−MEI4 complex has DNA-bridging activity (**Fig. 6D**). On the other hand, gel-shift analyses revealed significant DNA-binding activity only for the minimal complex of *A. thaliana* PHS1-PRD2 (**Supplemental Fig. S12A**), indicating that the DNA-binding activities of the minimal complexes are less robust than that of yeast Rec114−Mei4. This is likely explained by the fact that the electrostatic surface potential of the *S. cerevisiae* minimal complex is significantly more positive than that of the other orthologs (**Supplemental Fig. S12B, D**). Nevertheless, the C-terminus of all the Rec114 orthologs have positively-charged, putative DNA-binding residues that point outwards on the structural models (**Supplemental Fig. S12C**). This suggests that the mechanism for multivalent protein-DNA interactions is conserved in Rec114−Mei4 orthologs, albeit with quantitatively distinct activities.

### Conservation of Mer2 structure and DNA-binding properties

To investigate the structural conservation of Mer2 orthologs, we generated AlphaFold predictions of tetrameric coiled-coil domains of *M. musculus* IHO1, *A. thaliana* PRD3, *S. pombe* Rec15, *S. macrospora* ASY2, and *Z. mays* PAIR1 (Nonomura et al. 2004; De Muyt et al. 2009; Miyoshi et al. 2012; Stanzione et al. 2016; Tesse et al. 2017). AlphaFold modeling yielded high-confidence predictions of the different coiled coils, all arranged in a parallel configuration (**Fig. 7A, Supplemental Fig. S13**).

**Figure 7:**
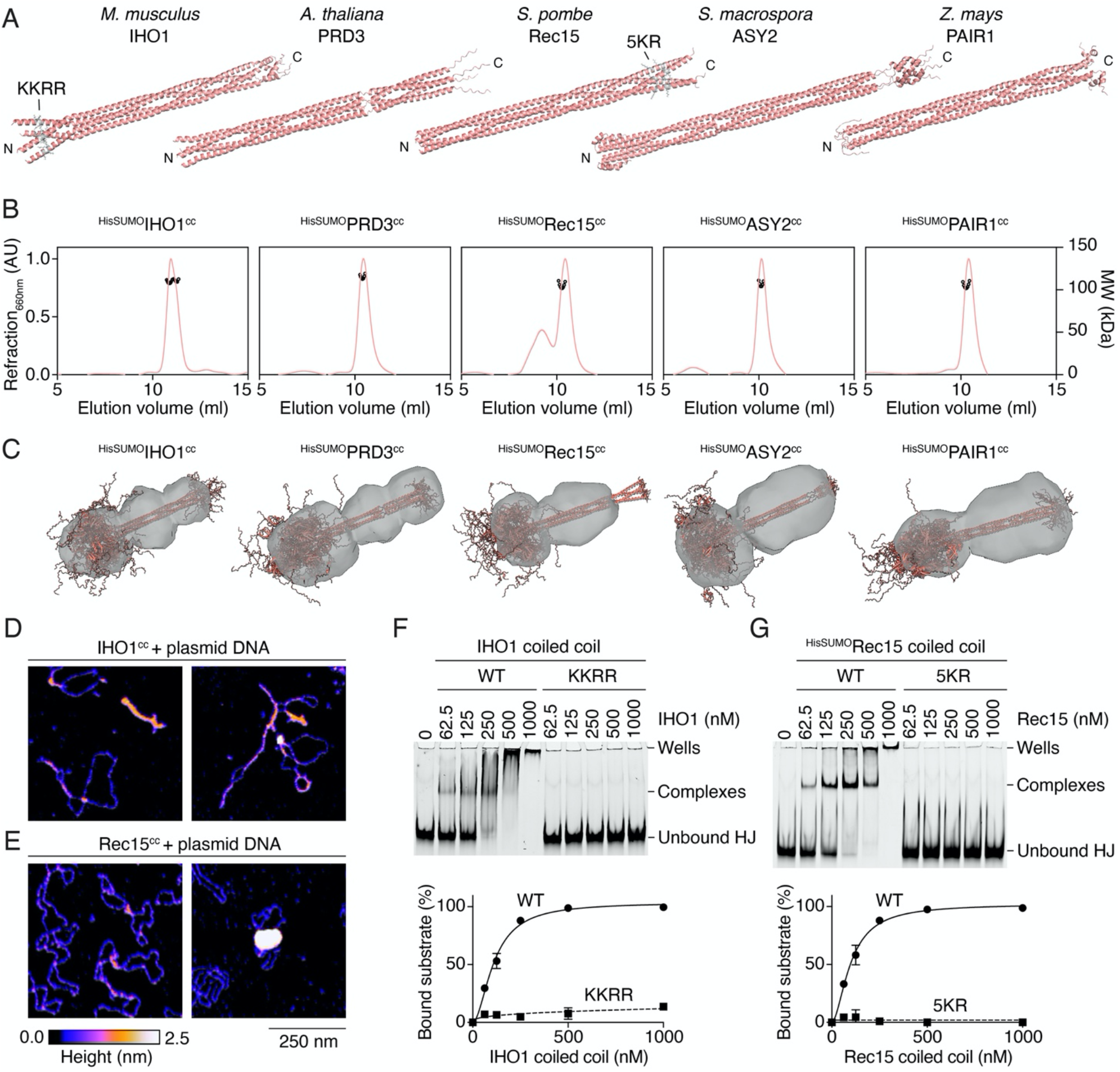
Conservation of Mer2 structure and DNA-binding properties. (A) AlphaFold models of homotetrameric coiled-coil domains of *M. musculus* IHO1 (residues 109-267, length 2.15 nm), *A. thaliana* PRD3 (residues 120-270, length 2.25 nm), *S. pombe* Rec15 (residues 1-160, length 2.3 nm), *S. macrospora* ASY2 (residues 55-275, length 2.85 nm), and *Z. mays* PAIR1 (residues 140-310, length 2.15 nm). IHO1 and Rec15 residues mutated in panels F and G are indicated (grey). (B) SEC-MALS analysis of HisSUMO-tagged coiled-coil domains (boundaries are identical to panel A). The traces show differential refraction at 660 nm (arbitrary units) and circles are molar mass measurements across the peak. Experimental molecular weight and expected molecular weight based on a tetrameric stoichiometry are as followed: IHO1: 113 kDa (expected 127 kDa); PRD3: 135 kDa (expected 129 kDa); Rec15: 132 kDa (expected 128 kDa); ASY2: 142 kDa (expected 155 kDa); PAIR1: 122 kDa (expected 132 kDa). (C) Overlay of parallel models of HisSUMO-tagged coiled coils with the *ab initio* reconstructed shape obtained from SEC-SAXS analyses. (D, E) AFM imaging of the coiled-coil domain of (D) IHO1 (100 nM) or (E) Rec15 (left 50 nM, right 100 nM) in the presence of 1 nM plasmid DNA (pUC19). (F, G) Gel-shift assays of wild type and mutant IHO1 (F) and Rec15 (G) coiled-coil domains binding to a fluorescent HJ substrate. The IHO1-KRRR mutant has four positively charged residues (K121, K122, R123, R124) located towards the N-terminus of the coiled coil mutated to alanine. The Rec15-5KR mutant has five positively charged residues (K134, K135, K141, K142, R143) located towards the C-terminus of the coiled coil mutated to alanine. Error bars are ranges from two independent experiments (most are too small to be visible).

Similar to *S. cerevisiae* Mer2, Twister analyses (Strelkov and Burkhard 2002) of the predicted coiled-coil structures identified heptad repeats and revealed the presence of discontinuities (stutters, skips and stammers), which are locally compensated by distortions of the coiled coils, primarily through increased radius of the coiled coils and local switches to right-handed helices (**Supplemental Figs. S14-16**). Hence the coiled-coil structure of Mer2 orthologs appears to be largely conserved.

To test these models, we expressed the five coiled-coil domains with N-terminal HisSUMO tags and analyzed purified complexes by SEC-MALS. This yielded experimental molecular masses consistent with homotetrameric complexes (**Fig. 7B**).

Next, we analyzed the coiled-coil domains by SEC-SAXS. For all the tagged coiled coils, SAXS curves and overall dimensions of the particles were in excellent agreement with structural models, and *ab initio* particle reconstitutions show an excess density at one end, consistent with parallel configurations (**Fig. 7C, Supplemental Fig. S17**). Thus, the SAXS analysis strongly supports the parallel homotetrameric coiled-coil models of the Mer2 orthologs.

Similar to Mer2, the coiled-coil domains of IHO1 and Rec15 were sufficient to bind DNA and showed a strong binding preference for an HJ substrate compared to duplex DNA in a competition assay (**Supplemental Fig. S18A-C**). PAIR1 also binds DNA but with significantly lower affinity (**Supplemental Fig. S18A**). Consistent with the interpretation that IHO1 and Rec15 coiled coils mediate multivalent protein-DNA interactions, AFM imaging analyses revealed that the coiled coils have DNA-bridging activity (**Fig. 7D, E**).

On the structural models, the helices dissociate at the N-terminus of IHO1 and the C-terminus of Rec15 over a 20-25 amino-acid sequence that contains patches of positively charged residues (**Fig. 7A**). Mutating these residues to alanine abolished the DNA-binding activity of the coiled coils (**Fig. 7F, G**). Hence, IHO1 and Rec15 share DNA-binding properties with Mer2, although the position of the key DNA-interacting residues may differ.

Finally, we observed by AFM analyses that the Rec15 coiled coil assembles condensates in the presence of DNA (**Fig. 7E**, center), but IHO1 does not. This is likely due to the formation of intermolecular interactions between Rec15 tetramers. Indeed, SEC-MALS and SAXS indicated that Rec15 and IHO1 coiled coils have similar molecular weights (**Fig. 7B, Supplemental Table. S7**), but a volume analysis from AFM images shows that Rec15 forms significantly larger particles than IHO1 (**Supplemental Fig. S18D**). Hence, the coiled-coil domain of Rec15 mediates low-affinity inter-tetrameric interactions that facilitate DNA-dependent condensation.

## Discussion

We presented structural models of *S. cerevisiae* Rec114−Mei4 and Mer2 complexes, supported by NMR, SAXS, and mutagenesis, and showed that their architecture is conserved in higher eukaryotes. In addition, we showed that Rec114−Mei4 and Mer2 complexes and their orthologs bind preferentially to branched DNA structures, which results from the combined action of multiple DNA-binding sites within the complexes. These multivalent interactions are key to their condensation activity that drives the assembly of the DSB machinery.

### Structural conservation of Rec114−Mei4 heterotrimers

Rec114 has an N-terminal PH domain, followed by a ∼250 amino-acid IDR. The C-terminal ∼30 amino acids form two α-helices that homodimerize to form a ring in which the N-terminus of Mei4 is inserted, yielding an asymmetric 2:1 complex. Mei4 presents an atypical HEAT-repeat like fold made up of four HEAT repeats that create a curved structure, with the C-terminal Rec114 dimer lodged along the concave side. The Rec114 PH domain, the Mei4 HEAT repeat structure, and the 2:1 Rec114−Mei4 interaction domain are generally conserved in *S. pombe, M. musculus, A. thaliana*, and *Z. mays*. In addition, AlphaFold modeling revealed a similar trimeric structure for the *C. elegans* homologs (DSB-1, DSB-2 and DSB-3) (Guo et al. 2022).

While the yeast Mei4 structure presents non-canonical HEAT repeats, the four HEAT repeats that compose the core structure of Mei4 orthologs displayed more typical helix A-turn-helix B motifs. The *Z. mays* Mei4 homolog, previously identified as Multipolar Spindle 1 (MPS1) (Kumar et al. 2010), also presents a similar HEAT-repeat architecture and N-terminal Rec114 (PHS1)-interacting helices; although, to our knowledge, its meiotic function has not been established. The AlphaFold model of *Z. mays* PHS1-MPS1 was of lower confidence than that of the other orthologs. Nevertheless, purification of a minimal PHS1-MPS1 complex confirmed that the two subunits interact, indicating that MPS1 is likely a *bona fide* Mei4 ortholog.

In contrast to most Rec114 orthologs, AlphaFold modeling of *A. thaliana* PHS1 suggested that its PH-domain is incomplete. This is surprising because PHS1 contains signature sequence motifs (SSMs) within its N-terminal domain that are shared with other Rec114 homologs (Kumar et al. 2010), and these SSMs constitute the core secondary structures of the PH fold (Kumar et al. 2018; Boekhout et al. 2019). Interestingly, *A. thaliana* PHS1 was recently shown to be dispensable for meiotic DSB formation (Vrielynck et al. 2021). Nevertheless, this does not seem to be shared with other plant species, since the predicted maize PHS1 structure shows a well-folded PH-domain, and maize PHS1 is required for meiotic DSB formation (Pawlowski et al. 2004).

### Structural conservation of Mer2 homotetramers

Mer2 forms a homotetramer with a ∼200 amino-acid central coiled coil that folds as a ∼25 nm parallel α-helical bundle. By SAXS analyses, we demonstrated that this parallel tetrameric configuration is conserved in *S. pombe, S. macrospora, M. musculus, Z. mays*, and *A. thaliana*.

Based on two lines of evidence, we previously proposed a parallel-antiparallel arrangement for the Mer2 tetrameric coiled coil (Claeys Bouuaert et al. 2021). First, XL-MS experiments revealed long-range crosslinks within the coiled-coil domain. Second, an engineered single-chain dimer with two copies of the coiled coil connected with a short flexible linker behaved as a tetramer in SEC-MALS analyses. Since the linker was too short to allow parallel folding of the coiled coil, we concluded that Mer2 tetramers most likely consisted of two pairs of parallel α-helices arranged in an antiparallel configuration. However, we show here that this is not the case. A possible explanation is that the coiled coil is flexible, which is consistent with the presence of multiple stutters and a skip that span the central region of the heptad repeats and likely destabilize the coiled coil. However, SAXS analyses suggested that Mer2 adopts a straight, uninterrupted parallel coiled-coil configuration in solution. Hence, our interpretation is that the fused dimeric construct forced the coiled coil to bend. The long-range crosslinks observed by XL-MS (Claeys Bouuaert et al. 2021), on the other hand, can be explained by intermolecular interactions between Mer2 tetramers.

### A model for Mer2-mediated condensation

DNA-dependent condensation by Rec114−Mei4 and Mer2 require protein-DNA interactions, protein-protein interactions, and multivalency. Theoretically, multivalency may arise from protein-protein interactions, protein-DNA interactions, or both. We previously identified DNA-binding residues within Rec114−Mei4 and Mer2 complexes (Claeys Bouuaert et al. 2021). However, insights into the nature of the multivalency remained limited. The preference of Rec114−Mei4 and Mer2 for branched DNA structures and the formation of defined stoichiometric complexes with these substrates indicates that they mediate multivalent protein-DNA interactions.

DNA binding by Mer2 involves the coiled-coil domain and an essential KRRR motif located within the C-terminal IDR. By AFM analysis, we found that the coiled-coil domain has DNA-bridging activity, but does not support condensation, in contrast to full-length Mer2. This is consistent with the hypothesis that the IDRs mediate low-affinity protein-protein interactions that drive condensation (**Supplemental Fig. S19**).

Although the DNA-binding residues within the coiled coil are yet to be identified, Mer2 has 21 lysines and 7 arginines spread along the entire length of the coiled coil. Hence, we propose that the DNA-bridging activity arises from multiple interactions of the tetrameric bundle with the phosphoribose backbone of two coaligned DNA molecules (**Supplemental Fig. S19**). This model also explains the binding preference of the complex for branched DNA.

The coiled-coil domains of IHO1 and Rec15 also bind preferentially to branched DNA substrates and have DNA-bridging activity, suggesting similar DNA-binding modes as Mer2. In these cases, we identified key DNA-binding residues, located at the N-terminus of the IHO1 coiled coil and at the C-terminus of the Rec15 coiled coil. We hypothesize that IHO1 and Rec15 mediate similar geometrical arrangements with co-aligned DNA molecules as Mer2, but the relative importance of different amino acid residues vary between species.

Finally, using AFM analyses of the Rec15 and IHO1 coiled-coil domains, we correlated the formation of condensates with the presence of low-affinity protein-protein interactions within the respective domains. That is, Rec15 showed inter-tetrameric interactions and condensation, while IHO1 did not show any. Like Mer2, the coiled coil of IHO1 is flanked by long IDRs, which are absent from our purified complexes. It is likely that these participate in protein-protein interactions, converting the DNA-bridging activity of the coiled-coil domain into a condensation activity.

Hence, our work suggests that the essential function of Mer2 in assembling the DSB machinery through DNA-driven condensation is likely conserved across eukaryotes.

### Functions of the DNA-binding activity of Rec114−Mei4

The AlphaFold model explains structurally how Rec114−Mei4 engages in multivalent protein-DNA interactions, revealing two duplex DNA binding sites that point in opposite directions. Since those cannot be occupied simultaneously by a single DNA duplex (under the persistence length of ∼150 bp), the complex binds more stably to a branched substrate presenting multiple flexibly-connected duplexes, which can be contacted simultaneously. Using an artificial fusion construct, we demonstrated that the combined action of two DNA-binding surfaces drive Rec114−Mei4 condensation *in vitro*.

Each DNA-binding surface of the complex is made up of eight positively-charged residues of Rec114, with functional relevance confirmed biochemically by mutational analyses (Claeys Bouuaert et al. 2021; Liu et al. 2023). However, we found that mutating these residues does not completely abolish Rec114 function *in vivo*, as determined by spore viability.

We previously reported that a *rec114-4KR* allele was defective for meiotic DSB formation, leading to inviable spores (Claeys Bouuaert et al. 2021). However, in those strains, Rec114 was tagged with a C-terminal myc-tag. The myc-tagged *REC114* allele has been shown to lead to synthetic defects when combined with *spp1, rft1* or *cdc73* deletions (Zhang et al. 2020). Hence, the myc-tag exacerbated the phenotype of the *rec114-4KR* mutant.

By comparing the phenotypes of untagged *rec114-4KR, rec114-6KR*, and *rec114-8KR* yeast mutants, we found, astonishingly, that increasing the number of mutations increases the spore viability of the strain. Hence, the DNA-binding activity of the minimal trimeric Rec114−Mei4 domain is important, but not essential, and may participate in an as-yet unidentified regulatory function of the complex.

While the yeast Rec114−Mei4 complex has robust DNA-binding activity, this was not the case for its orthologs from *M. musculus, S. pombe, A. thaliana* and *Z. mays*. Nevertheless, the mouse orthologs retained DNA-bridging activity, indicating similar – albeit quantitatively different – DNA-binding modes. Hence, it is likely that the DNA-binding activities of Rec114−Mei4 orthologs also play supportive roles in meiotic DSB formation. Nevertheless, Rec114 and Mei4 orthologs carry additional functions, including protein-protein interactions with other partners (Miyoshi et al. 2012; Boekhout et al. 2019; Claeys Bouuaert et al. 2021; Nore et al. 2022; Laroussi et al. 2023).

Structurally, the protein-protein interactions between Rec114−Mei4 and Mer2 complexes remain to be established. While interactions between these partners can be detected *in vitro*, stoichiometric Rec114−Mei4−Mer2 complexes have not been purified (Claeys Bouuaert et al. 2021). On the other hand, a recent study showed that the mouse REC114 PH domain binds to the N-terminus of IHO1 (Laroussi et al. 2023), which contains a conserved sequence motif (Tesse et al. 2017). Hence, a similar protein-interaction mode might be expected for the yeast proteins.

We propose that Mer2 condensation is the primary driver of assembly of the DSB machinery. The DNA-binding activity of Rec114 contributes a non-essential function, which probably depends on protein-protein interactions with Mer2. This is consistent with the observation that chromatin binding by Rec114 requires Mer2, while Mer2 can bind chromatin in the absence of Rec114 and Mei4 (Panizza et al. 2011). In addition, Rec114 chromatin association is contingent on CDK-dependent Mer2 phosphorylation, which promotes protein-protein interactions between Mer2 and Rec114 (Henderson et al. 2006; Panizza et al. 2011). In summary, our results yield insights into the structure of Rec114−Mei4 and Mer2 complexes and the multivalent protein-DNA interactions that drive their DNA-dependent condensation activity and reveal that these structural and functional properties are conserved throughout eukaryotes.

## Materials and Methods

### Preparation of expression vectors

The sequences of the oligos are listed in **Supplemental Table S3** and gBlocks (Integrated DNA Technologies) are listed in **Supplemental Table S4**. Plasmids are listed in **Supplemental Table S5**. The vectors used to express ^HisFlag^Rec114 (pCCB789), ^HisFlag^Rec114-4KR (pCCB848), ^MBP^Mei4 (pCCB791) from Sf9 cells and ^HisSUMO^Mer2 (pCCB750) from *E. coli* were previously described (Claeys Bouuaert et al. 2021).

The Rec114 C-terminal domain and Mei4 N-terminal domain were amplified from pCCB649 and pCCB652 using primers dd015 and dd016, and dd017 and dd018, and cloned into pCCB206 vector by Gibson assembly to yield pDD003 and pDD004 respectively. The Rec114 C-terminal domain was amplified from pDD003 using dd027 and dd028 and cloned into pETDuet-1 vector by Gibson assembly to yield pDD006. N-terminal ^HisSUMO^Mei4 was amplified from pDD004 using primers dd025 and dd026 and cloned into pDD006 vector by Gibson assembly to yield pDD009.

Expression vectors for minimal trimeric Rec114−Mei4 mutant complexes were obtained using pCCB825 as a template by inverse PCR and self-ligation using dd084 and dd085, dd095 and dd114 to yield, respectively, pDD044 (^HisSUMO^Rec114-K405A, Mei4-WT) and pDD045 (^HisSUMO^Rec114-WT, Mei4-E16A/D18A). Plasmid pDD051(^HisSUMO^Rec114-K405A/E419A, Mei4-WT) was obtained using pDD044 as a template and dd080 and dd118 as primers.

Expression vectors for the ^HisFlag^Rec114-6KR (pDD100) and ^HisFlag^Rec114-8KR (pDD101) mutants were obtained by inverse PCR and self-ligation with dd172 and dd173, dd174 and dd175 as primers respectively and pCCB848 as a template.

Expression vector for the DNA-binding mutant minimal trimeric complex (Rec114-6KR, ^HisSUMO^Mei4) was obtained by PCR amplification of pDD100 with primers dd27 and dd28, and pDD009 with primers cb1522 and cb1523 followed by Gibson assembly to yield pCCB1001. Expression vectors for fusion Rec114 constructs were generated by PCR amplification of Rec114 fragment from pDD009 with primers cb1524 and dd0028, and the pDD009 vector with primers cb1525 and cb1523, followed by Gibson assembly to yield pCCB1002. The expression vector with the 6KR mutation in the C-terminal subunit (pCCB1003) was generated using the same strategy using pDD100 as a template to amplify the Rec114-6KR fragment. The expression vector with the 6KR mutation in the N-terminal subunit (pCCB1004) and in both subunits (pCCB1005) were generated following the same procedure using templates pDD009 and pCCB1001, and pDD100 and pCCB1001, respectively.

Expression vectors for Mer2 truncations were obtained by inverse PCR and self-ligation using pCCB750 as a template to generate pCCB973 (^HisSUMO^Mer2(41-314)), pCCB975 (^HisSUMO^Mer2(161-314)) and pCCB978 (^HisSUMO^Mer2(1-110)) using cb1346 and cb1497, cb1346 and cb1495, and cb1342 and cb1492 primers, respectively. Plasmids pCCB973, pCCB978 and pCCB975 were further used as templates to generate pCCB981 (^HisSUMO^Mer2(41-224)), pCCB979 (^HisSUMO^Mer2(41-110)) and pCCB980 (^HisSUMO^Mer2(161-224)) using primers cb1342 and dd121, cb1346 and cb1497, and cb1342 and dd121, respectively. Plasmid pDD078 (^HisSUMO^Mer2(138-224)) and pDD079 (^HisSUMO^Mer2(41-136)) were obtained by using, respectively, cb1346 and dd146, or cb1342 and dd147 as primers and pCCB981 as a template. The expression vector for the ^HisSUMO^Mer2-KKTK mutant (pDD015) was obtained by inverse PCR and self-ligation by using dd060 and dd067 as primers and pCCB750 as a template.

The sequence coding for the coiled-coil domains of PRD3, Rec15, ASY2 and PAIR1 were synthetized as gBlocks and cloned by Gibson assembly in pSMT3 vector to yield HisSUMO fusion constructs pCCB990, pCCB991, pCCB992 and pCCB993 respectively. The expression vector for the ^HisSUMO^Rec15(coiled-coil)-5KR mutant (pDD086) was obtained by inverse PCR and self-ligation with dd157 and dd158 as primers and pCCB991 as a template.

Expression vectors for *M. musculus* REC114, MEI4 and IHO1 were generated by PCR-amplification of mouse testes cDNA (using primers cb1315 and cb1316, cb1317 and cb1318, and cb1322 and cb1327, respectively), and cloned by Gibson assembly into vectors pFastBac1-Flag (REC114), pFastBac1-MBP (MEI4), or pSMT3 (IHO1) to yield pCCB805, pCCB806, and pCCB808. The coiled-coil domain of IHO1 was amplified from pCCB808 using primers cb1498 and cb1499 and cloned into pSMT3 by Gibson assembly to yield pCCB982. The expression vector for the ^HisSUMO^IHO1(coiled-coil)-KKRR mutant (pDD081) was obtained by inverse PCR and self-ligation with dd150 and dd151 as primers and pCCB982 as a template.

The REC114 C-terminal domain and the N-terminal Mei4 were amplified from pCCB805 and pCCB806 using cb1507 and cb1503, cb1505 and cb1508 respectively and cloned into pETDuet-1 vector by Gibson assembly to yield pCCB984. Minimal ^HisSUMO^REC114-F230A, ^MBP^MEI4 mutant was generated by QuikChange mutagenesis using cb1509 and cb1510 as primers and pCCB984 as a template to yield pCCB987. Minimal ^HisSUMO^REC114-F240A, ^MBP^MEI4 and minimal ^HisSUMO^REC114-F230A/F240A/F243A, ^MBP^MEI4 mutants were generated by inverse PCR and self-ligation using pCCB984 and pCCB987 as template and dd129 and dd130, dd154 and dd167 as primers to yield pCCB988 and pDD082 respectively. Sequences coding for *S. pombe* Rec7(289-339) and Rec24(1-50), *A. thaliana* PHS1(260-310) and PRD2(1-50) and *Z. mays* PHS1(297-347) and MPS1(1-87) were ordered as gBlocks and cloned into a pETDuet-1-derived vector to yield dual HisSUMO and MBP-tagged co-expression vectors pDD085, pDD093 and pDD095, respectively. Co-expression vectors for minimal ^HisSUMO^Rec7-F325A−^MBP^Rec24, ^HisSUMO^PSH1-F290A−^MBP^PRD2, and ^HisSUMO^PSH1-F327A−^MBP^MPS1 mutants were generated by reverse PCR and self-ligation using dd169 and dd170, edj20 and edj21, edj24 and edj25 as primers and pDD085, pDD093 and pDD095 as templates to yield pDD091, pEDJ11 and pEDJ12 respectively. Co-expression vectors for minimal ^HisSUMO^Rec7-Y319A/F325A/L328A−^MBP^Rec24, ^HisSUMO^PSH1-Y284A/F290A/L294A−^MBP^PRD2, and ^HisSUMO^PSH1-F327A/L334A/I338A−^MBP^MPS1 mutants were generated by reverse PCR and self-ligation using edj18 and edj19, edj22 and edj23, edj24 and edj26 as primers and pDD091, pEDJ11 and pEDJ12 as templates to yield pEDJ10, pEDJ13 and pEDJ14 respectively.

### Expression and purification of recombinant proteins

Viruses were produced using a Bac-to-Bac Baculovirus Expression System (Invitrogen) according to the manufacturer’s instructions. We infected 2×10^9^ *Spodoptera frugiperda* Sf9 cells (Gibco, Thermo Fisher) with combinations of viruses at a multiplicity of infection (MOI) of 2.5 each. Expression of ^HisFlag-TEV^Rec114, ^MBP-TEV^Mei4 used viruses generated from pCCB789 and pCCB791, ^HisFlag-TEV^Rec114 4KR, ^MBP-TEV^Mei4, used viruses generated from pCCB848 and pCCB791, ^HisFlag-TEV^Rec114 6KR, ^MBP-TEV^Mei4, used viruses generated from pDD100 and pCCB791, ^HisFlag-TEV^Rec114 8KR, ^MBP-TEV^Mei4, used viruses generated from pDD101 and pCCB791, and mouse ^HisFlag-TEV^REC114, ^MBP-TEV^MEI4 used viruses generated from pCCB805 and pCCB806. After 72h infection, cells were collected, washed with phosphate buffer saline (PBS), frozen in dry ice and kept at -80 °C until use. All purification steps were carried out at 0-4 °C. Cell pellets were resuspended in 3 volumes of lysis buffer (50 mM HEPES-NaOH pH 7.5, 0.17 mM DTT, 65 mM imidazole, 4.6 μM leupeptin, 0.3 μM aprotinin, 3.3 μM antipain, 2.9 μM pepstatin, 3.3 μM chymostatin and 1 mM phenylmethanesulfonyl fluoride (PMSF)) and then pooled in a beaker and mixed slowly with a stir bar for 20 minutes. 10% of ice-cold glycerol and 335 mM NaCl were added to the cell lysate that was then centrifuged at 43,000 g for 30 min. The cleared extract was loaded onto 1 ml pre-equilibrated Ni-NTA resin (Thermo Scientific). The column was washed extensively with nickel buffer (25 mM HEPES-NaOH pH 7.5, 500 mM NaCl, 10% glycerol, 0.1 mM DTT, 20 mM imidazole, 0.1 mM PMSF). The tagged complexes were then eluted in nickel buffer containing 500 mM imidazole. The complexes were further purified on amylose resin (NEB). Fractions containing protein were pooled and diluted in 3 volumes of amylose buffer (25 mM HEPES-NaOH pH 7.5, 500 mM NaCl, 10% glycerol, 2 mM DTT, 5 mM EDTA). Next, the complexes were bound to 1 ml of the amylose resin in a disposable chromatography column (Thermo Scientific) and the resin was washed extensively. Complexes were eluted from amylose resin with buffer containing 10 mM maltose. Fractions containing protein were concentrated in 50-kDa cutoff Amicon centrifugal filters (Millipore). Aliquots were flash frozen in liquid nitrogen and stored at −80 °C.

For expression of recombinant proteins in E. coli, expression vectors were transformed in BL21 DE3 cells and plated on LB plates containing the appropriate antibiotic. For the expression of IHO1, the expression vector was transformed in *E. coli* C41 cells. Cells were then cultured in liquid medium at 37 °C to an optical density (OD_600_) of 0.6. Expression was carried out at 30 °C for 3 hours with 1 mM isopropyl β-D-1-thiogalactopyranoside (IPTG). Cells were resuspended in nickel buffer (25 mM HEPES-NaOH pH 7.5, 500 mM NaCl, 10% glycerol, 0.1 mM DTT, 20 mM imidazole, 0.1 mM PMSF), frozen dropwise in liquid nitrogen and kept at −80 °C until use. All the purification steps were carried out at 0–4 °C. Cells were lysed by sonication and centrifuged at 30-43,000 g for 30 minutes. The cleared extract was loaded onto 1 ml pre-equilibrated Ni-NTA resin (Thermo Scientific). The column was washed extensively with nickel buffer then eluted in buffer containing 500 mM imidazole. The 6His–SUMO tag was cleaved with Ulp1 during overnight dialysis in nickel buffer. The sample was then loaded on a second nickel column to remove 6His–SUMO and Ulp1. The flow-through was then loaded on a Superose 6 Increase 10/300 GL column preequilibrated with gel filtration buffer (25 mM HEPES-NaOH pH 7.5, 300 mM NaCl, 10% glycerol, 1 mM DTT, 5 mM EDTA). After gel filtration, fractions containing protein were concentrated in 10-kDa cutoff Amicon centrifugal filters (Millipore). Aliquots were flash frozen in liquid nitrogen and stored at −80 °C.

For the production of the doubly-labeled U-[^13^C,^15^N] protein, the minimal medium contained M9 salts (6.8 g/L Na_2_HPO_4_, 3 g/L KH_2_PO_4_, and 1 g/L NaCl), 2 mM MgSO_4_, 0.2 mM CaCl_2_, trace elements (60 mg/L FeSO_4_·7H_2_O, 12 mg/L MnCl_2_·4H_2_O, 8 mg/L CoCl_2_·6H_2_O, 7 mg/L ZnSO_4_·7H_2_O, 3 mg/L CuCl_2_·2H_2_O, 0.2 mg/L H_3_BO_3_, and 50 mg/L EDTA), BME vitamin mix (Sigma), and 1 g/L ^15^NH_4_Cl and 2 g/L [^13^C_6_]glucose (CortecNet) as the sole nitrogen and carbon sources, respectively. Expression was carried out at 30 °C for 6 h with 1 mM IPTG. After affinity chromatography on Ni-NTA resin, the sample was loaded to Superdex 75 Increase 10/300 GL column. Proteins were eluted in a buffer containing 20 mM sodium phosphate (NaP) and 30 mM NaCl (pH 6.0).

### Nuclear Magnetic Resonance (NMR) spectroscopy

The samples contained 0.8 mM of U-[^13^C,^15^N] Rec114 dimer in 20 mM sodium phosphate 20 mM NaCl pH 6.0, 0.02% NaN_3_ and 10% D_2_O or 1.4 mM of U-[^13^C,^15^N] 2:1 Rec114−Mei4 complex in 20 mM sodium phosphate 30 mM NaCl pH 6.0, 5 mM DTT, 0.02 % NaN_3_ and 10% D_2_O. All NMR spectra were acquired at 298 °K on a Bruker Avance III HD 800 MHz spectrometer, equipped with a TCI cryoprobe. The experimental set comprised 2D [^1^H,^15^N] HSQC, [^1^H,^13^C] HSQC, and constant-time [^1^H,^13^C] HSQC for the aromatic region; 3D ^15^N edited NOESY-HSQC and ^13^C-edited NOESY-HSQC for aliphatic and aromatic regions (all recorded with the mixing time of 120 ms); and triple-resonance BEST-HNCACB, BESTHN(CO)CACB, BEST-HNCO, BEST-HN(CA)CO, HBHA(CO)NH, (H)CCH-TOCSY, and H(C)CH-TOCSY spectra. The NMR data were processed in TopSpin 3.6 (Bruker) or NMRPipe (Delaglio et al. 1995), and analyzed in CCPNMR (Vranken et al. 2005). Semi-automatic assignment of the protein backbone was performed in CCPNMR (Vranken et al. 2005). The assignments of N, NH, Hα, Hβ, CO, Cα, and Cβ atoms were obtained from the identification of intra- and inter-residue connectivities in HNCACB, HN(CO)CACB, HNCO, HN(CA)CO, and HBHA(CO)HN experiments at the ^1^H,^15^N frequencies of every peak in the [^1^H,^15^N] HSQC spectrum. Assignments were extended to the side chain signals using correlations within (H)CCH-TOCSY and H(C)CH-TOCSY experiments. Aromatic ^1^H and ^13^C assignments were obtained from constant-time [^1^H,^13^C] HSQC and ^13^C-edited NOESY-HSQC spectra focused on the aromatic region. Remaining aliphatic and aromatic side-chains were assigned from 3D ^15^N- and ^13^C-edited NOESY-HSQC spectra. The ^1^H, ^13^C and ^15^N chemical shifts for the 2:1 Rec114−Mei4 complex were deposited in the Biological Magnetic Resonance Bank under the accession number 26335.

The average chemical shift difference (Δδ_avg_) reported in **Supplemental Fig. S2C** was calculated as Δδ_avg_ = (Δδ_N_^2^/50 + Δδ_H_^2^/2)^0.5^, where Δδ_N_ and Δδ_H_ are the chemical shift differences of the backbone amide nitrogen and proton, respectively, for the double HSQC resonances of a given Rec114 residue, The secondary structure of Rec114 and Mei4 in their 2:1 complex (**Supplemental Fig. S2B**) was predicted from the backbone chemical shifts using the chemical shift index function and the DANGLE module (Cheung et al. 2010) in CCPNMR (Vranken et al. 2005).

### Thermal shift assays

Aggregation (T_agg_) and melting (T_m_) temperatures were obtained from analysis of the static light scattering (SLS) and intrinsic tryptophan fluorescence, respectively, measured simultaneously in UNcle UNI (Unchained Labs, CA, USA). Series of samples containing 2-3 mg/ml of ternary Rec114−Mei4 complexes or Rec114 dimer in 25 mM HEPES-NaOH pH 7.5, 10% Glycerol, 300 mM NaCl, 2 mM DTT, 5 mM EDTA were prepared in triplicate and loaded into the UNcle quartz cells (9 μl per cell). The tryptophan fluorescence spectra and SLS at 266 or 434 nm were measured during a linear temperature gradient of 1 °C/min from 20 to 95 °C. To maximize the frequency of measurements, a holding time was not used. The T_agg_ and T_m_ values – defined as the inflection points of the corresponding thermal curves – were obtained, respectively, from the analysis of the SLS absorption at 266 nm or the barycentric mean given by the equation 1:

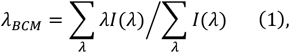

where λ and I(λ) are the wavelength and the corresponding intensity in the fluorescence spectrum, while the summation covers the 300-430 nm region. The thermal curves were analyzed with a two-state transition model given by the equation 2 (van Nuland et al. 1998):

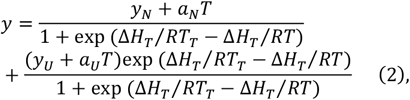

where y is the signal observed at temperature T, R is the absolute gas constant, y_N_ + a_N_T and y_U_ + a_U_T are the linear slopes of the pre- and post-transitional regions of the thermal curves, respectively, and ΔH_T_ is the change in enthalpy at the transition temperature T_T_. Thus, T_T_ values obtained from the non-linear fit of the SLS and tryptophan fluorescence thermal scan curves correspond to T_agg_ and T_m_, respectively. The obtained values agreed well with those determined by the UNcle Analysis software.

### DNA substrates and gel-shift assays

Short double-strand DNA substrates were generated by annealing complementary oligos. The substrates were the following (with oligo names in parentheses): dsDNA (cb95 and cb100), HJ20 (cb922, cb923, cb924 and cb925), HJ40 (cb095, cb096, cb097, cb098), 3’-Flaps (cb095, cb098 and cb122), Fork (cb095, cb098, cb122 and cb120), Y (cb095, cb098 and cb101). The 40-nt and 80-nt oligos were first purified on 10% polyacrylamide-urea gels. Oligos were subsequently mixed in equimolar concentrations (10 μM) in STE (100 mM NaCl, 10 mM Tris-HCl pH 8, 1 mM EDTA), heated and slowly cooled on a PCR thermocycler (98 °C for 3 min, 75 °C for 1 h, 65 °C for 1 h, 37 °C for 30 min, 25 °C for 10 min). For radioactive labeling, 1/20^th^ of the annealed substrates was 5′-end-labelled with [γ-^32^P]-ATP and T4 polynucleotide kinase (New England Biolabs). For fluorescently labeled HJ40 substrates, oligo cb095 was replaced by 5’-6FAM modified version (dd077). Labeled and unlabeled substrates were purified by native polyacrylamide gel electrophoresis.

Binding reactions (20 μl) were carried out in 25 mM Tris-HCl pH 7.5, 7.5% glycerol, 100 mM NaCl, 2 mM DTT, 1 mM EDTA, and 1 mg/ml BSA. Unless stated otherwise, reactions contained 1 nM radiolabeled substrate or 10 nM fluorescently labeled substrate and the indicated concentration of protein. For Mer2 and IHO1 complexes, concentrations are expressed as monomers. For Rec114−Mei4 complexes, concentrations are expressed as 2:1 heterotrimers. Complexes were assembled for 30 minutes at 30 °C and separated by gel electrophoresis. Binding reactions were separated on 5% TAE-polyacrylamide gels at 150 V for 2 h 30 min, and fluorescent gels were visualized using a Typhoon scanner (Cytiva), while radioactive gels were dried and imaged by autoradiography.

### Filter binding Assays

Binding reactions (20 μl) were carried out in 25 mM Tris-HCl pH 7.5, 7.5% glycerol, 100 mM NaCl, 2 mM DTT, 1 mM EDTA, and 1 mg/ml BSA. Reactions contained 10 nM fluorescently labeled HJ40 and the indicated concentration of unlabeled DNA competitors and protein. Complexes were assembled for 30 minutes at 30 °C then passed through a nitrocellulose membrane (Amersham Protran 0.45 μm NC) to retain protein and protein-bound DNA using a 96-well Bio-Dot apparatus (Bio-Rad). Each well was then washed twice with 50 μl buffer containing 25 mM Tris-HCL pH 7.5, 100 mM NaCl and 1 mM DTT. Fluorescent signal was visualized using a Typhoon scanner (Cytiva) and quantified using ImageJ.

### In vitro condensation assay

Rec114–Mei4 complexes were first diluted to 5 μl in storage buffer adjusted to a final salt concentration of 360 mM NaCl. After 5 min at room temperature, condensation was induced by threefold dilution in reaction buffer containing DNA and no salt, to reach final 15-μl reactions that contained 25 mM Tris-HCl pH 7.5, 5% glycerol, 120 mM NaCl, 2 mM DTT, 1 mg/ml BSA, 5 mM MgCl_2_, 5% PEG 8000. A typical binding reaction contained 150 ng supercoiled pUC19 (5.7 nM) and 4 μM Alexa488-labeled Rec114^(375-428)^–Mei4^(1-43)^. After 30 minutes incubation at 30 °C with occasional mixing, 4 μl was dropped on a microscope slide and covered with a coverslip. Images were captured on Zeiss Axio observer with a 100×/1.4 NA oil immersion objective. Images were analyzed with ImageJ using a custom-made script. In brief, a fixed threshold was applied to a 129.24 × 129.24-μm (2048 × 2048-pixel) images. The intensity inside the foci mask was integrated. Data points represent averages of 5–16 images per sample. Data were analyzed using Graphpad Prism 9.

### AFM imaging

For AFM imaging of Mer2 and its orthologs bound to plasmid DNA, protein complexes were diluted to the indicated concentration in the presence of 1 nM supercoiled pUC19 in 25 mM HEPES-NaOH pH 6.8, 5 mM MgCl_2_, 50 mM NaCl, 10% glycerol. Complexes were assembled at 30 °C for 30 minutes. A volume of 40 μl of the protein-DNA binding reaction was deposited for 2 minutes onto freshly cleaved mica treated with aminopropyl silane (APTES). The sample was rinsed with 10 ml ultrapure deionized water and the surface was dried using a stream of nitrogen. AFM images were captured using a Multimode 8 nanoscope (Bruker AXS Corporation, Santa Barbara, CA) in tapping mode at room temperature. Scan Asyst air cantilevers (Bruker AXS Corporation) with a spring constant of 0.2–0.4 N/m was used for imaging. Images were collected at a speed of 0.5–1 Hz with an image size of 1 μm at 256 × 256 pixels resolution. Data was analyzed using the Nanoscope Analysis software (Bruker AXS Corporation).

### Pulldown assay

Tagged wild-type and mutants minimal Rec114−Mei4 orthologs were expressed in 50 ml *E. coli* BL21 cultures and purified by affinity chromatography on Nickel resin following a similar procedure as described above. Cells were lysed by sonication and centrifuged at 15000 rpm for 20 minutes. The cleared extract was loaded onto 100 μl of pre-equilibrated Ni-NTA resin (Thermo Scientific) in nickel buffer containing 25 mM Hepes pH 7.5, 300 mM NaCl, 10% glycerol, 0.1 mM DTT, 40 mM imidazole, and 0.5% Triton. After 30 min of incubation on a rotating wheel at 4 °C, the resin was washed five times with 1 ml nickel buffer, then eluted in a buffer containing 500 mM imidazole. Proteins were resuspended in Laemmli buffer and analyzed by SDS–PAGE.

### SEC-MALS

Light scattering data were collected using a Superdex 200 increase 10/300 GL Size Exclusion Chromatography (SEC) column connected to a AKTA Pure Chromatography System (Cytiva). The elution from SEC was monitored by a differential refractometer (Optilab, Wyatt), and a static and dynamic, multiangle laser light scattering (LS) detector (miniDAWN, Wyatt). The SEC–UV/LS/RI system was equilibrated in buffer 25 mM HEPES-NaOH pH 7.5, 500 mM NaCl, 10% glycerol, 2 mM EDTA at a flow rate of 0.3 ml/min. The weight average molecular masses were determined across the entire elution profile at intervals of 0.5 s from static LS measurement using ASTRA software.

### Structural refinement of the Rec114−Mei4 complex

All simulations were performed in Xplor-NIH v 2.49 (Schwieters et al. 2003), starting from the AlphaFold models of the minimal Rec114−Mei4 complex and the full-length Mei4 obtained in this work. The intermolecular XL-MS data were converted into pairwise distance restraints between lysine sidechains as described elsewhere (Gong et al. 2020). In all refinement runs, the position of the core Rec114−Mei4 minimal complex (residues 399-426 of Rec114 and 16-42 of Mei4) was kept fixed; Mei4 globular domain (residues 65-401) treated as a rigid body group; while the intervening linkers, N- and C-terminal tails of both proteins, and sidechains of crosslinked lysines given full torsional degree of freedom. The computational protocol comprised an initial simulated annealing step followed by the side-chain energy minimization. The total minimized energy function consisted of the standard geometric (bonds, angles, dihedrals, and impropers) and steric (van der Waals) terms, a knowledge-based dihedral angle potential (Schwieters et al. 2003) and the experimental XL-MS restraints term (Gong et al. 2020). In each refinement run, 100 structures were calculated and 10 lowest-energy solutions – representing the best agreement with the experimental data – retained for the subsequent analysis. To model the DNA-bound Rec114−Mei4 complex, we generated the double helical structure of the canonical Watson-Crick paired B-DNA (20-bp oligomer 5’-GAGATGTCCATGGACATCTC-3’), and docked two copies of the DNA duplex to the best structure of the XL-MS refined Rec114−Mei4 complex. The model was obtained by minimizing the distance between the DNA-binding Rec114 residues R395, K396, K399, and R400 and the central grooves of the double-stranded DNA oligomer, while avoiding steric clashes between protein sidechains and the DNA. The resulting model shows the DNA-bound Rec114−Mei4 complex, where Rec114 residues R395, K396, K399, and R400 make a number of intermolecular contacts with the DNA phosphate backbone and nucleotide bases.

### Twister analysis of predicted coiled-coil structures

AlphaFold predicted coiled-coil structures were analyzed using Twister (Strelkov and Burkhard 2002). A coiled coil can be described by the radius and pitch (i.e., distance along the axis that corresponds to a full turn) of the superhelix, and the radius and pitch of the α-helices. In addition, the pitch of the coiled coil and the α-helices can be expressed per residue (phase yield). Analyses of the coiled-coil and the α-helical phase yields reveal whether the coiled coil and/or the individual helices are locally distorted. Analyses of coiled-coil left and right pitch parameters reveal whether the normally left-handed coiled coil locally switches to a right-handed superhelix. Finally, Crick’s angle describes the position of a specific residue relative to the axis of the coiled coil. Variations in Crick’s angles between residues that occupy equivalent positions (e.g., a or d) further indicates geometrical distortions of the coiled coil.

### SEC-SAXS

All experiments were performed at the SOLEIL BioSAXS beamline SWING (Gif-sur-Yvette, France). SEC-SAXS data were collected in HPLC mode using a Shodex KW404-4F column pre-equilibrated with SAXS buffer (25 mM HEPES-NaOH, 500 mM NaCl, 5 mM EDTA, 5% glycerol, 1 mM TCEP, pH 7.5). Samples were concentrated on-site to approximately 5-10 mg/ml using a 10-kDa cut-off centrifugal filter (Amicon). 90 μL samples were injected and eluted at a flow rate of 0.2 ml/min while scattering data was collected with an exposure time of 990 ms and a dead time of 10 ms. The scattering of pure water was used to calibrate the intensity to absolute units. Data reduction was performed using FoxTrot. Data were processed using BioXTas RAW (Hopkins et al. 2017) and analyzed using RAW and the ATSAS package (Manalastas-Cantos et al. 2021). The information on data collection and derived structural parameters is summarized in **Supplemental Table S7**. *Ab initio* calculations were performed with the DENSS suite (Grant 2018) and four-fold symmetry was imposed during shape reconstruction.

Molecular models of the parallel and antiparallel coiled-coils fused to SUMO domains were generated from the AlphaFold models obtained in this work and the X-ray structure of the SUMO protein (PDB 7P47) (Varejão et al. 2021) and refined in Xplor-NIH v 2.49 (Schwieters et al. 2003). In all refinement runs, the positions of the coiled-coil domains were kept fixed, SUMO globular domains treated as rigid body groups, while the intervening linkers and terminal tails given full torsional degree of freedom. The computational protocol comprised an initial simulated annealing step followed by the side-chain energy minimization. The total minimized energy function consisted of the standard geometric (bonds, angles, dihedrals, and impropers) and steric (van der Waals) terms, and a knowledge-based dihedral angle potential (Schwieters et al. 2003). In each refinement run, 100 structures were calculated and 10 lowest-energy solutions retained. The parallel coiled-coil SUMO fusion constructs were further refined in Xplor-NIH against the experimental SAXS data. In addition to the aforementioned energy terms, the energy function included a SAXS energy term incorporating the experimental data (Schwieters and Clore 2014). The agreement between the experimental and calculated SAXS curves (obtained with the calcSAXS helper program, which is part of the Xplor-NIH package) was assessed by calculating the χ^2^:

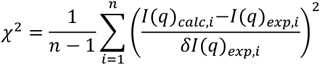

where *I*(*q*)_)*caic,i*_ and *I*(*q*)_*exp,i*_ are the scattering intensities at a given q for the calculated and experimental SAXS curves, *δI*(*q*)_*exp,i*_ is an experimental error on the corresponding *I*(*q*)_*exp,i*_ value, and n is the number of data points defining the experimental SAXS curve. The models were fitted into the *ab initio* densities with UCSF ChimeraX (Goddard et al. 2018).

### Yeast targeting vectors, strains construction and spore viability assay

Yeast strains were from the *SK1* background. All strains used in this study are listed in **Supplemental Table S6**.

To produce a yeast targeting vector for *MER2*, the *HphMX6* selection cassette was first amplified using primers mh001 and mh002 and inserted 50 bp downstream of the *MER2* locus by transformation in CBY006. The *MER2::HphMX* sequence was then amplified from genomic DNA with primers mh003 and mh004 and cloned into a TOPO vector by TOPO Blunt cloning, yielding pMH002. The Mer2(K265A, R266A, R267A, R268A) (KRRR) mutation and Mer2(K136A, K137A, K139A) (KKTK) mutation were generated by QuikChange mutagenesis using primers cb1186 and cb1187, dd067 and mh036 with pMH002 as a template to yield pMH026 and pMH030, respectively. The *Mer2-KRRR::HphMX* and *mer2-KKTK::HphMX* sequences were inserted at the *MER2* locus following digestion of pMH026 and pMH030 with SpeI and NotI and transformation into CBY006 to yield strains CBY612 and CBY614, respectively. Strains were genotyped by PCR and sequencing.

The *REC114::HphMX* sequence was amplified from genomic DNA of CBY388 with primers mh008 and mh009, and cloned into a TOPO vector by TOPO Blunt cloning, yielding pCCB929. The *rec114-R395A/K396A/K399A/K400A* (4KR) mutant was generated by inverse PCR and self-ligation of pCCB929 with primers cb1332 and cb1334 to yield pMH029. The *rec114-R395A/K396A/K399A/R400A/K403A/K407A* (6KR) and *rec114-R395A/K396A/K399A/R400A/K403A/K407A/K417A/K424A* (8KR) mutants were generated by inverse PCR and self-ligation of pMH029 with primers dd172 and dd176, dd174 and dd177 to yield pDD104 and pDD105 respectively. The *rec114-6KR::HphMX, rec114-8KR::HphMX* and *rec114-4KR::HphMX* cassettes were inserted at the *REC114* locus following digestion of pDD104, pDD105 and pMH029 with XhoI and HindIII and transformation into CBY006 to yield strains CBY718, CBY720 and CBY724, respectively. Strains were genotyped by PCR and sequencing.

To measure spore viability, diploid strains were induced to undergo sporulation in 2% potassium acetate for 2 days, followed by tetra dissection and grown on YPD plates.

## End Matter

### Author Contributions and Notes

D.D. and C.C.B designed research; D.D. carried out all experiments except as noted; E.D.J. performed pulldown analyses in Fig. 6B and Supplemental Fig. S11, gel-shift analyses in Fig. 7G and Supplemental Figs. S12 and S18A, C, and assisted D.D. with plasmid constructions and protein purifications; P.L. provided technical assistance; K.M. performed AFM imaging experiments, M.H. generated and analyzed yeast mutants; S.C. and Y.G.-J.S. performed and analyzed SAXS experiments; A.N.V. performed NMR analyses, thermal-shift assays, generated AlphaFold models and performed structural modeling. D.D., A.N.V. and C.C.B wrote the paper with input from all authors. C.C.B. supervised the research and secured funding.

The authors declare no conflict of interest.

## Acknowledgments

We thank members of the CCB lab for discussions and comments on the manuscript, particularly Cédric Oger and David Álvarez Melo. We thank Sylvie Derclaye (MICA core facility, UCLouvain) for help with AFM experiments and Joseph Nader (FYMO, UCLouvain) for technical advice. We thank Gholamreza Hassanzadeh Ghassabeh and Ema Romão of the VIB Nanobody Core Facility for providing access to the UNcle setup and help with the measurements, and Javier Perez and Aurélien Thureau of the SWING beam line at SOLEIL synchrotron for outstanding support. This work was supported by the European Research Council under the European Union’s Horizon 2020 research and innovation program (ERC grant agreement 802525 to CCB), and the Fonds National de la Recherche Scientifique (MIS-Ulysse grant F.6002.20 to CCB). DD and KM are funded by FNRS Aspirant fellowships (projects FC36183 and FC42859) and MH by a FRIA fellowship (project FC45991). SC is funded by the FWO-Vlaanderen (project G017221N). CCB is a FNRS Research Associate.

