## Supplemental Files for "Evolutionary conservation of the structure and function of meiotic Rec114−Mei4 and Mer2 complexes"

### Supplemental Figures

- Supplemental Figure S1: Quality assessment of the Rec114–Mei4 AlphaFold model.
- Supplemental Figure S2: NMR validation of minimal trimeric Rec114–Mei4 complex.
- Supplemental Figure S3: Modeling of the C-terminus of Rec114 in complex with full-length Mei4.
- Supplemental Figure S4: DNA-binding properties of the Rec114–Mei4 complex.
- Supplemental Figure S5: Quality assessment of Mer2 AlphaFold models.
- Supplemental Figure S6: SEC-MALS analyses of Mer2 truncations.
- Supplemental Figure S7: DNA-binding properties of Mer2.
- Supplemental Figure S8: Conservation of Rec114 and Mei4 structures.
- Supplemental Figure S9: Analysis of the HEAT-repeat structure of Mei4 orthologs.
- Supplemental Figure S10: Quality assessment of AlphaFold models of Rec114–Mei4 orthologs.
- Supplemental Figure S11: Interaction between Rec114–Mei4 orthologs.
- Supplemental Figure S12: DNA-binding properties of Rec114–Mei4 orthologs.
- Supplemental Figure S13: Quality assessment of AlphaFold models of Mer2 orthologs.
- Supplemental Figure S14: Twister analysis of Mer2 and IHO1 coiled coils.
- Supplemental Figure S15: Twister analysis of PRD3 and Rec15 coiled coils.
- Supplemental Figure S16: Twister analysis of ASY2 and PAIR1 coiled coils.
- Supplemental Figure S17: SAXS analysis of Mer2 orthologs.
- Supplemental Figure S18: DNA-binding properties of Mer2 orthologs.
- Supplemental Figure S19: Model of Mer2 condensate assembly.

### Supplemental Tables

- Supplemental Table S1: Nuclear Overhauser effect spectra analyses.
- Supplemental Table S2: Violation analysis of Rec114–Mei4 crosslinks.
- Supplemental Table S3: List of oligonucleotides.
- Supplemental Table S4: Sequence of synthetic DNA fragments.
- Supplemental Table S5: List of plasmids.
- Supplemental Table S6: List of yeast strains.
- Supplemental Table S7: SAXS data collection and scattering-derived parameters.

### Structural models (pdb files)

- AlphaFold models of 2:1 *S. cerevisiae* Rec114–Mei4 complexes and orthologs in *M. musculus, S. pombe, A. thaliana, Z. mays*.
- AlphaFold models of tetrameric *S. cerevisiae* Mer2 complexes and orthologs in *M. musculus*, *S. pombe*, *S. macrospora*, *A. thaliana*, *Z. mays*.

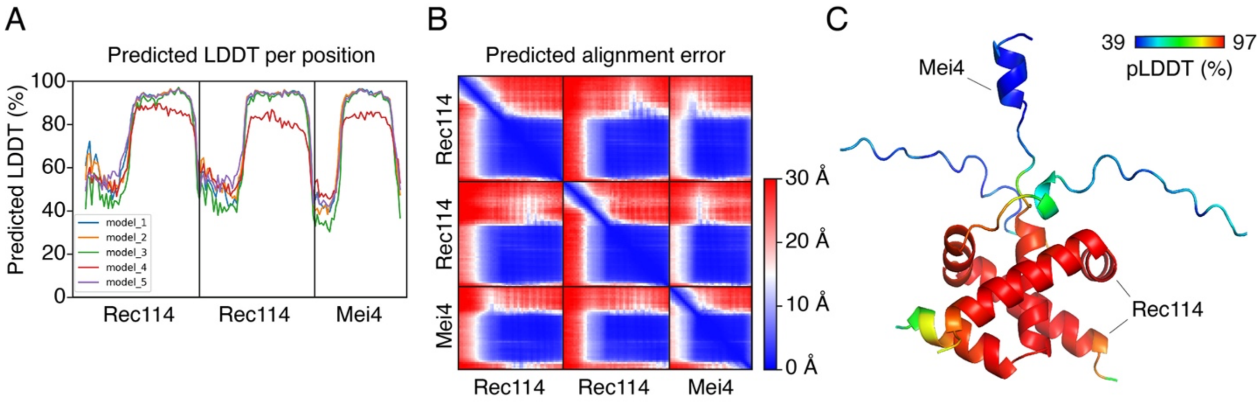

**Supplemental Figure S1. Quality assessment of the Rec114–Mei4 AlphaFold model.**
(A) Predicted local distance difference test (pLDDT) of the five best AlphaFold models of the Rec114–Mei4 complex. Large values signify that the local geometry of the folded protein regions is predicted with high confidence. All five models show similar structures, except for model 4. (B) Predicted alignment error of the highest-scoring AlphaFold Rec114–Mei4 model (model 1). Low values indicate the high confidence in the relative orientations of the interacting domains. (C) AlphaFold model of the minimal trimeric Rec114–Mei4 complex (model 1) color-coded by the confidence score. The Rec114–Mei4 interaction domain is predicted with high confidence.

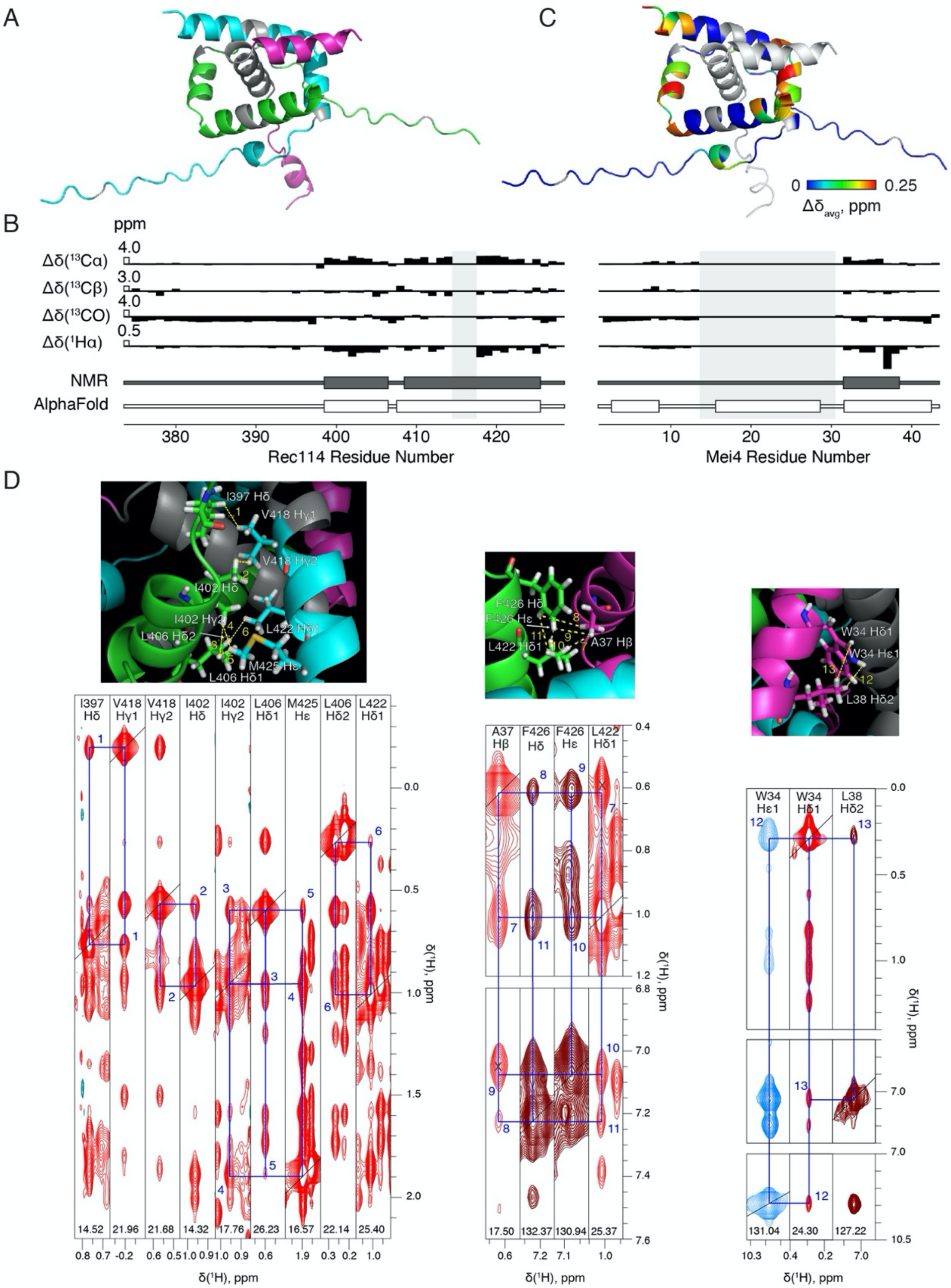

**Supplemental Figure S2. NMR validation of the Rec114–Mei4 complex predicted by AlphaFold.**
(A) Structure of the complex predicted with AlphaFold. Two Rec114 chains are in green and cyan, Mei4 is in magenta. The residues for which no backbone amide resonances were observed in the HSQC spectrum are in grey. Mei4 residues 14-30 and Rec114 residues 216-218 exhibited very few, if any, NMR signals and could not be unambiguously assigned. This, combined with a strong peak overlap (particularly in the [$^{1}$H,$^{13}$C] HSQC spectrum), weak signals and few cross-peaks in NOESY spectra, precluded *de novo* structure determination of the complex by NMR spectroscopy. (B) Threshold deviations from random-coil Cα, Cβ, CO, and Hα chemical shifts (Δδ), with the open bars indicating the Δδ size for each nucleus type. The α-helices predicted from the NMR chemical shift index analysis or the AlphaFold model are shown as grey and white rectangles, respectively. The grey shades delineate protein regions for which no NMR signals were observed. (C) AlphaFold model with Rec114 residues colored by the difference in chemical shifts ($\Delta\delta_{avg}$) of their double backbone amide resonances (bold labels in **Fig. 1E**). Mei4 is in grey. (D) Close contacts between protein side chains in the AlphaFold model that give rise to observed NOEs. Color code as in panel A. Strips of 3D $^{13}$C-edited aliphatic NOESY (red), $^{13}$C-edited aromatic NOESY (brown), and $^{15}$N-edited NOESY (blue) spectra showing protons of interest. The NOE cross-peaks are identified by blue rectangles. Black crosses denote additional, overlapping signals. The numbering corresponds to that in **Supplemental Table S1**.

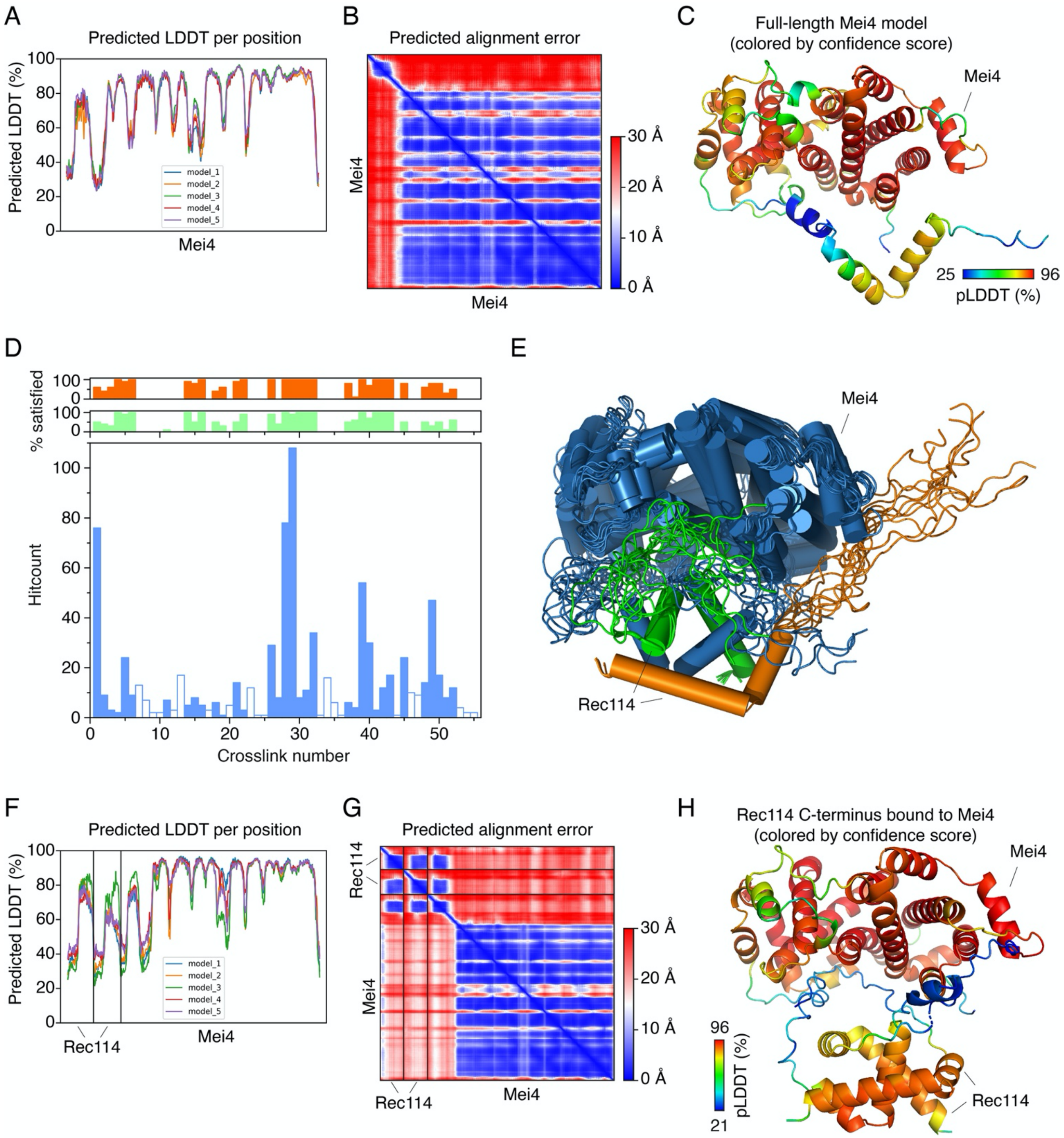

**Supplemental Figure S3: Modeling Rec114 C-terminus bound to full-length Mei4.**
(A, B, C) Quality assessment of Mei4 AlphaFold models. (A) Predicted local distance difference test (pLDDT) of the five best AlphaFold models of full-length Mei4. (B) Predicted alignment error of the highest-scoring Mei4 model (model 3). (C) AlphaFold model of full-length Mei4 (model 3) color-coded by the confidence score. (D) Violation analysis of the XL-MS restraints. The main plot shows mass spectrometry counts for the intermolecular Rec114–Mei4 crosslinks (Claeys Bouuaert et al. 2021) used as distance restraints for the structure calculation. The crosslink numbering corresponds to that in **Supplemental Table S2**. The open bars identify XL-MS restraints consistently violated in all solutions (excluded in subsequent structure refinement runs). The plots above illustrate the extent of restraint violation in the best 10 solutions for the structure calculations with all XL-MS restraints (green) or a refined set with the consistent violators excluded (orange). (E) Ten best (lowest energy) structures of the dimeric Rec114 C-terminus with full-length Mei4 aligned by the minimal Rec114–Mei4 binding core. Two chains of the Rec114 dimer are in orange and green, while Mei4 is in blue. The rmsd for the positions of Cα atoms of the Mei4 globular domain is 3.4 ± 1.7 Å. (F, G, H) Quality assessment of AlphaFold models of Rec114 C-terminus bound to full-length Mei4. (F) Predicted local distance difference test (pLDDT) of the five best AlphaFold models. (G) Predicted alignment error of the highest-scoring model (model 3). (H) AlphaFold model of the Rec114 C-terminus bound to full-length Mei4 (model 3) color-coded by the confidence score.

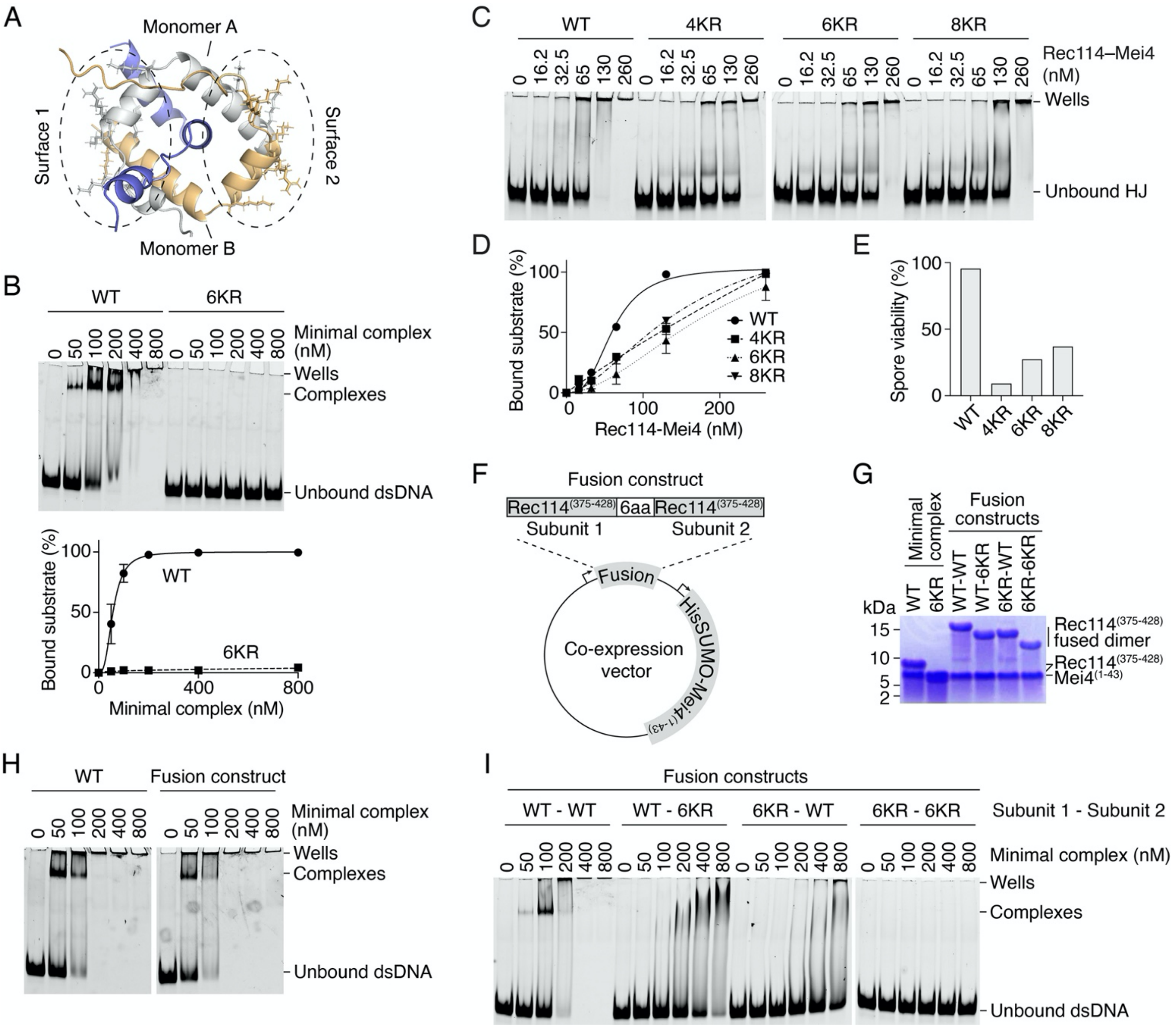

**Supplemental Figure S4: DNA-binding properties of Rec114–Mei4.**

(A) The two DNA-binding surfaces within the heterotrimeric Rec114–Mei4 complex involve residues from both Rec114 subunits (colored yellow and grey). Residues K417 and K424 of one monomer contribute to the DNA-binding surface composed of residues R395, K396, K399 and R400, K403, K407 (6KR) of the second monomer. (B) Gel-shift assay of wild type and mutant minimal trimeric Rec114–Mei4 complex. In the mutant, the 6KR residues are mutated to alanine. (C) Gel-shift assay of wild-type (WT) and mutant $^{HisFlag}$Rec114–$^{MBP}$Mei4 binding to a fluorescent HJ substrate. The 4KR mutant combines R395A, K396A, K399A and R400A (Claeys Bouuaert et al. 2021). The 8KR mutant combines the 6KR mutations with K417A and K424A. (D) Quantification of the gel-shift assay in panel C. Error bars are ranges from two independent experiments. (E) Spore viability of wild-type and mutant Rec114 strains (n = 126 tetrads/strain). We previously characterized the 4KR mutation *in vivo* and reported a spore viability of 0% (Claeys Bouuaert et al. 2021). The discrepancy is due to the presence of a myc-tag in the previous study, while in this study Rec114 is untagged. Indeed, Rec114 alleles with C-terminal myc-tags have been shown to lead to synthetic defects in other contexts (Zhang et al. 2020). (F) Design of the Rec114 fusion construct. The HisSUMO-tag at the N-terminus of Mei4 is removed during the purification. (G) SDS-PAGE analyses of purified fusion complexes used in panels H and I. (H) Gel-shift assay of wild-type minimal trimeric complexes compared to the fusion constructs (quantification presented in **Fig. 3G**). (I) Gel-shift assays of fusion complexes with a DNA-binding 6KR mutation in one or both Rec114 subunits (quantification presented in **Fig. 3H**).

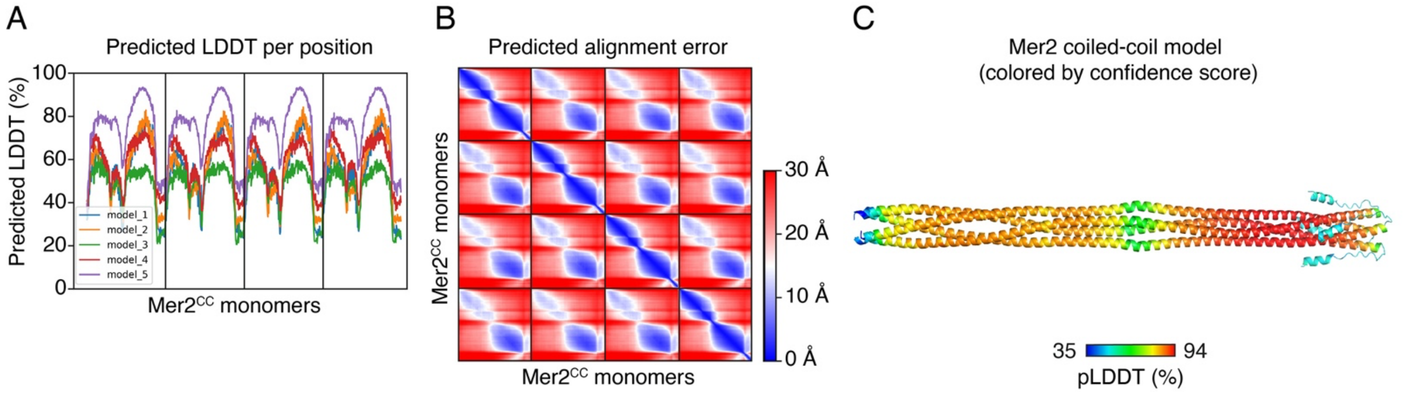

**Supplemental Figure S5: Quality assessment of Mer2 AlphaFold models.**
(A) Predicted local distance difference test (pLDDT) of the five best AlphaFold models of the Mer2 tetrameric coiled coil. All five models show parallel tetrameric structures, except for model 2. Models 3 and 4 predict that the center of the coiled coil is unfolded. (B) Predicted alignment error of the highest-scoring Mer2 tetrameric coiled coil model (model 5). (C) AlphaFold model of the Mer2 tetrameric coiled coil complex (model 5) color-coded by the confidence score.

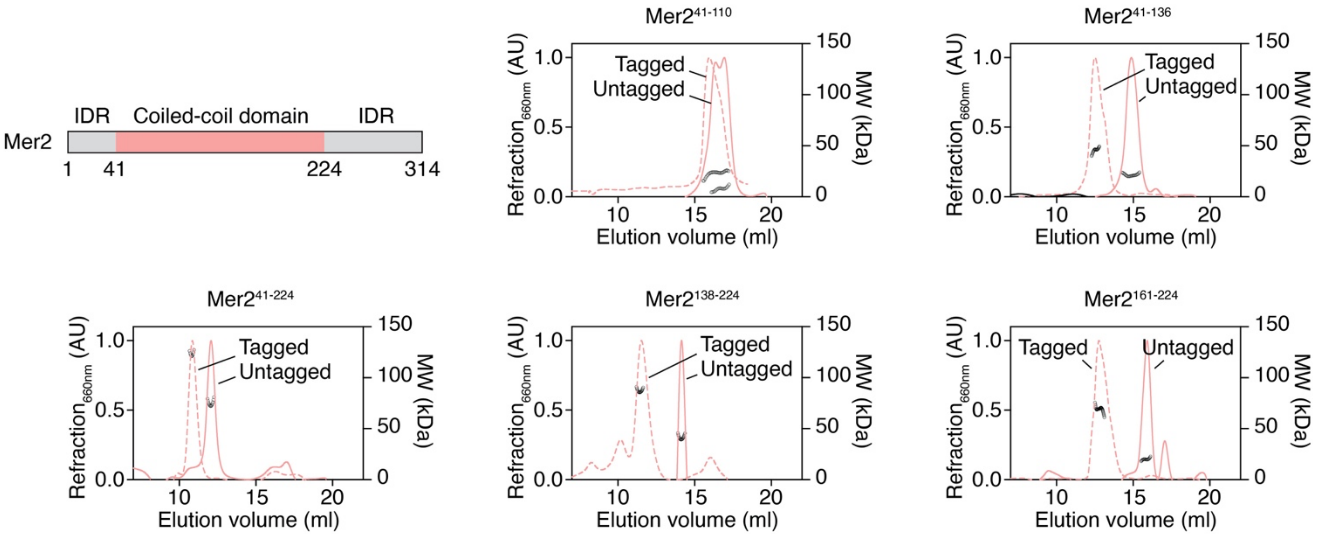

**Supplemental Figure S6: SEC-MALS analyses of untagged and HisSUMO-tagged Mer2 truncations.**
The traces show differential refraction at 660 nm (arbitrary units) and circles are molar mass measurements across the peak. See summary of the data in **Fig. 4D**.

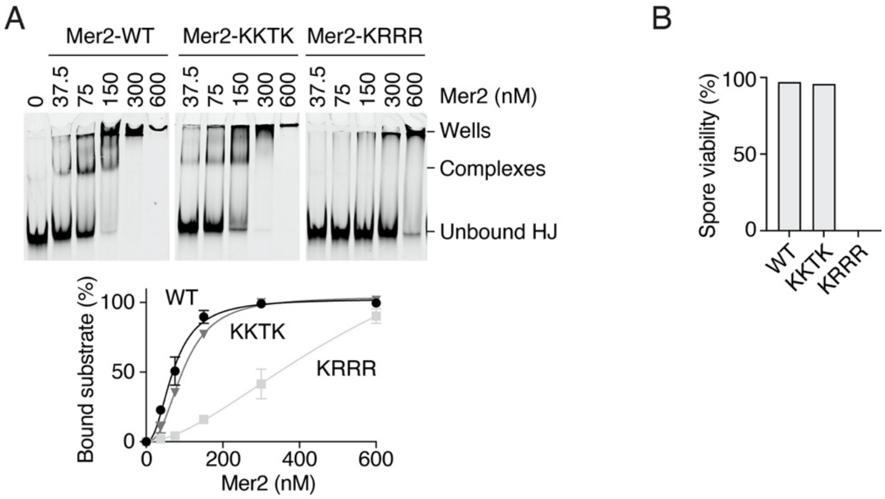

**Supplemental Figure S7: DNA-binding properties of Mer2.**
(A) Gel-shift assay of wild-type and mutant Mer2 binding to a fluorescent HJ substrate. In the mutants, the lysines and arginines of the respective KKTK and KRRR motifs are mutated to alanine. Error bars are ranges from two independent experiments. (B) Spore viability of wild-type or mutant Mer2 strains (n = 18 tetrads/strain).

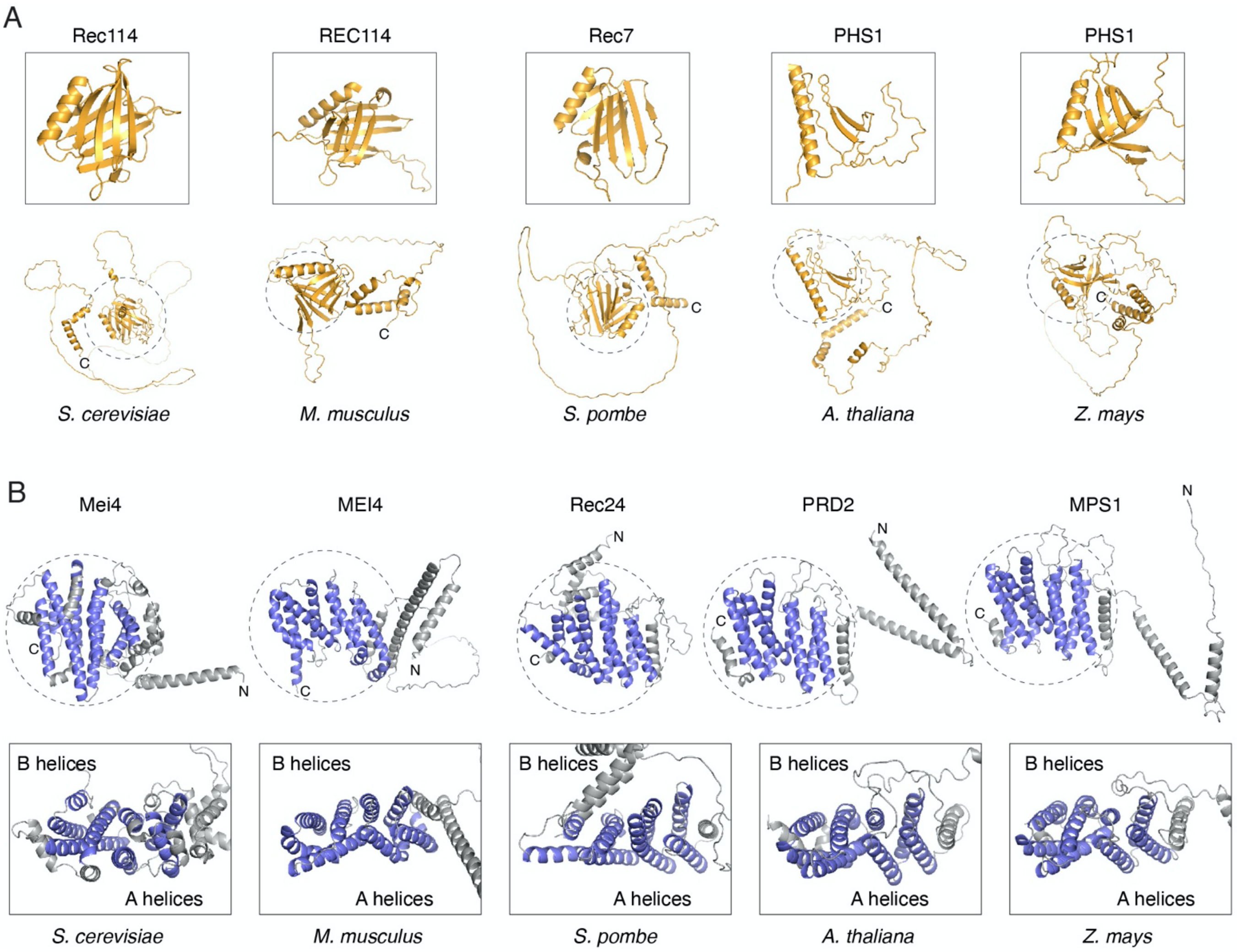

**Supplemental Figure S8: Conservation of Rec114 and Mei4 structures.**
(A) AlphaFold models of *S. cerevisiae* Rec114 (AF-A0A816AIC5), *M. musculus* REC114 (AF-Q9CWH4), *S. pombe* Rec7 (AF-P36625), *A. thaliana* PHS1 (AF-A0A178W8W3), and *Z. mays* PHS1 (AF-Q6W2J0). All Rec114 orthologs show an N-terminal PH domain (zoom), a central IDR and C-terminal helices, except *A. thaliana* PHS1 that does not display a well-folded PH domain. (B) AlphaFold models of S. cerevisiae Mei4 (AF-P29467), *M. musculus* MEI4 (AF-Q8BRM6), *S. pombe* Rec24 (AF-Q9UUJ4), *A. thaliana* PRD2 (AF-F4KDF5), and *Z. mays* MPS1 (AF-K7UF18). All Mei4 orthologs show a α-helical structure made up of four HEAT repeats (blue, see also **Supplemental Fig. S9**), flanked by flexibly-connected N-terminal helices that bind dimers of Rec114 orthologs (see **Fig. 6A**).

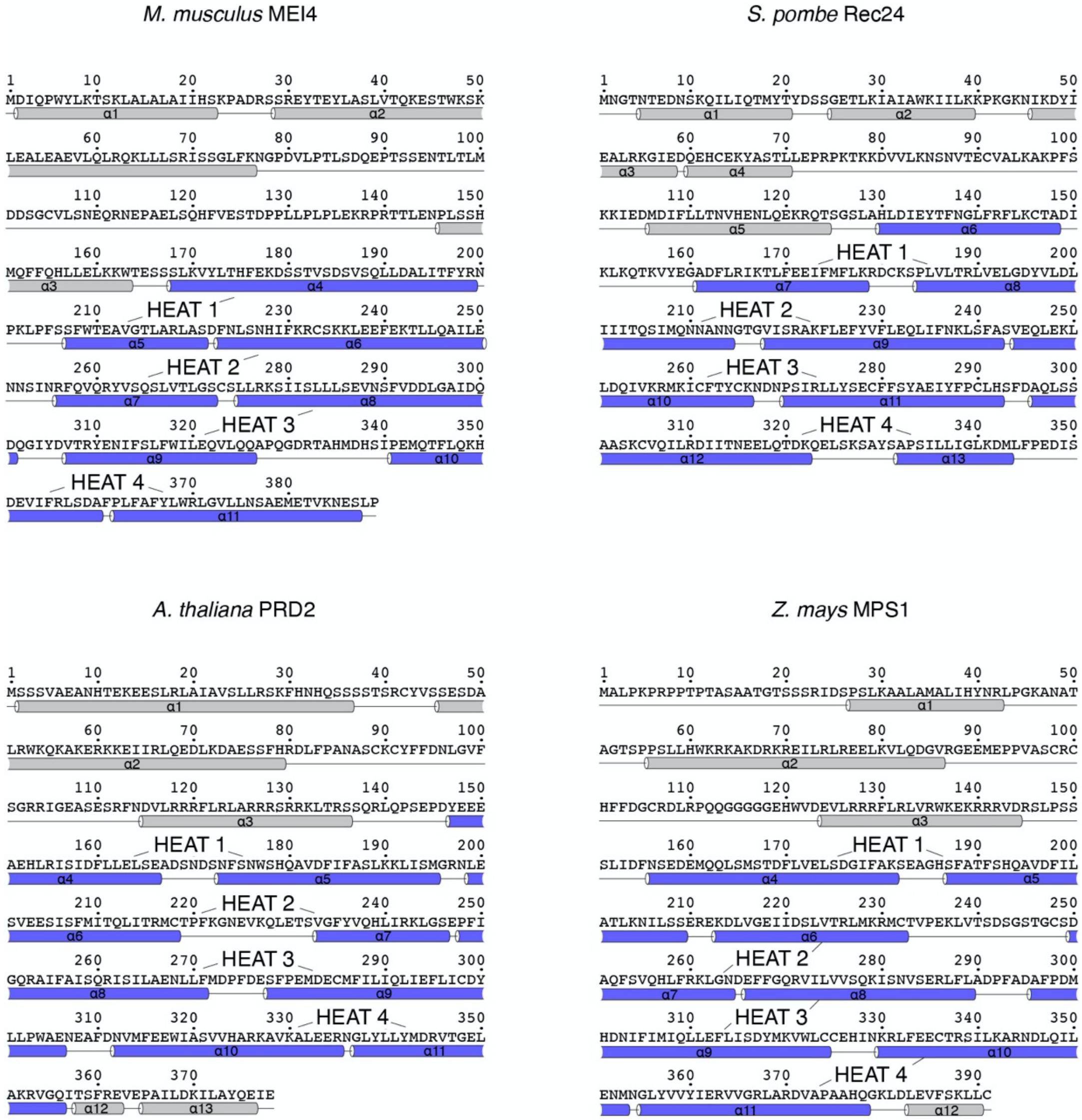

**Supplemental Figure S9: Analysis of the HEAT-repeat structure of Mei4 orthologs.**
Sequences of *M. musculus* MEI4, *S. pombe* Rec24, *A. thaliana* PRD2 and *Z. mays* MPS1 with secondary structures based on AlphaFold predictions. Helices that constitute the HEAT repeats are annotated.

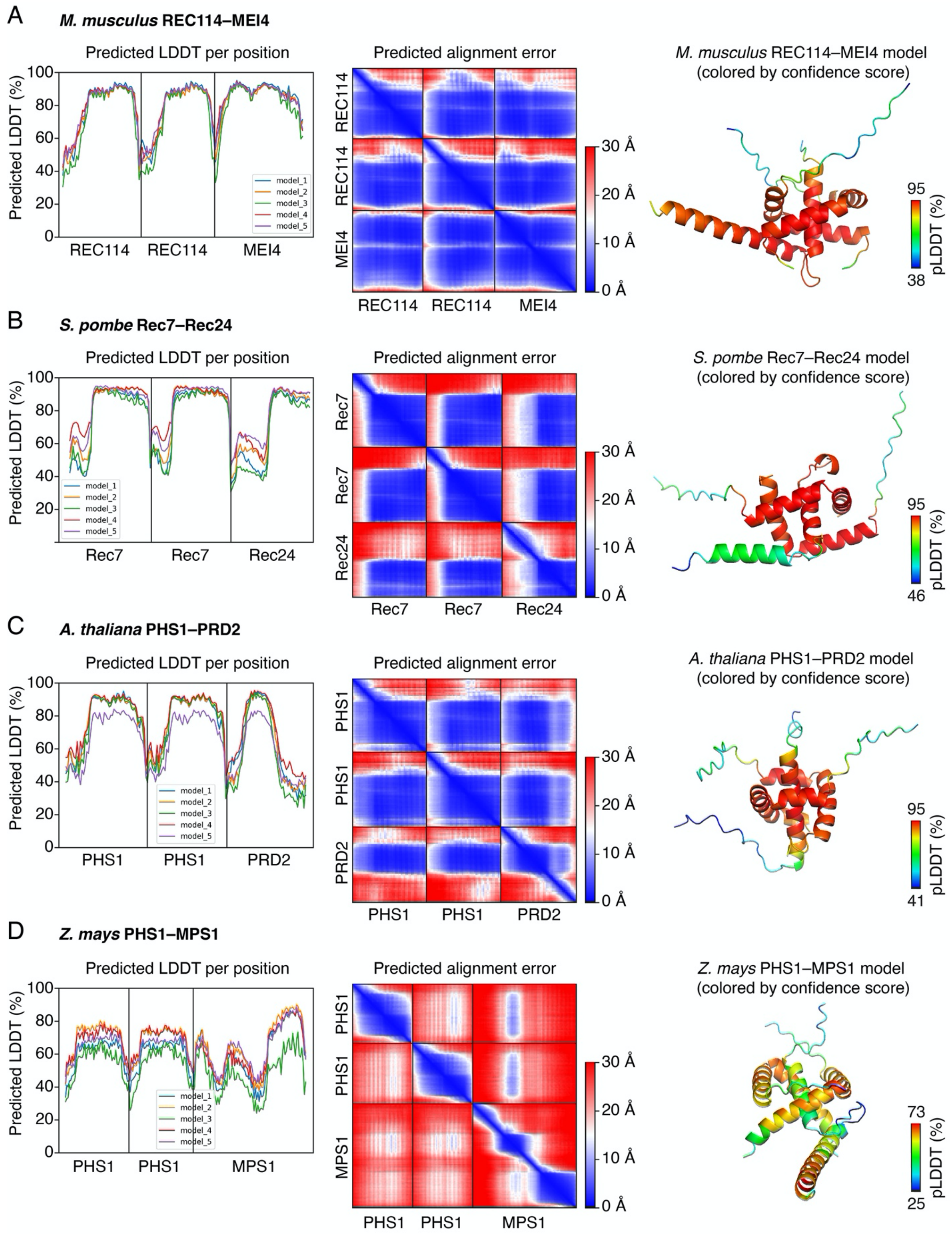

**Supplemental Figure S10: Quality assessment of AlphaFold models of Rec114–Mei4 orthologs.**
Quality assessment of structural models for the minimal trimeric complexes of Rec114–Mei4 orthologs in (A) *M. musculus*, (B) *S. pombe*, (C) *A. thaliana*, and (D) *Z. mays*. Predicted local distance difference test (pLDDT) (left), predicted alignment error (center), and structural models colored by the confidence score (right) are shown. All the models show high confidence, except for the *Z. mays* complex. For *Z. mays*, we selected model 2 based on closest similarity with other predicted structures. In all other cases, the best-scoring models are shown.

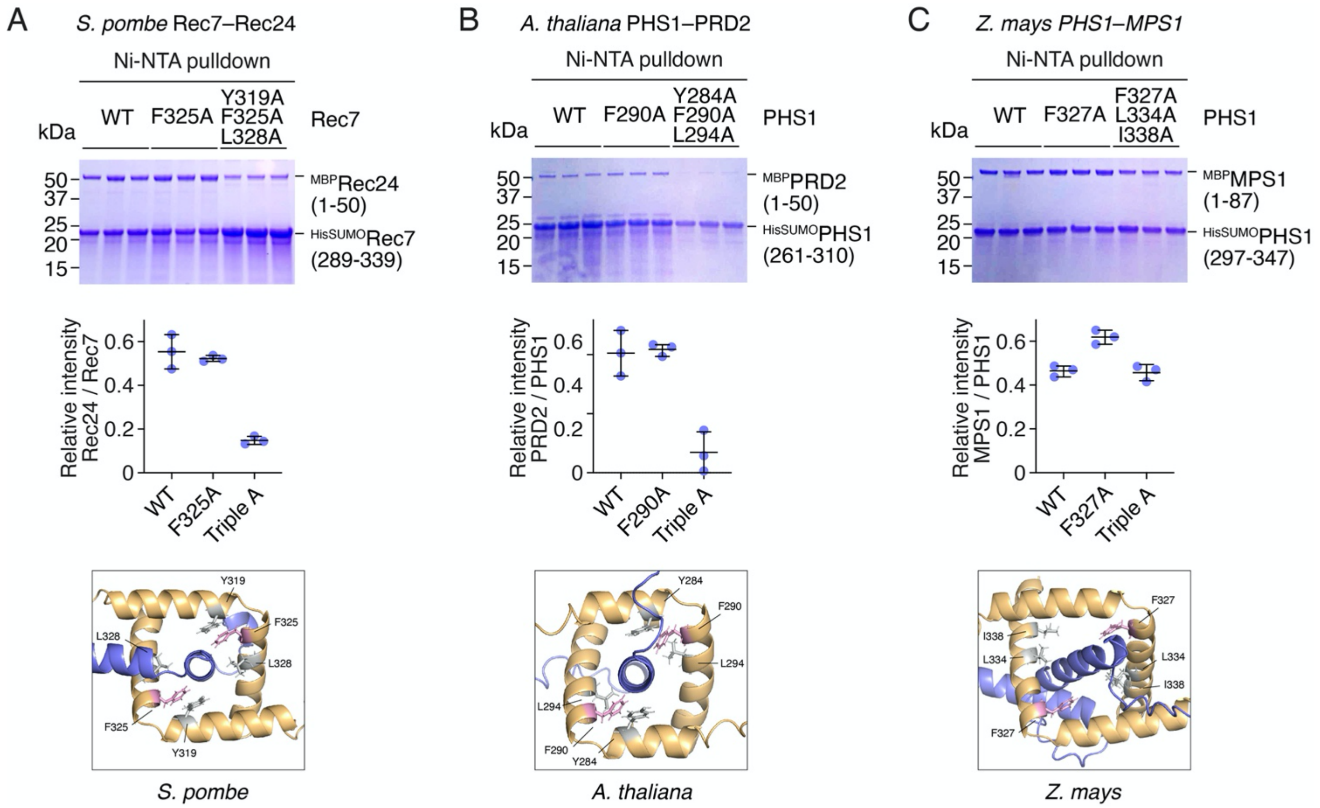

**Supplemental Figure S11: Mutational analysis of the interface of Rec114–Mei4 orthologs.**
Pulldown analyses of tagged fragments of *S. pombe* Rec7 and Rec24 (A), *A. thaliana* PHS1 and PRD2 (B), and *Z. mays* PHS1 and MPS1 (C). Rec114 orthologs are tagged with HisSUMO, Mei4 orthologs are tagged with MBP. The effect of mutating hydrophobic residues at the predicted interface between the proteins is quantified by the relative intensity of Mei4 ortholog vs. Rec114 ortholog following Ni-NTA affinity pulldown. Error bars are ranges from three replicates. The *S. pombe* Rec7-F325A mutation was previously shown to abolish the interaction with Rec24 in a yeast-two-hybrid assay (Steiner et al. 2010). Presumably, the effect on the interaction is not detectable in our biochemical assay because it is less stringent than a yeast-two-hybrid.

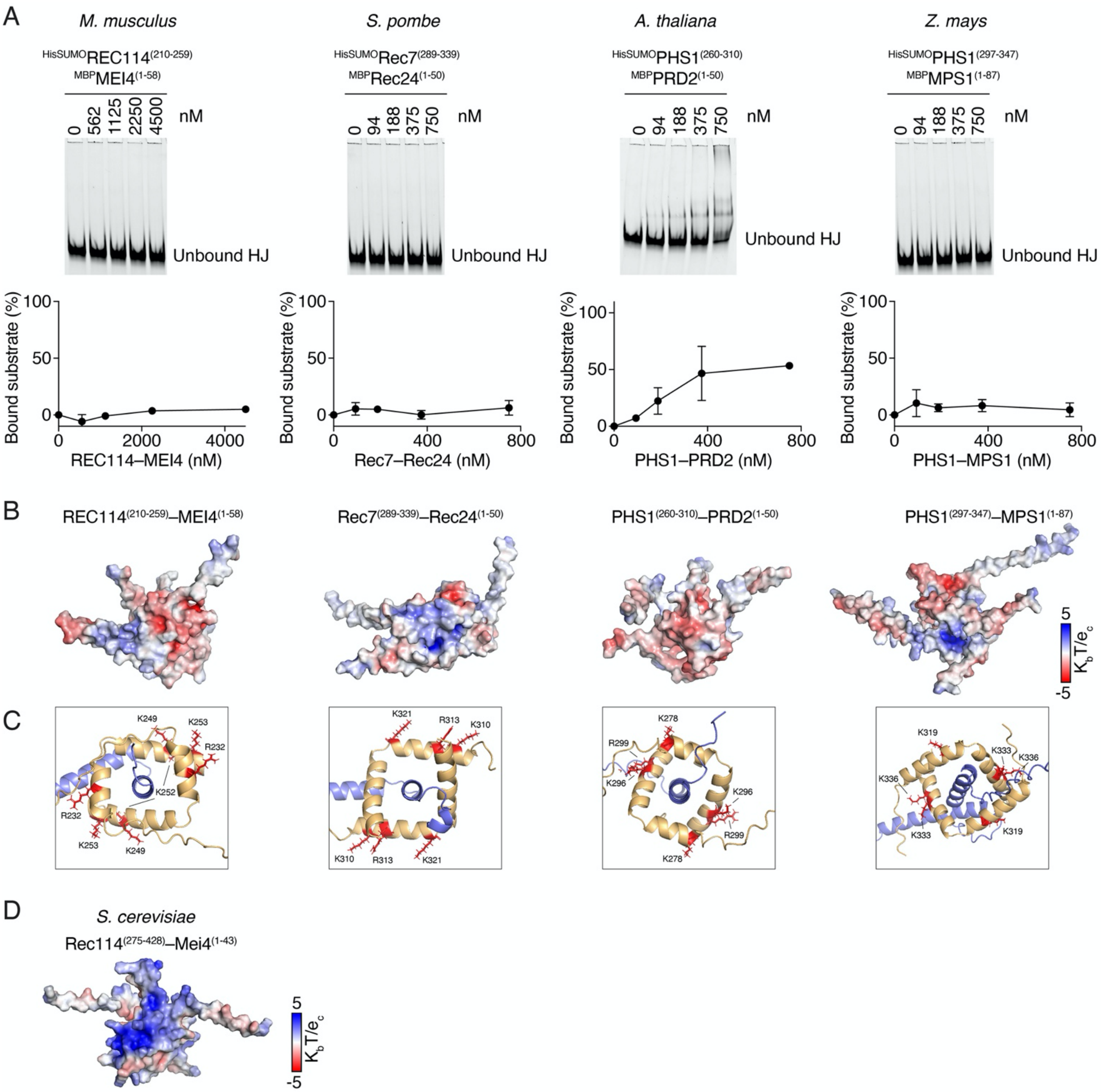

**Supplemental Figure S12: DNA-binding properties of Rec114–Mei4 orthologs.**

(A) Gel-shift analysis of tagged minimal trimeric domains of *M. musculus* REC114–MEI4, *S. pombe* Rec7–Rec24, *A. thaliana* PHS1–PRD2, *Z. mays* PHS1–MPS1 binding to a fluorescent HJ substrate. Error bars are ranges from two independent experiments. (B) Electrostatic surface potential of the minimal trimeric domain of Rec114–Mei4 orthologs. (C) Putative DNA-binding residues within the complexes. (D) Electrostatic surface potential of the minimal trimeric domain of *S. cerevisiae* Rec114–Mei4.

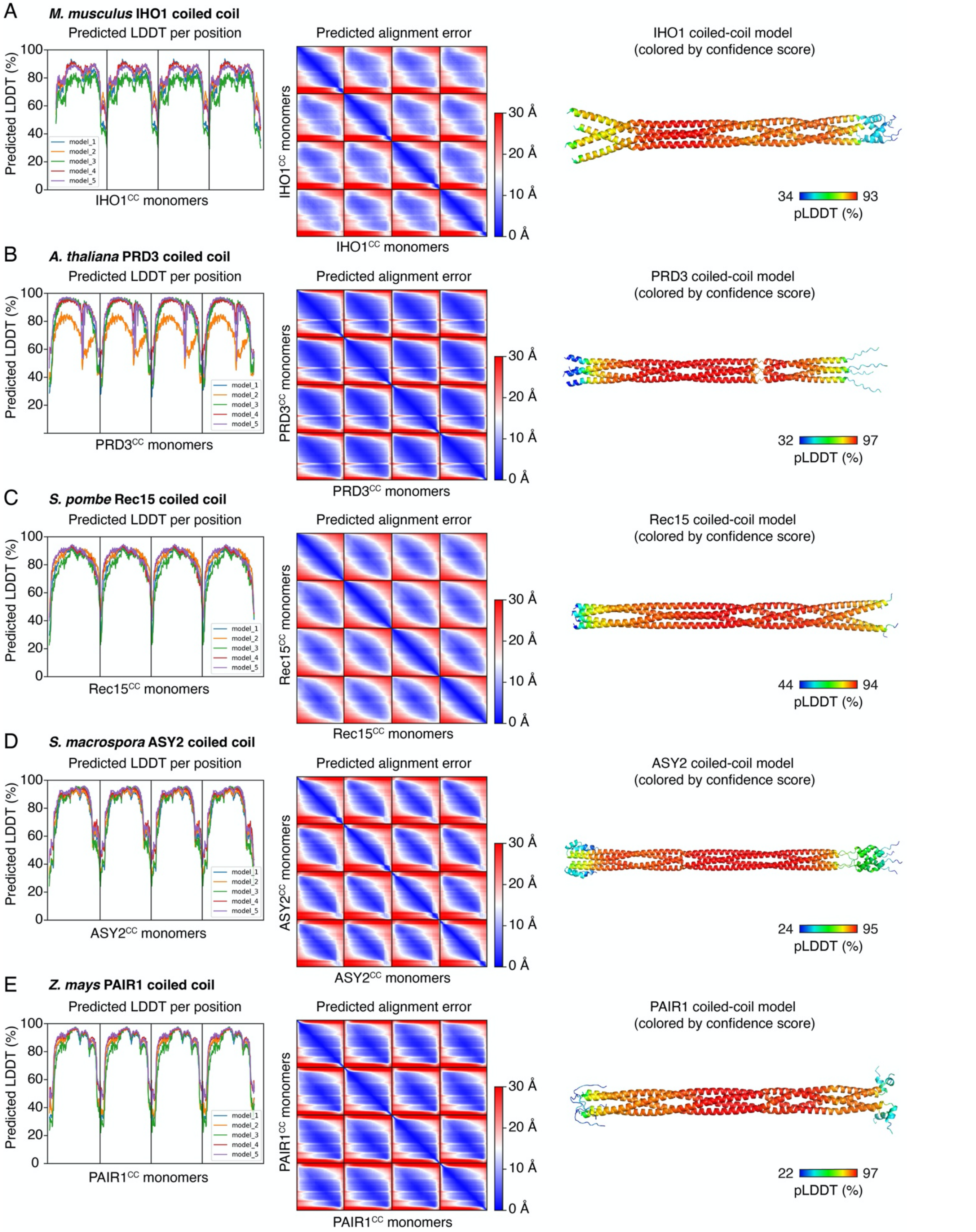

**Supplemental Figure S13: Quality assessment of AlphaFold models of Mer2 orthologs.**
Quality assessment of structural models for the tetrameric coiled-coil domain of Mer2 orthologs in (A) *M. musculus*, (B) *A. thaliana*, (C) *S. pombe*, (D) *S. macrospora*, and (E) *Z. mays*. Predicted local distance difference test (pLDDT) (left), predicted alignment error (center), and structural models colored by confidence score (right) are shown. Best-scoring models are shown. All qualitative assessments indicate higher confidence than for the yeast Mer2 model (see **Supplemental Fig. S5**).

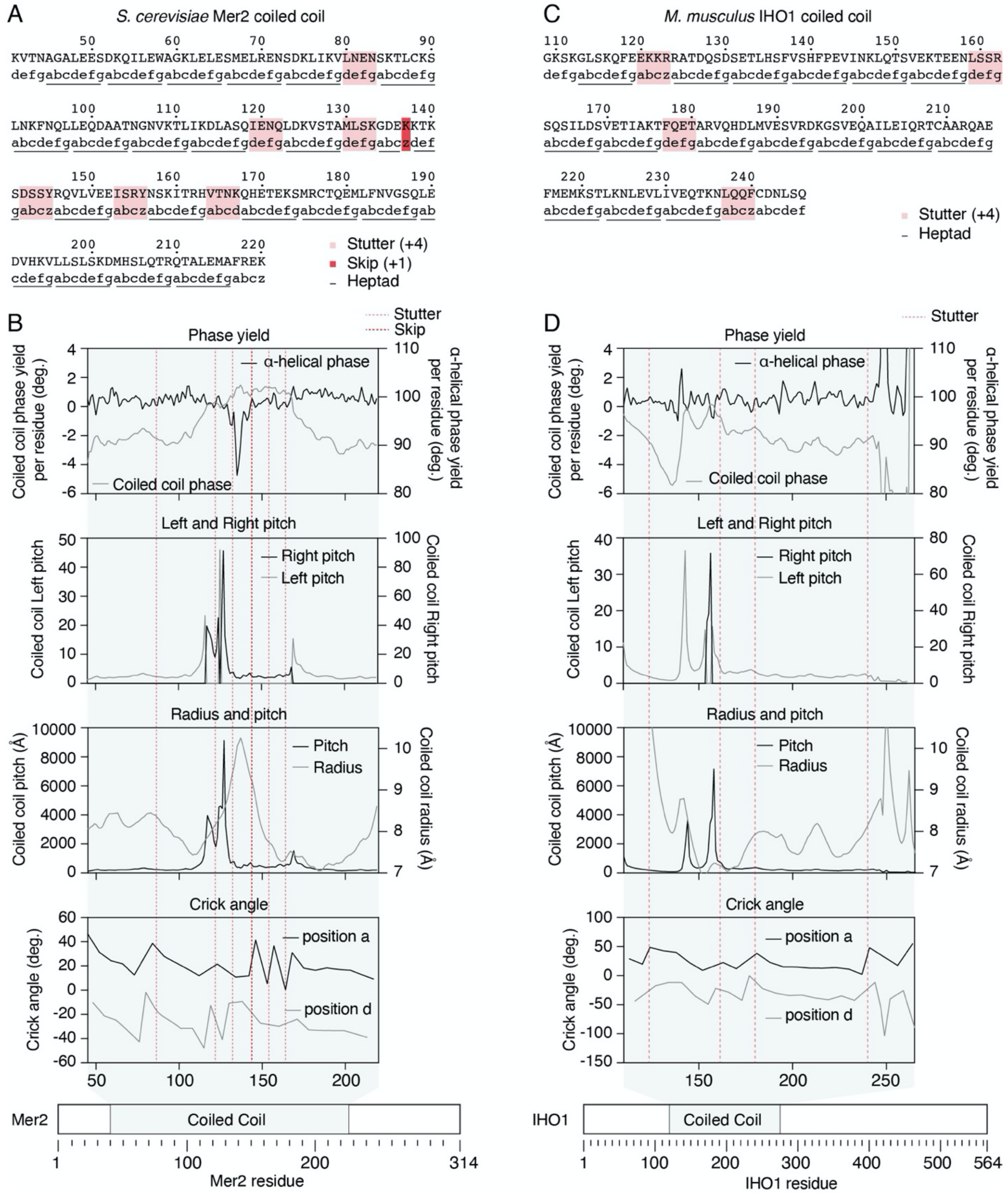

**Supplemental Figure S14: Twister analysis of predicted *S. cerevisiae* Mer2 and *M. musculus* IHO1 coiled-coil structures.**
(A) Sequence of the Mer2 coiled-coil domain, with heptad repeats (abcdef) and discontinuities (stutters and skip) indicated. Typically, positions a and d of heptad repeats are occupied by apolar residues which form the hydrophobic core of coiled coils (Parry et al. 2008). (B) Analysis of the Mer2 coiled-coil parameters using Twister (Strelkov and Burkhard 2002) (see the Materials & Methods for an explanation regarding coiled-coil parameters). The coiled-coil tends to unwind in the vicinity of the stutters, which is apparent from an increased coiled-coil phase yield and a corresponding increased coiled-coil pitch. This is accompanied by a local switch of the coiled coil from a left-handed to a right-handed geometry (the coiled-coil phase yield becomes positive). The α-helical pitch remains essentially constant, except in the vicinity of the skip, indicating that the stutters are mostly compensated by distortions of the coiled coil. In addition, the radius of the coiled coil increases at the center of the coiled coil, which likely reflects sub-optimal hydrophobic packaging of the coiled coil that results in less stable structure. (C) Sequence of the IHO1 coiled-coil domain with heptad repeats and stutters indicated. (D) Twister analysis of the predicted IHO1 coiled-coil structure also reveals local geometrical distortions of the coiled coil that coincide with the position of the stutters. The coiled coil locally switches to a right-handed geometry to accommodate a stutter around position 160.

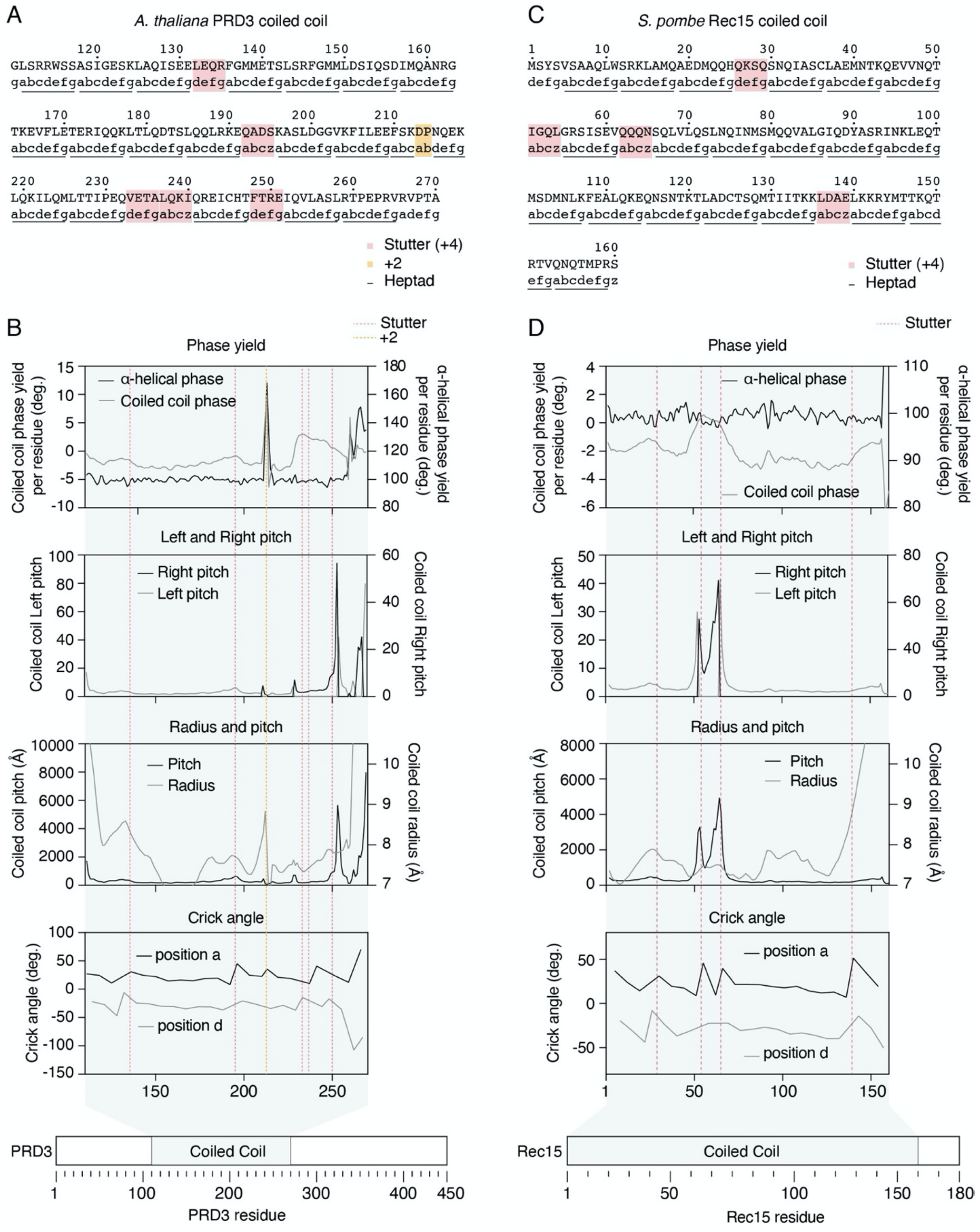

**Supplemental Figure S15: Twister analysis of predicted *A. thaliana* PDR3 and *S. pombe* Rec15 coiled-coil structures.**
(A) Sequence of the PRD3 coiled-coil domain, with heptad repeats and discontinuities based on the AlphaFold model indicated. (B) Analysis of the PRD3 coiled-coil parameters using Twister (see the Materials & Methods for an explanation regarding coiled-coil parameters). PRD3 has a +2 insertion at position 213, which is not tolerated and leads to local unfolding of the α-helices. Other stutters are compensated by a local unwinding of the coiled coil, and also lead to a switch from a left to a right-handed helix between positions ~230 and 250. (C) Sequence of the Rec15 coiled-coil domain, with heptad repeats and stutters based on the AlphaFold model indicated. (D) Twister analysis of Rec15 reveals coiled-coil discontinuities in the vicinity of the stutters, compensated by unwinding of the coiled coil and switch to a right-handed helix between residues 50 and 70.

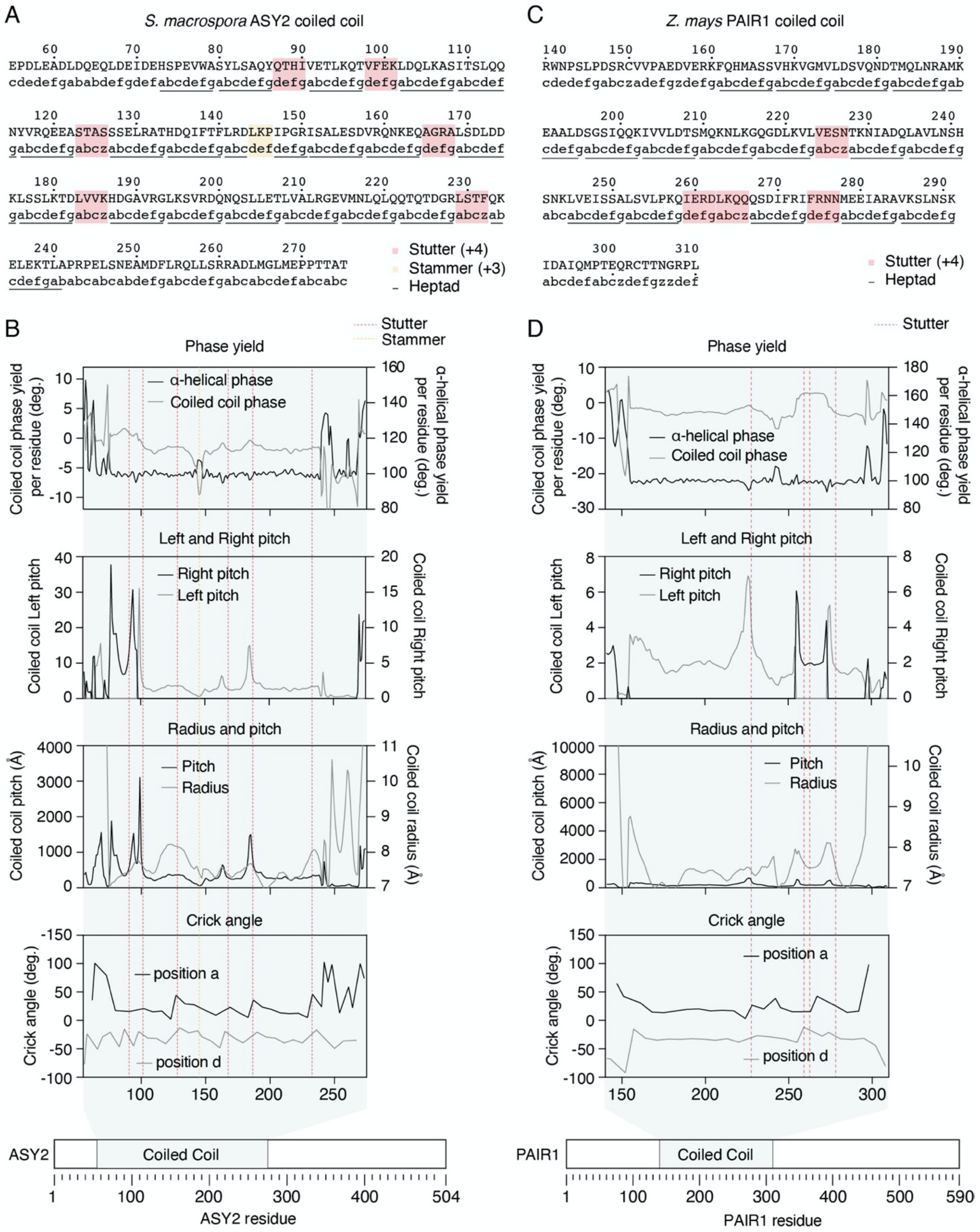

**Supplemental Figure S16: Twister analysis of predicted *S. macrospora* ASY2 and *Z. mays* PAIR1 coiled-coil structures.**
(A) Sequence of the ASY2 coiled-coil domain, with heptad repeats and discontinuities based on the AlphaFold model indicated. (B) Analysis of the ASY2 coiled-coil parameters using Twister (see the Materials and Methods for an explanation regarding coiled-coil parameters). A stammer around position 145 and a stutter around residue 230 lead to local unwinding of α-helices. Other stutters are compensated by coiled coil unwinding. (C) Sequence of the PAIR1 coiled-coil domain, with heptad repeats and stutters based on the AlphaFold model indicated. (D) Stutters are compensated by distortions of the coiled coil and local switch to a right-handed helix between residues 255 and 270.

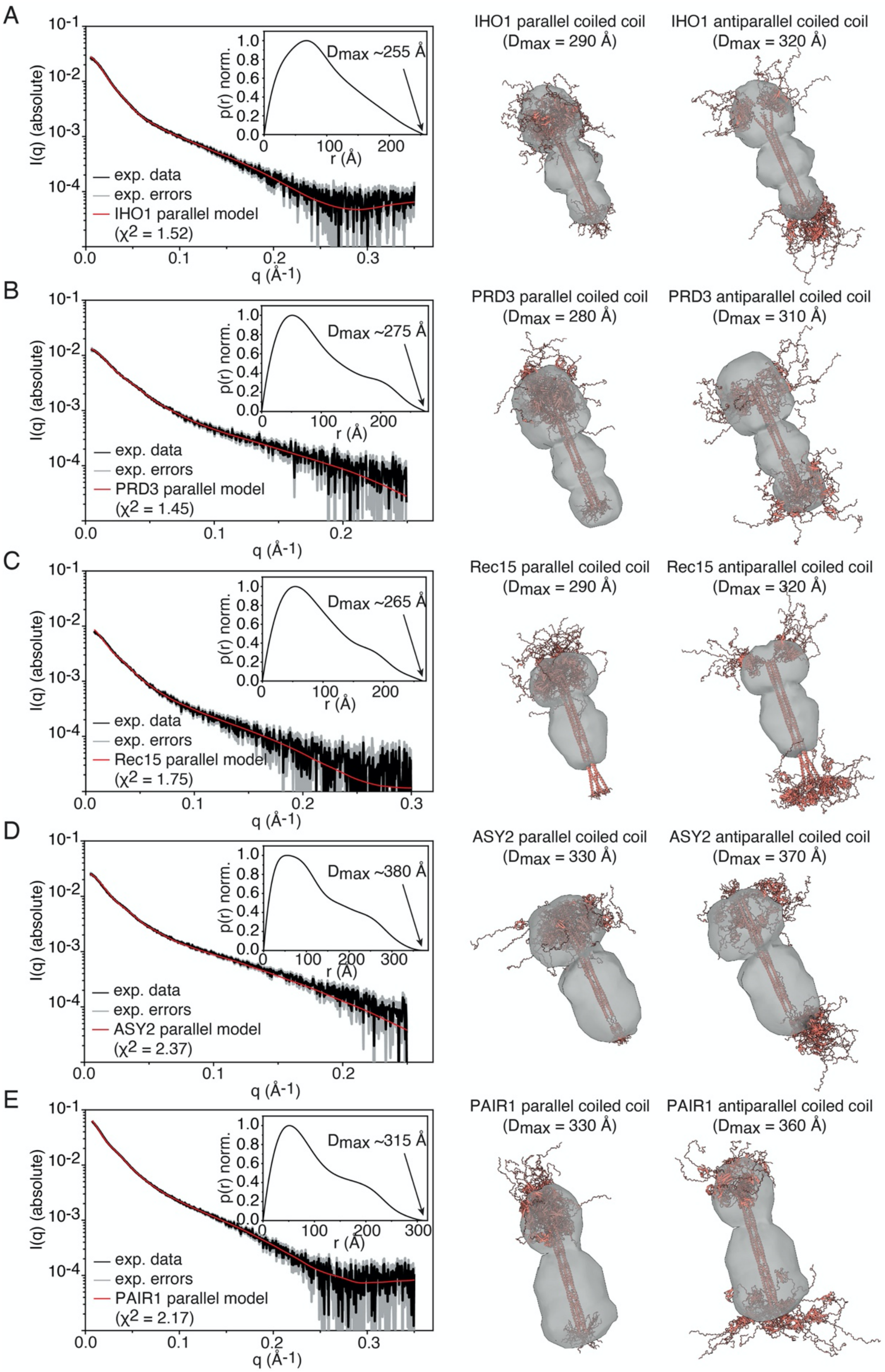

**Supplemental Figure S17: SAXS analyses of Mer2 orthologs.**
SEC-SAXS curves of HisSUMO-tagged coiled-coil domains of *M. musculus* IHO1 (A), *A. thaliana* PRD3 (B), *S. pombe* Rec15 (C), *S. macrospora* ASY2 (D), and *Z. mays* PAIR1 (E). In each panel, the graph on the left shows the experimental data (black), error margins (gray) and the fit of the parallel coiled-coil model ensemble to the data (red). The inset shows a normalized probability distance distribution function obtained based on the experimental SAXS data, with the approximate $D_{max}$ value indicated. The overlay of the different model ensembles with the *ab initio* reconstructed shape are shown at the center, with calculated $D_{max}$ values for the model ensembles indicated for convenience. In some cases (e.g., Rec15), the models stick out from the SAXS envelopes, which is probably due to fraying of the ends, resulting in flexible tails.

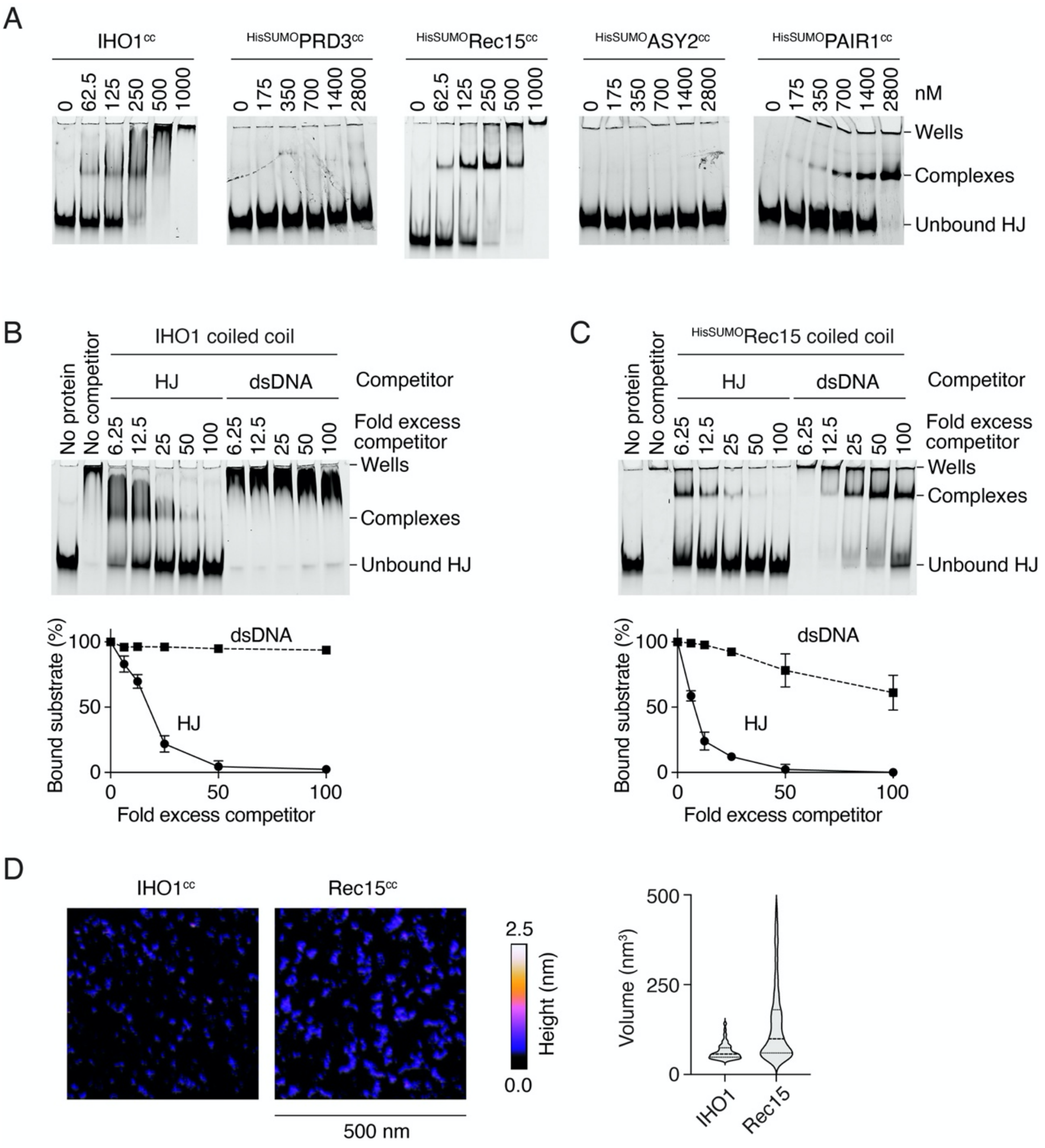

**Supplemental Figure S18: DNA-binding properties of the coiled-coil domains of Mer2 orthologs.**
(A) Gel-shift assay of the coiled-coil domains of *M. musculus* IHO1, *A. thaliana* PRD3, *S. pombe* Rec15, *S. macrospora* ASY2, and *Z. mays* PAIR1. In the presence of a fluorescent HJ substrate (10 nM). (B, C) Competition assays of the coiled-coil domain of IHO1 (600 nM) (B) Rec15 (600 nM) (C) complexes binding to a fluorescent HJ substrate (10 nM) in the presence of unlabeled dsDNA or HJ40 substrates. Error bars are ranges from two independent experiments. (D) AFM imaging of IHO1 and Rec15 coiled coil domains in the absence of DNA. The violin plot shows the volume of particles measured from two 1-μm$^2$ fields of view each. The theoretical volume of tetrameric IHO1 and Rec15 coiled-coil domains is about 115 nm$^3$.

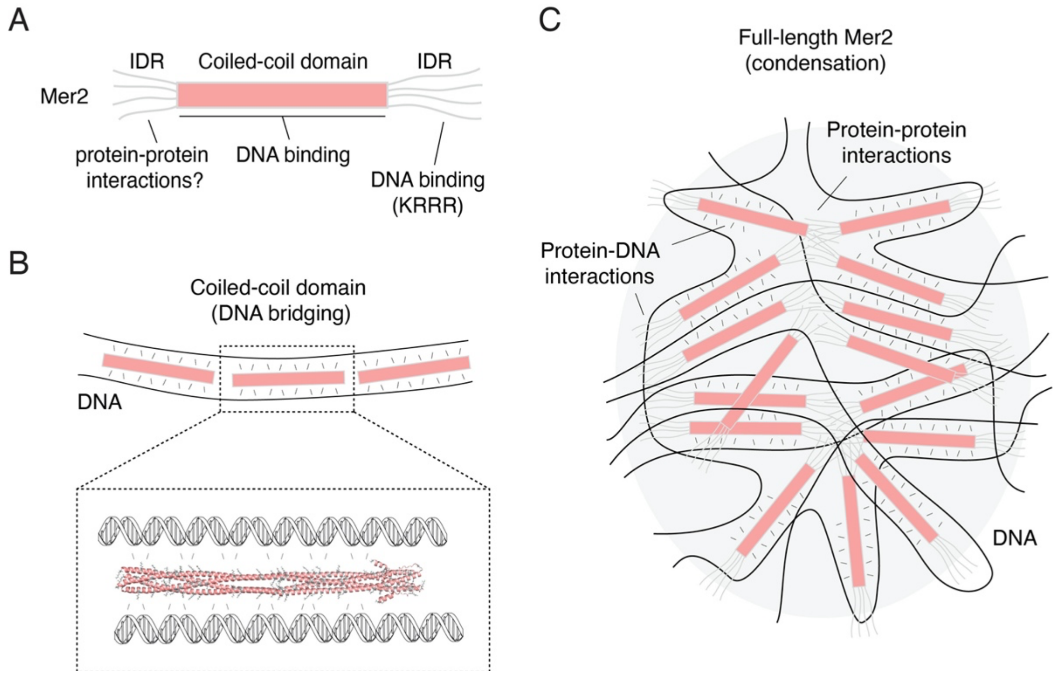

**Supplemental Figure S19: Model of condensate assembly by Mer2.**
(A) The Mer2 coiled-coil domain is flanked by N- and C-terminal intrinsically disordered regions (IDR). The coiled coil has DNA-binding/bridging activity. The C-terminal IDR is also important for DNA binding via the KRRR motif. Here, the N-terminal IDR is hypothesized to participate in low-affinity protein-protein interactions. (B) We propose that the DNA bridging activity of the coiled coil arises from multiple electrostatic interactions with lysine and arginine residues (highlighted in grey) spread along the length of the coiled coil, with the phosphoribose backbone of co-aligned DNA molecules. (C) In the context of the full-length Mer2, the combined action of protein-protein and protein-DNA interactions leads to condensation.

### SUPPLEMENTAL TABLES

| | Atom 1 | Atom 2 | Distance[a] (Å) | Class[b] |
|---|---|---|---|---|
| **Observed NOEs[c]** | | | | |
| 1 | Rec114 I397 Hδ | Rec114 V418 Hγ1 | 2.4/2.2 | A |
| 2 | Rec114 I402 Hδ | Rec114 V418 Hγ2 | 2.5/2.8 | A |
| 3 | Rec114 I402 Hγ2 | Rec114 L406 Hδ1 | 2.7/2.3 | B |
| 4 | Rec114 I402 Hγ2 | Rec114 M425 Hε | 2.9/2.7 | A |
| 5 | Rec114 L406 Hδ1 | Rec114 M425 Hε | 2.8/3.5 | A |
| 6 | Rec114 L406 Hδ2 | Rec114 L422 Hδ1 | 2.9/4.1 | A |
| 7 | Rec114 L422 Hδ1 | Mei4 A37 Hβ | 2.3 | C |
| 8 | Rec114 F426 Hδ | Mei4 A37 Hβ | 3.9 | C |
| 9 | Rec114 F426 Hε | Mei4 A37 Hβ | 3.3 | C |
| 10 | Rec114 F426 Hε | Rec114 L422 Hδ1 | 2.2/2.2 | B |
| 11 | Rec114 F426 Hδ | Rec114 L422 Hδ1 | 3.5/4.0 | B |
| 12 | Mei4 L38 Hδ2 | Mei4 W34 Hε1 | 2.6 | D |
| 13 | Mei4 L38 Hδ2 | Mei4 W34 Hδ1 | 3.5 | D |
| **Possible NOEs[d]** | | | | |
| 14 | Rec114 I397 Hγ2 | Rec114 I402 Hδ | 2.7/2.5 | B |
| 15 | Rec114 I402 Hγ2 | Rec114 L422 Hδ1 | 2.5/3.4 | A |
| 16 | Rec114 L406 Hδ1 | Rec114 L422 Hδ1 | 2.6/2.7 | B |
| 17 | Rec114 V418 Hγ2 | Rec114 L422 Hδ2 | 2.8/5.2 | B |
| 18 | Rec114 L422 Hδ2 | Mei4 A37 Hβ | 2.5 | C |

**Supplemental Table S1:** $^1$H-$^1$H NOEs detected in NOESY spectra of the minimal trimeric Rec114–Mei4 complex. [a] Closest distance between two protons in the AlphaFold structural model. When applicable, values for both Rec114 chains are provided. [b] Classification: A – interchain (Rec114); B – intrachain (Rec114); C – intermolecular (Rec114–Mei4); D – intrachain (Mei4). [c] Observed, unambiguously assigned NOEs (also see **Supplemental Fig. S2D**). [d] Possible NOEs that could not be unambiguously assigned because of the spectral overlap.

| Crosslink # | Protein 1 | Position 1 | Protein 2 | Position 2 | Hit Count | % Violated | % Satisfied |
|---|---|---|---|---|---|---|---|
| 1 | Rec114 | 387 | Mei4 | 5 | 76 | 50 | 50 |
| 2 | Rec114 | 387 | Mei4 | 35 | 9 | 100 | 0 |
| 3 | Rec114 | 387 | Mei4 | 41 | 3 | 90 | 10 |
| 4 | Rec114 | 387 | Mei4 | 52 | 2 | 0 | 100 |
| 5 | Rec114 | 387 | Mei4 | 55 | 24 | 20 | 80 |
| 6 | Rec114 | 387 | Mei4 | 58 | 9 | 20 | 80 |
| 7 | Rec114 | 387 | Mei4 | 66 | 13 | 100 | 0 |
| 8 | Rec114 | 387 | Mei4 | 102 | 7 | 100 | 0 |
| 9 | Rec114 | 387 | Mei4 | 111 | 2 | 100 | 0 |
| 10 | Rec114 | 387 | Mei4 | 134 | 2 | 100 | 0 |
| 11 | Rec114 | 387 | Mei4 | 178 | 7 | 90 | 10 |
| 12 | Rec114 | 387 | Mei4 | 201 | 3 | 100 | 0 |
| 13 | Rec114 | 387 | Mei4 | 202 | 17 | 100 | 0 |
| 14 | Rec114 | 387 | Mei4 | 229 | 5 | 30 | 70 |
| 15 | Rec114 | 387 | Mei4 | 300 | 8 | 100 | 0 |
| 16 | Rec114 | 396 | Mei4 | 5 | 5 | 10 | 90 |
| 17 | Rec114 | 396 | Mei4 | 35 | 3 | 100 | 0 |
| 18 | Rec114 | 396 | Mei4 | 42 | 6 | 50 | 50 |
| 19 | Rec114 | 396 | Mei4 | 55 | 1 | 80 | 20 |
| 20 | Rec114 | 396 | Mei4 | 58 | 5 | 100 | 0 |
| 21 | Rec114 | 399 | Mei4 | 5 | 14 | 50 | 50 |
| 22 | Rec114 | 399 | Mei4 | 41 | 3 | 20 | 80 |
| 23 | Rec114 | 399 | Mei4 | 58 | 12 | 100 | 0 |
| 24 | Rec114 | 399 | Mei4 | 66 | 1 | 100 | 0 |
| 25 | Rec114 | 399 | Mei4 | 202 | 1 | 100 | 0 |
| 26 | Rec114 | 403 | Mei4 | 5 | 29 | 70 | 30 |
| 27 | Rec114 | 403 | Mei4 | 35 | 8 | 90 | 10 |
| 28 | Rec114 | 403 | Mei4 | 41 | 78 | 10 | 90 |
| 29 | Rec114 | 403 | Mei4 | 42 | 108 | 30 | 70 |
| 30 | Rec114 | 403 | Mei4 | 52 | 8 | 20 | 80 |
| 31 | Rec114 | 403 | Mei4 | 55 | 11 | 10 | 90 |
| 32 | Rec114 | 403 | Mei4 | 58 | 34 | 0 | 100 |
| 33 | Rec114 | 403 | Mei4 | 100 | 1 | 100 | 0 |
| 34 | Rec114 | 403 | Mei4 | 102 | 16 | 100 | 0 |
| 35 | Rec114 | 403 | Mei4 | 202 | 6 | 100 | 0 |
| 36 | Rec114 | 403 | Mei4 | 251 | 1 | 100 | 0 |
| 37 | Rec114 | 407 | Mei4 | 5 | 6 | 50 | 50 |
| 38 | Rec114 | 407 | Mei4 | 35 | 4 | 90 | 10 |
| 39 | Rec114 | 407 | Mei4 | 41 | 54 | 60 | 40 |
| 40 | Rec114 | 407 | Mei4 | 42 | 30 | 50 | 50 |
| 41 | Rec114 | 407 | Mei4 | 52 | 3 | 0 | 100 |
| 42 | Rec114 | 407 | Mei4 | 55 | 12 | 0 | 100 |
| 43 | Rec114 | 407 | Mei4 | 58 | 17 | 20 | 80 |
| 44 | Rec114 | 407 | Mei4 | 315 | 1 | 100 | 0 |
| 45 | Rec114 | 417 | Mei4 | 5 | 24 | 90 | 10 |
| 46 | Rec114 | 417 | Mei4 | 58 | 10 | 100 | 0 |
| 47 | Rec114 | 424 | Mei4 | 201 | 9 | 100 | 0 |
| 48 | Rec114 | 428 | Mei4 | 35 | 14 | 100 | 0 |
| 49 | Rec114 | 428 | Mei4 | 41 | 47 | 100 | 0 |
| 50 | Rec114 | 428 | Mei4 | 42 | 17 | 70 | 30 |
| 51 | Rec114 | 428 | Mei4 | 55 | 8 | 100 | 0 |
| 52 | Rec114 | 428 | Mei4 | 58 | 12 | 50 | 50 |
| 53 | Rec114 | 428 | Mei4 | 66 | 4 | 100 | 0 |
| 54 | Rec114 | 428 | Mei4 | 201 | 2 | 100 | 0 |
| 55 | Rec114 | 428 | Mei4 | 300 | 2 | 100 | 0 |

**Supplemental Table S2:** Violation analysis of the XL-MS data of Rec114–Mei4 (Claeys Bouuaert et al. 2021) used as restraints to model the Rec114 C-terminus bound to full-length Mei4. See **Supplemental Fig. S3D.**

| Oligo | Sequence |
|---|---|
| dd015 | TCACAGAGAACAGATTGGTGGAGAAGCATCGCCAAGCG |
| dd016 | CGACGGAGCTCGAATTCGGATCACTTTTCGAACATTTTATTG |
| dd017 | CACAGAGAACAGATTGGTGGAATGAGTAGAGGCAAACTGG |
| dd018 | GACGGAGCTCGAATTCGGATTATCCTTTCTTTCTTGAAAGTG |
| dd025 | CTTTAAGAAGGAGATATACATGGGCAGCAGCCATCATCAT |
| dd026 | GGTGATGGCTGCTGCCCATGTTATCCTTTCTTTCTTGAAAG |
| dd027 | GTATAAGAAGGAGATATACAATGGAAGCATCGCCAAGCGA |
| dd028 | CCAATTGAGATCTGCCATATCACTTTTCGAACATTTTATTG |
| dd060 | AGGTATTAGTCGAAGAAATA |
| dd067 | GCCTGTAGCTGGAATCCGACGCTGTCGCCGCCTCGTCGCCTTTCGATAACA |
| dd077 | 6FAM-CTAGTATAGAGCCGGCGCGCCATGTCTAGATAGCGTTAGGTCTGCCGAATAGTACTACTCGGATCCCGAGCGAACCACGC |
| dd080 | TCAATAAAATGTTCGAAAAG |
| dd084 | TTTCAAAGCTTCCTTGATTA |
| dd085 | GACGAAGAATTCATAAAATG |
| dd095 | TGTTTTGCTTTAATTCAATC |
| dd114 | GATTATCCAGGCAACTGCCGATG |
| dd118 | GAACCGTTGCAACCTTATTAAC |
| dd121 | GGCGTGATCTGCCTTTTC |
| dd129 | CAAAACGCCCCAGCATTTG |
| dd130 | ATCCATGAGACATAAGCG |
| dd146 | AAGACAAAGTCGGAT |
| dd147 | CTCGTCGCCTTTCGATAAC |
| dd150 | CCGCTGCTGCTGCCTCCTCAAACTG |
| dd151 | CAACGGACCAATCTGACAGTGAG |
| dd154 | GTGGAAGAGGTGGAAAAGGAAC |
| dd157 | GAGCTTGCAGCGGCTTATATGACTAC |
| dd158 | AGCGTCAAGCGCTGCTGTAATAATCGTC |
| dd167 | AGCTGCTGGGGCGTTTTGATCCATG |
| dd169 | GGCGCTGTCTTGTTTTAG |
| dd170 | ATCCAATTATGTCAAAG |
| dd172 | TGCCAACTTTTCCGCGATTAGCGCCGC |
| dd173 | GACGAAGAATTCATAAAATGGGTTAATAAG |
| dd174 | GGTTCTCAATGCAATGTTCGAAAAG |
| dd175 | GTTTCAACCGCATTAACCCATTTTATG |
| dd176 | GACGAAGAATTCATAAAATGG |
| dd177 | GTTTCAACCGCATTAACCTTTTC |
| cb095 | CTAGTATAGAGCCGGCGCGCCATGTCTAGATAGCGTTAGGTCTGCCGAATAGTACTACTCGGATCCCGAGCGAACCACGC |
| cb096 | GCGTGGTTCGCTCGGGATCCGAGTAGTACTATTCGGCAGAGGATTCGAATAGGCCTAATCGAATTCCGCCATCGATGCAC |
| cb097 | GTGCATCGATGGCGGAATTCGATTAGGCCTATTCGAATCCAGACGCGAGTAGATCTTCACGGTACCCGCGGTTACCCGTG |
| cb098 | CACGGGTAACCGCGGGTACCGTGAAGATCTACTCGCGTCTCCTAACGCTATCTAGACATGGCGCGCCGGCTCTATACTAG |
| cb100 | GCGTGGTTCGCTCGGGATCCGAGTAGTACTATTCGGCAGACCTAACGCTATCTAGACATGGCGCGCCGGCTCTATACTAG |
| cb101 | GCGTGGTTCGCTCGGGATCCGAGTAGTACTATTCGGCAGAAGACGCGAGTAGATCTTCACGGTACCCGCGGTTACCCGTG |
| cb120 | GCGTGGTTCGCTCGGGATCCGAGTAGTACTATTCGGCAGA |
| cb122 | AGACGCGAGTAGATCTTCACGGTACCCGCGGTTACCCGTG |
| cb922 | CAACGTGGGCAAAGATGTCCTAGCAATGTAATCGTCTATG |
| cb923 | ATTCTACCAGTGCCAGTGATGGACATCTTTGCCCACGTTG |
| cb924 | ATCCTCTAGACAGCTCCATGATCACTGGCACTGGTAGAAT |
| cb925 | CATAGACGATTACATTGCTACATGGAGCTGTCTAGAGGAT |
| cb1186 | GATGGTCACAAGGTCCATGGCAGCCGCAGCATCCAGCTCCCCAACCCTATC |
| cb1187 | GATAGGGTTGGGGAGCTGGATGCTGCGGCTGCCATGGACCTTGTGACCATC |
| cb1315 | CAAGGACGATGATGACAAAGGTGGATCCGAAAACCTGTACTTCCAATCCAATATGTCTGAAGCGGGAAATG |
| cb1316 | CGGCCGCGACTAGTGAGCTCGTCGACTTAGTTTCTCAACCCTG |
| cb1317 | AACCTCGGGATCGAGGGAAGGGAATTCACCGAAAACCTGTACTTCCAATCCAATATGGATATTCAGCCATGG |
| cb1318 | CGACTAGTGAGCTCGTCGACCTAAGGCAACGATTCATTC |
| cb1322 | TGTCGACGGAGCTCGAATTCGGATCCTTAGAAATTATCATCGCTG |
| cb1327 | CTCACAGAGAACAGATTGGTGGATCCATGAATTTTAATGTCTG |
| cb1332 | TATTGCTGCTGACGCATTCCGATTAAC |
| cb1334 | TCTGCGGCGCTAATCAAGGAAAAGTTG |
| cb1342 | TGACTCGAGCACCACCAC |
| cb1346 | GGATCCACCAATCTG |
| cb1492 | GGTTTTCACGTTTCCGTTC |
| cb1495 | ACGCGACACGTAACTAAC |
| cb1497 | AAAGTAACGAATGCAGGC |
| cb1498 | CTCACAGAGAACAGATTGGTGGATCCGGGAAATCAAAAGGACTC |
| cb1499 | TGTCGACGGAGCTCGAATTCGGATCCTTACCCAGCCACCTGAGG |

| | |
|---|---|
| cb1503 | GCGCGCCGAGCTCGAATTTAGTTTCTCAACCCTGTAATC |
| cb1505 | GAAAACCTGTACTTCCAATCCATGGATATTCAGCCATG |
| cb1507 | CAGAGAACAGATTGGTGGACCAGAGAAGCTGACCC |
| cb1508 | GTTTCTTTACCAGACTCGAGCTAAACTTCTGCTTCTAAG |
| cb1509 | CAGAAGAATTGGGACCTGCTCTACGCTTATGTCTCATG |
| cb1510 | CATGAGACATAAGCGTAGAGCAGGTCCCAATTCTTCTG |
| cb1524 | TCAATAAAATGTTCGAAAAGGGTTCTGGAGCATCCGGAGAAGCATCGCCAAGCG |
| cb1525 | CTTTTCGAACATTTTATTGA |
| edj18 | GCCATCCAAGCATGTCAAAGTTTGGAAC |
| edj19 | GCTGTCTTGTTTTAGAGCGAATAATACCTTG |
| edj20 | CTTCAGCTCAAGATATGCTGCAG |
| edj21 | AATCTTCCATGTACTTCAGTATC |
| edj22 | GAAGAATCTTCCATGGCCTTCAGTATCTGCG |
| edj23 | AGCTCAAGATATGGCGCAGAAAGTAGAGAG |
| edj24 | AACATATCATTAGCGGATTCATCAGCCATATATG |
| edj25 | GCTCAAGCTTGATAAAGCCATTG |
| edj26 | GCTCAAGGCTGATAAAGCCGCTGATGAACTTGGTG |
| mh001 | AACGGGCGTCCTGCTTTTTCATTCTACACAGGCATATGTAACAGCAGTGTAGATCTGTTTAGCTTGCCTcGTCC |
| mh002 | CGATACTAACGCCGCCATCCAATAGGCTATGTAAATCGGCCCAATCAACAAACAAGCGTTGCTTATTGGAC |
| mh003 | TCTTTCAAGGTGGCTGAC |
| mh004 | TTTTATGCAATCTTGTCTTATC |
| mh008 | CATTTTGGTGGGTTCCTG |
| mh009 | CTACCACTTATTCGATCATGG |
| mh0036 | GTCGGATTCCAGCTACAGGC |

**Supplemental Table S3:** Oligonucleotides used in this study.

| gBlocks | Sequence |
|---|---|
| *A. thaliana* PRD3 | CACAGAGAACAGATTGGTGGATCCGGACTGTCGCGCCGCTGGTCATCCGCTAGCATCGGAGAGAGTAAGTTAGC<br>GCAGATCTCTGAAGAGTTGGAACAACGCTTCGGTATGATGGAAACGTCCTTATCTCGTTTTGGCATGATGCTGGAT<br>TCGATCCAGTCCGATATTATGCAAGCGAATCGTGGCACAAAGGAAGTATTTCTTGAAACAGAACGTATCCAACAAA<br>AACTTACCCTGCAAGATACGAGTCTGCAGCAACTGCGTAAGGAGCAAGCAGATAGTAAGGCAAGTCTTGATGGCG<br>GGGTGAAATTTATTTTAGAGGAGTTTTCCAAAGACCCAAACCAAGAAAAGTTGCAGAAAATCTTGCAAATGCTGACT<br>ACAATTCCCGAACAAGTAGAGACCGCGCTGCAGAAAATTCAGCGCGAGATCTGCCATACCTTCACACGTGAAATT<br>CAAGTTCTGGCATCGTTACGTACCCCTGAGCCACGTGTGCGCGTTCCGACTGCTTAAGGATCCGAATTCGAGCTC<br>CGTCGA |
| *S. pombe* Rec15 | CACAGAGAACAGATTGGTGGATCCATGTCGTACTCAGTAAGCGCTGCACAGCTGTGGAGTCGTAAATTAGCTATG<br>CAGGCGGAGGACATGCAGCAACATCAAAAATCTCAGAGTAATCAAATCGCTTCATGTTTAGCGGAGATGAACACC<br>AAACAAGAAGTTGTCAATCAGACAATCGGGCAGTTAGGTCGCTCTATTAGCGAGGTCCAGCAGCAGAATAGTCAA<br>CTGGTACTTCAGTCTCTTAACCAGATTAACATGAGTATGCAGCAAGTTGCCCTTGGAATCCAGGACTACGCATCAC<br>GCATCAATAAACTTGAGCAAACGATGAGTGATATGAATCTTAAATTCGAGGCGTTACAAAAGGAGCAAAACAGTAA<br>CACTAAAACATTAGCCGACTGCACAAGTCAGATGACGATTATTACAAAAAAGCTTGACGCTGAGCTTAAAAAGCGT<br>TATATGACTACAAAACAGACACGTACAGTACAAAATCAGACCATGCCTCGTTCATAAGGATCCGAATTCGAGCTCC<br>GTCGA |
| *S. macrospora* ASY2 | CACAGAGAACAGATTGGTGGATCCGAGCCGGATTTGGAGGCTGACCTGGACCAGGAGCAACTGGATGAGATCGA<br>TGAGCACTCTCCCGAGGTGTGGGCAAGTTACCTGTCGGCTCAATATCAAACGCATATTGTCGAAACATTAAAGCAG<br>ACCGTATTCGAGAAATTAGACCAACTTAAGGCTTCGATCACGAGCTTACAGCAGAACTATGTCCGTCAGGAAGAG<br>GCATCCACTGCGTCGTCTTCAGAGTTACGTGCGACTCACGACCAGATTTTTACATTTTTACGCGACCTTAAGCCTA<br>TTCCAGGTCGTATCAGCGCACTGGAGTCAGATGTTCGTCAAAATAAAGAACAGGCTGGCCGTGCCCTTAGTGACT<br>TAGACGATAAGTTGTCCTCCTTGAAGACCGATCTGGTTGTAAAACATGATGGTGCGGTACGCGGTTTAAAAAGCGT<br>GCGTGATCAAAATCAATCTCTTTTGGAAACCTTGGTTGCGTTGCGTGGAGAAGTTATGAACCTTCAGTTACAACAA<br>ACACAAACTGACGGCCGTTTGTCTACGTTTCAAAAGGAATTAGAAAAAACTTTGGCCCCCCGTCCTGAGTTATCAA<br>ACGAGGCTATGGATTTCCTTCGCCAACTTCTGAGCCGTCGCGCCGACTTGATGGGTTTGATGGAACCGCCGACCA<br>CGGCCACCTAAGGATCCGAATTCGAGCTCCGTCGA |
| *Z. mays* PAIR1 | CACAGAGAACAGATTGGTGGATCCCGTTGGAATCCGAGCTTGCCGGACTCTCGTTGTGTCGTCCCCGCTGAAGAC<br>GTGGAACGCAAGTTCCAGCATATGGCCTCCTCCGTTCATAAAGTGGGAATGGTGCTGGACAGTGTTCAAAACGAT<br>ACAATGCAGTTGAACCGCGCCATGAAGGAAGCAGCTTTAGACAGTGGAAGTATCCAACAGAAGATTGTCGTCCTT<br>GACACTTCTATGCAGAAGAATCTGAAAGGGCAGGGGGACCTGAAGGTCCTGGTAGAGTCAAATACTAAAAATATC<br>GCGGACCAGTTAGCCGTTTTGAATTCCCATTCTAATAAATTAGTCGAAATCTCCAGCGCCCTTTCTGTTCTGCCAAA<br>ACAAATCGAACGCGATTTAAAACAACAGCAGAGTGACATCTTTCGTATTTTCCGTAATAATATGGAGGAAATTGCTC<br>GTGCGGTGAAGTCACTGAATTCTAAAATCGACGCCATCCAGATGCCTACTGAACAACGTTGCACAACTAATGGGC<br>GTCCACTGTAAGGATCCGAATTCGAGCTCCGTCGA |
| *A. thaliana* PHS1 | CAGAGAACAGATTGGTGGAGACTCCACTCTAGATGCAGGCCAAACAACTGTGAAGCAGAATCCTGATCTAAAGTC<br>GCAGATACTGAAGTACATGGAAGATTCTTCATTTCAAGATATGCTGCAGAAAGTAGAGAGAATTATCGACGAAATT<br>GGAGGTAACTGGATTACCTAGATTCGAGCTCGGCGC |
| *A. thaliana* PRD2 | GAAAACCTGTACTTCCAATCCATGAGTTCAAGCGTAGCTGAAGCAAATCACACAGAGAAGGAAGAGTCGCTGAGA<br>CTGGCAATCGCTGTTTCTCTGCTACGATCTAAGTTTCAGAATCATCAATCTTCTTCTTCTACTTCCCGTTGTTATGTT<br>TCTTCTGAATCTGATGCATAGCTCGAGTCTGGTAAAGAAAC |
| *Z. mays* PHS1 | CAGAGAACAGATTGGTGGAGGAATGGATGCTGCTGAAGGAGTTGATGCCAGTATATTAACATATGACCTAATGGC<br>ACGGATAAAGACATATATGGCTGATGAATCCTTTAATGATATGTTGCTCAAGCTTGATAAAGCCATTGATGAACTTG<br>GTGGTGACATGTCGCTGTAGATTCGAGCTCGGCGC |
| *Z. mays* MPS1 | GAAAACCTGTACTTCCAATCCATGGCTCTTCCCAAACCCCGCCCGCCAACGCCGACGGCGTCAGCGGCCACGGG<br>CACCTCCTCGTCGCGAATAGACTCGCCATCGCTGAAGGCCGCGCTCGCAATGGCTCTCATCCACTACAACCGCCT<br>CCCTGGCAAAGCCAACGCCACCGCGGGCACATCACCCCCGTCTCTCCTCCACTGGAAGCGCAAGGCCAAGGACC<br>GGAAGCGCGAAATCCTCCGCCTCCGCGAGGAGCTCAAGGTCCTCCAAGATGGGGTGCGCTAGCTCGAGTCTGGT<br>AAAGAAAC |
| *S. pombe* Rec7 | CAGAGAACAGATTGGTGGATTGATTCCATCACCCCCAACGACTTCACAAATCCTACCAACAGAACTTACTGAAGAG<br>GAAAAGCAATTACGCTCCAAGGTATTATTCTATCTAAAACAAGACAGCTTCATCCAATTATGTCAAAGTTTGGAACG<br>GGTATGGAACAAGATGTAGATTCGAGCTCGGCGC |
| *S. pombe* Rec24 | GAAAACCTGTACTTCCAATCCATGAATGGCACAAATACTGAAGACAATAGTAAACAAATACTCATACAGACAATGTA<br>CACGTACGATTCGTCGGGAGAGACACTGAAAATAGCAATAGCATGGAAGATAATTTTAAAAAAACCGAAAGGAAAA<br>AATATCAAAGACTACATATAGCTCGAGTCTGGTAAAGAAAC |

**Supplemental Table S4:** Synthetic DNA fragments used in this study (gBlocks, Integrated DNA Technologies).

**Supplemental Table S4:** Oligonucleotides used in this study.

| Plasmid | Description | Reference |
|---|---|---|
| pDD003 | HisSUMO-Rec114(375-428) in pSMT3 | This study |
| pDD004 | HisSUMO-Mei4(1-43) in pSMT3 | This study |
| pDD006 | Rec114(375-428) in pETDuet1 | This study |
| pDD009 | Rec114(375-428), HisSUMO-Mei4(1-43) in pETDuet1 | This study |
| pDD015 | HisSUMO-Mer2 (K137A/K138A/K140A) in pSMT3 | This study |
| pDD044 | HisSUMO-Rec114(375-421) K405A, Mei4(1-43) in pETDuet1 | This study |
| pDD045 | HisSUMO-Rec114(375-421), Mei4(1-43) E16A, D18A in pETDuet1 | This study |
| pDD051 | HisSUMO-Rec114(375-421) K405A, E419A, Mei4(1-43) in pETDuet1 | This study |
| pDD078 | HisSUMO-Mer2 (138-224) in pSMT3 | This study |
| pDD079 | HisSUMO-Mer2 (41-136) in pSMT3 | This study |
| pDD081 | HisSUMO-IHO1(109-267) (K121A/K122A/R123A/R124A) mutant in pCCB982 | This study |
| pDD082 | mREC114-F230A/F240A/F243A mutant in pCCB987 | This study |
| pDD084 | HisSUMO-Rec7(289-339) in pETDuet1 | This study |
| pDD085 | HisSUMO-Rec7(289-339), MBP-Rec24(1-50) in pETDuet1 | This study |
| pDD086 | HisSUMO-Rec15 (K134A/K135A/K141A/K142A/R143A) mutant in pCCB991 | This study |
| pDD091 | mRec7 F325A mutant in pDD085 | This study |
| pDD092 | HisSUMO-PHS1(260-310) in pETDuet1 | This study |
| pDD093 | HisSUMO-PHS1(260-310), MBP-PRD2(1-50) in pETDuet1 | This study |
| pDD094 | HisSUMO-PHS1(297-347) in pETDuet1 | This study |
| pDD095 | HisSUMO-PHS1(297-347), MBP-MPS1(1-87) in pETDuet1 | This study |
| pDD100 | HisFlag-TEV-Rec114 (R395A/K396A/K399A/R400A/K403A/K407A) in pFastbac1 | This study |
| pDD101 | HisFlag-TEV-Rec114 (R395A/K396A/K399A/R400A/K403A/K407A/K417A/K424A) in pFastbac1 | This study |
| pDD0104 | Rec114 R395A/K396A/K399A/R400A/K403A/K407A -HphMX in Topo blunt vector | This study |
| pDD0105 | Rec114 R395A/K396A/K399A/R400A/K403A/K407A/K417A/K424A -HphMX in Topo blunt vector | This study |
| pCCB649 | SK1 Rec114 in pFastbac1 | Claeys Bouuaert et al., 2021 |
| pCCB652 | SK1 Mei4 in pFastbac1 | Claeys Bouuaert et al., 2021 |
| pCCB750 | HisSUMO-Mer2 in pSMT3 | Claeys Bouuaert et al., 2021 |
| pCCB789 | HisFlag-TEV-Rec114 in pFastbac1 | Claeys Bouuaert et al., 2021 |
| pCCB791 | MBP-TEV-Mei4 in pFastbac1 | Claeys Bouuaert et al., 2021 |
| pCCB805 | HisFlag-TEV-REC114 in pFastbac1 | This study |
| pCCB806 | MBP-TEV-MEI4 in pFastbac1 | This study |
| pCCB808 | HisSUMO-IHO1 in pSMT3 | This study |
| pCCB825 | HisSUMO-Rec114(375-428)-Mei4(1-43) in pETDuet | Claeys Bouuaert et al., 2021 |
| pCCB848 | HisFlag-TEV-Rec114 (R395A/K396A/K399A/R400A) in pFastbac1 | Claeys Bouuaert et al., 2021 |
| pCCB929 | Rec114-HphMX in Topo blunt vector | This study |
| pCCB973 | HisSUMO-Mer2 (41-314) in pSMT3 | This study |
| pCCB975 | HisSUMO-Mer2 (161-314) in pSMT3 | This study |
| pCCB978 | HisSUMO-Mer2 (1-110) in pSMT3 | This study |
| pCCB979 | HisSUMO-Mer2 (41-110) in pSMT3 | This study |
| pCCB980 | HisSUMO-Mer2 (161-224) in pSMT3 | This study |
| pCCB981 | HisSUMO-Mer2 (41-224) in pSMT3 | This study |
| pCCB982 | HisSUMO-IHO1(109-267) in pSMT3 | This study |
| pCCB984 | HisSUMO-mREC114 (210-259) and MBP-mMEI4(1-58) in pETDuet1 | This study |
| pCCB987 | mREC114-F230A mutant in pCCB984 | This study |
| pCCB988 | mREC114-F240A mutant in pCCB984 | This study |
| pCCB990 | HisSUMO-PRD3 (120-270) in pSMT3 | This study |
| pCCB991 | HisSUMO-Rec15 (1-160) in pSMT3 | This study |
| pCCB992 | HisSUMO-ASY2 (55-275) in pSMT3 | This study |
| pCCB993 | HisSUMO-PAIR1 (140-310) in pSMT3 | This study |
| pCCB1001 | Rec114 (375-428) (R395A/K396A/K399A/R400A/K403A/K407A), HisSUMO-Mei4(1-43) in pETDuet1 | This study |
| pCCB1002 | Rec114 (375-428)-6aa linker-Rec114(375-428), HisSUMO-Mei4(1-43) in pETDuet1 | This study |
| pCCB1003 | Rec114 (375-428)-6aa linker-Rec114(375-428) 6KR, HisSUMO-Mei4(1-43) in pETDuet1 | This study |
| pCCB1004 | Rec114 (375-428) 6KR-6aa linker-Rec114(375-428), HisSUMO-Mei4(1-43) in pETDuet1 | This study |
| pCCB1005 | Rec114 (375-428) 6KR-6aa linker-Rec114(375-428) 6KR, HisSUMO-Mei4(1-43) in pETDuet1 | This study |
| pEDJ10 | HisSUMO-Rec7 (289-339) (Y320A/F325A/L328A), MBP- Rec2 (41-50) in pETDuet1 | This study |
| pEDJ11 | HisSUMO-PSH1 (260-310) F290A, MBP-PRD2 (1-50) in pETDuet1 | This study |
| pEDJ12 | HisSUMO-PSH1 (297-347) (F327A), MBP-MPS1 (1-87) in pETDuet1 | This study |
| pEDJ13 | HisSUMO-PSH1 (260-310) (Y284A/F290A/L294A) MBP-PRD2 (1-50) in pETDuet1 | This study |

| | | |
|---|---|---|
| pEDJ14 | HisSUMO-PSH1 (297-347) (F327A/L334A/I338A), MBP-MPS1 (1-87) in pETDuet1 | This study |
| pMH002 | Mer2-HphMX in Topo blunt vector | This study |
| pMH026 | Mer2 (KRRR mutant)-HphMX in Topo blunt vector | This study |
| pMH029 | Rec114 (4KR)-HphMX in Topo blunt vector | This study |
| pMH030 | Mer2 (KKTK mutant)-HphMX in Topo blunt vector | This study |

**Supplemental Table S4:** Plasmids used in this study.

| Strain | Genotype | Background | Reference |
|---|---|---|---|
| CBY006 | *MATa, ho::LYS2, lys2, ura3, leu2::hisG, trp1::hisG* | SK1 | |
| CBY007 | *MATα, ho::LYS2, lys2, ura3, leu2::hisG, trp1::hisG* | SK1 | |
| CBY612 | *MATa, ho::LYS2, lys2, ura3, leu2::hisG, trp1::hisG, Mer2-KRRR::hphMX4* | SK1 | This study |
| CBY613 | *MATα, ho::LYS2, lys2, ura3, leu2::hisG, trp1::hisG, Mer2-KRRR::hphMX4* | SK1 | This study |
| CBY614 | *MATa, ho::LYS2, lys2, ura3, leu2::hisG, trp1::hisG, Mer2-KKTK::hphMX4* | SK1 | This study |
| CBY615 | *MATα, ho::LYS2, lys2, ura3, leu2::hisG, trp1::hisG, Mer2-KKTK::hphMX4* | SK1 | This study |
| CBY718 | *MATa, ho::LYS2, lys2, ura3, leu2::hisG, trp1::hisG, Rec114-6KR::hphMX4* | SK1 | This study |
| CBY719 | *MATα, ho::LYS2, lys2, ura3, leu2::hisG, trp1::hisG, Rec114-6KR::hphMX4* | SK1 | This study |
| CBY720 | *MATa, ho::LYS2, lys2, ura3, leu2::hisG, trp1::hisG, Rec114-8KR::hphMX4* | SK1 | This study |
| CBY721 | *MATα, ho::LYS2, lys2, ura3, leu2::hisG, trp1::hisG, Rec114-8KR::hphMX4* | SK1 | This study |
| CBY724 | *MATa, ho::LYS2, lys2, ura3, leu2::hisG, trp1::hisG, Rec114-4KR::hphMX4* | SK1 | This study |
| CBY725 | *MATα, ho::LYS2, lys2, ura3, leu2::hisG, trp1::hisG, Rec114-4KR::hphMX4* | SK1 | This study |

**Supplemental Table S5:** Yeast strains used in this study.

| | *S. cerevisiae* Mer2 (^HisSUMO^Mer2) | *M. musculus* IHO[1] (^HisSUMO^IHO1) | *S. pombe* Rec15 (^HisSUMO^Rec15) |
|---|---|---|---|
| **Data collection parameters** | | | |
| Beam line | SWING (SOLEIL) | SWING (SOLEIL) | SWING (SOLEIL) |
| Wavelength (Å) | 0.99 | 0.99 | 0.99 |
| *q* range ($Å^{-1}$)* | 0.009 - 0.493 | 0.003 - 0.493 | 0.003 - 0.493 |
| Concentration (mg $ml^{-1}$) (mode) | 10.5 (SEC-SAXS) | 19.4 (SEC-SAXS) | 5.5 (SEC-SAXS) |
| Buffer conditions | 25 mM HEPES-NaOH, 500 mM NaCl, 5 mM EDTA, 5% glycerol, 1 mM DTT, pH 7.5 | 25 mM HEPES-NaOH, 500 mM NaCl, 5 mM EDTA, 5% glycerol, 1 mM DTT, pH 7.5 | 25 mM HEPES-NaOH, 500 mM NaCl, 5 mM EDTA, 5% glycerol, 1 mM DTT, pH 7.5 |
| Temperature (°C) | 15 | 15 | 15 |
| **Structural parameters[§]** | | | |
| *I(0)* ($cm^{-1}$) [from Guinier] | 0.06 | 0.03 | 0.0087 |
| $R_g$ (Å) [from Guinier] | 83.28 | 72.15 | 69.37 |
| *I(0)* ($cm^{-1}$) [from *p(r)*] | 0.06 | 0.03 | 0.0090 |
| $R_g$ (Å) [from *p(r)*] | 90.41 | 76.19 | 76.70 |
| E.R. | 2.14 | 1.68 | 2.39 |
| $D_{max}$ (Å) | ~320 | ~255 | ~265 |
| Porod volume estimate, $V_p$ ($Å^3$) | 236398 | 214021 | 208227 |
| Porod exponent | 4.0 | 3.3 | 3.7 |
| **Molecular mass determination** | | | |
| MM (kDa) [from *SAXSMoW* on final merged curve] | 132.10 ($q$ = 0.45 $Å^{-1}$) | 125.20 ($q$ = 0.35 $Å^{-1}$) | 127.00 ($q$ = 0.35 $Å^{-1}$) |
| MM (kDa) [from $Q_R$ on final merged curve] | 137.29 ($q$ = 0.40 $Å^{-1}$) | 122.66 ($q$ = 0.35 $Å^{-1}$) | 129.16 ($q$ = 0.30 $Å^{-1}$) |
| MM (kDa) [from $V_p$/1.7] | 139.06 | 125.89 | 122.47 |
| Calculated MM from sequence (kDa) | 138.89 | 126.96 | 128.44 |
| ***Ab initio* modeling** | DENSS | DENSS | DENSS |
| **Software employed** | | | |
| Data processing and analysis | ATSAS, Scatter, BioXTAS RAW | ATSAS, Scatter, BioXTAS RAW | ATSAS, Scatter, BioXTAS RAW |
| Computation of theoretical intensities and fitting | XPLOR-NIH | XPLOR-NIH | XPLOR-NIH |
| **SASDB entry** | SASDSP2 | SASDSU2 | SASDST2 |

**Supplemental Table S7A:** SAXS data collection and scattering-derived parameters for Mer2, IHO1 and Rec15.
Abbreviations: *I(0)*, extrapolated scattering intensity at zero angle; $R_g$, radius of gyration calculated using either Guinier approximation (from Guinier) or the indirect Fourier transform package GNOM [from *p(r)*]; *MM*, molecular mass; $D_{max}$, maximal particle dimension; $V_p$, Porod volume; E.R., elongation ratio.
*Momentum transfer $|q| = 4\pi \sin(\theta)/\lambda$.

| | *A. thaliana* **PRD3** **(**$^{HisSUMO}$**PRD3)** | *Z. mays* **PAIR1** **(**$^{HisSUMO}$**PAIR1)** | *S. macrospora* **ASY2** **(**$^{HisSUMO}$**ASY2)** |
|---|---|---|---|
| **Data collection parameters** | | | |
| Beam line | SWING (SOLEIL) | SWING (SOLEIL) | SWING (SOLEIL) |
| Wavelength (Å) | 0.99 | 0.99 | 0.99 |
| *q* range (Å$^{-1}$)* | | | |
| Concentration (mg ml$^{-1}$) (mode) | 9.0 (SEC-SAXS) | 13.4 (SEC-SAXS) | 6.9 (SEC-SAXS) |
| Buffer conditions | 25 mM HEPES-NaOH, 500 mM NaCl, 5 mM EDTA, 5% glycerol, 1 mM DTT, pH 7.5 | 25 mM HEPES-NaOH, 500 mM NaCl, 5 mM EDTA, 5% glycerol, 1 mM DTT, pH 7.5 | 25 mM HEPES-NaOH, 500 mM NaCl, 5 mM EDTA, 5% glycerol, 1 mM DTT, pH 7.5 |
| Temperature (°C) | 15 | 15 | 15 |
| **Structural parameters§** | | | |
| *I(0)* (cm$^{-1}$) [from Guinier] | 0.013 | 0.068 | 0.027 |
| $R_g$ (Å) [from Guinier] | 77.76 | 81.40 | 95.95 |
| *I(0)* (cm$^{-1}$) [from *p(r)*] | 0.013 | 0.070 | 0.028 |
| $R_g$ (Å) [from *p(r)*] | 81.95 | 88.00 | 104.40 |
| E.R. | 2.67 | 3.00 | 3.38 |
| $D_{max}$ (Å) | ~275 | ~315 | ~380 |
| Porod volume estimate, $V_p$ (Å$^3$) | 222290 | 218003 | 222484 |
| Porod exponent | 3.4 | 3.8 | 3.6 |
| **Molecular mass determination** | | | |
| MM (kDa) [from *SAXSMoW* on final merged curve] | 135.60 (*q* = 0.25 Å$^{-1}$) | 129.8 (*q* = 0.45 Å$^{-1}$) | 152.70 (*q* = 0.30 Å$^{-1}$) |
| MM (kDa) [from $Q_R$ on final merged curve] | 121.04 (*q* = 0.25 Å$^{-1}$) | 130.79 (*q* = 0.35 Å$^{-1}$) | 154.35 (*q* = 0.20 Å$^{-1}$) |
| MM (kDa) [from $V_p$/1.7] | 130.76 | 128.24 | 130.87 |
| Calculated MM from sequence (kDa) | 128.78 | 131.80 | 155.04 |
| ***Ab initio* modeling** | DENSS | DENSS | DENSS |
| **Software employed** | | | |
| Data processing and analysis | ATSAS, Scatter, BioXTAS RAW | ATSAS, Scatter, BioXTAS RAW | ATSAS, Scatter, BioXTAS RAW |
| Computation of theoretical intensities and fitting | XPLOR-NIH | XPLOR-NIH | XPLOR-NIH |
| **SASDB entry** | SASDSS2 | SASDSR2 | SASDSQ2 |

**Supplemental Table S7B:** SAXS data collection and scattering-derived parameters for PRD3, PAIR1 and ASY2.
Abbreviations: *I(0)*, extrapolated scattering intensity at zero angle; $R_g$, radius of gyration calculated using either Guinier approximation (from Guinier) or the indirect Fourier transform package GNOM [from *p(r)*]; *MM*, molecular mass; $D_{max}$, maximal particle dimension; $V_p$, Porod volume; E.R., elongation ratio.
*Momentum transfer $|q| = 4\pi\sin(\theta)/\lambda$.
